# Response diversity in the context of multifarious environmental change

**DOI:** 10.1101/2023.12.13.571413

**Authors:** Francesco Polazzo, Romana Limberger, Frank Pennekamp, Samuel R. P.-J. Ross, Gavin L. Simpson, Owen L. Petchey

**Author notes:** **Email:** (FP); (RL); (FP); (SRP-JR); (GLS); (OLP). **Corresponding author:** Owen L. Petchey. **Contact author:** Francesco Polazzo. **Open Research statement:** Code and data to reproduce the analysis can be found on Github: https://github.com/opetchey/multifarious_response_diversity/releases/tag/v0.1.

## Abstract

Response diversity represents the inter- and intraspecific trait variation in organismal responses to the environment. Assemblages composed of organisms displaying large variation in their response to the environment (that is, having high response diversity) are expected to have higher temporal stability of aggregate community and ecosystem properties such as ecosystem functioning (i.e., an insurance effect). Yet, response diversity is not commonly measured in empirical studies, and when it is measured, this is done in different ways. Moreover, most proposed measures of response diversity concern situations with only one driver of environmental change. Thus far, no specific approach exists to measure response diversity in the context of multiple simultaneously changing (multifarious) environmental drivers. Here, we propose a new method to empirically quantify response diversity in the context of multifarious environmental change. First, we illustrate this method using simulated data. Next, we reveal the role of the direction of environmental change in shaping response diversity when multiple drivers of environmental change fluctuate over time. We show that, when the direction of the environmental change is unknown (that is, there is no information or *a priori* expectation about how an environmental condition has changed or will change in future), we can quantify the *potential* response diversity for a given community under any possible future environmental change scenario. That is, we can estimate the potential response capacity of a system under a range of extreme or realistic environmental changes, capturing its complete insurance capacity, with utility for predicting future responses to even multifarious environmental change. Finally, we investigate the drivers of response diversity in a multifarious environmental change context, showing how response diversity depends on: 1) the diversity of species responses to each environmental variable considered, 2) the relative effect of each environmental variable on species’ performance, 3) the correlation between the diversity in species’ responses to different environmental variables, and 4) the mean temporal value of the environmental variable. In doing so, we take an important step towards understanding the insurance capacity of ecological communities exposed to multifarious environmental change.

## 2. Introduction

Ecological stability concerns responses of ecosystems and their components to environmental change (Donohue *et al*. 2013; Mccann 2000; Pimm 1984). A stable population, community, or ecosystem will change less than a less stable or unstable one or will recover more rapidly after disturbance. Greater stability can mean lower chance of population extinction (Fussmann *et al*. 2014) or lower variability of an aggregate property such as total community biomass or ecosystem processes or functions (Hautier *et al*. 2015; Pennekamp *et al*. 2018). Mechanistic understanding of ecological stability is key to ecological sustainability; that is, the use of natural resources to withstand production for finite time without environmental deterioration, and ideally without losing native biodiversity (Aarts 1999; Walker *et al*. 2023). Accordingly, the study of stability has occupied a central place in ecological research for decades, and has emerged as a multidimensional concept, including components such as resistance, resilience, persistence, robustness, and temporal or spatial variability (Donohue *et al*. 2013; Pimm 1984). The temporal variability (or temporal stability) of aggregate properties has been a central dimension of stability in theoretical and empirical work for decades (Hautier *et al*. 2015; Tilman & Downing 1994; Yachi & Loreau 1999). Low temporal variability (that is, high stability) is associated with predictable and sustainable delivery of ecosystem functions (Elmqvist *et al*. 2003; Renard & Tilman 2019). Here, we will focus primarily on this critical dimension of ecological stability.

There is a consensus that biodiversity dampens variability in aggregate community and ecosystem properties (Hautier *et al*. 2015; Isbell *et al*. 2015; Tilman *et al*. 2014; Yachi & Loreau 1999). This is because, in species rich communities, there is a higher chance that a reduction in abundance or performance (e.g., intrinsic growth rate) of one species may be compensated for by an increase in abundance or performance of another species (Yachi & Loreau 1999). Such dynamics have been named compensatory or asynchronous and have been identified as a key driver of temporal stability across a range of systems (Craven *et al*. 2018; Gonzalez & Loreau 2009; Sasaki *et al*. 2019; White *et al*. 2023).

In ecological communities with a greater number of species, the likelihood of compensatory dynamics emerging by chance increases (Yachi and Loreau 1999). Compensatory dynamics are more likely to occur if species differ in their responses to environmental conditions (McCann 2016); species with identical environmental tolerances are unlikely to exhibit compensatory dynamics (Mori *et al*. 2013). When environmental conditions change over time, as is common in natural systems, communities composed of species that differ in their environmental responses are more likely to maintain stable levels of aggregate biomass or functioning through time, owing to asynchronous (compensatory) fluctuations (Mori *et al*. 2013), if that they perform the same function. This diversity of responses to environmental change is often termed response diversity (Elmqvist *et al*. 2003; Mori *et al*. 2013; Nyström 2006).

The stabilising effect of response diversity has good conceptual foundations. For example, Ives et al. (1999) showed that differences in species responses to environmental change (response diversity) may determine community stability by dampening total abundance variability over time. Using a modelling approach of randomly constructed competitive communities, Ives and Carpenter (2007) found that response diversity is the most significant driver of temporal stability, outweighing the destabilising effects of interspecific interactions. These findings suggested that it is not biodiversity *per se* that enhances stability, but rather the diversity of species responses to the environment. That is, response diversity may be an ecological mechanism (although likely not the only one) underlying the insurance effect of biodiversity (Yachi & Loreau 1999), and promoting asynchronous population dynamics in changing environments (Downing *et al*. 2008).

A limited number of empirical studies have quantified response diversity explicitly. Critically, the few empirical studies of response diversity that do exist, have quantified response diversity in diverse ways (Ross *et al*. 2023). For example, the majority of empirical studies of response diversity use functional traits that approximate some feature of species response to the environment, such as clutch size or specific leaf area (Hordley *et al*. 2021; Sasaki *et al*. 2019; White *et al*. 2023). Another approach measures a species-specific interaction between environment and abundance in a community context; if species show different abundance–environment relationships, there is response diversity in a binary sense (Bartomeus *et al*. 2013; Cariveau *et al*. 2013; Winfree & Kremen 2008). Alternatively, response diversity can be quantified by measuring some aspect of a population’s performance, such as its intrinsic rate of increase or biomass change in response to the environment (McCann 2016; Ross *et al*. 2023). The range of performance–environment model slopes then acts as a direct estimate of response diversity (Leary & Petchey 2009). Recently, Gladstone-Gallagher *et al*. (2023) also measured response diversity using ecological network analysis of traits, where a higher network complexity represents higher response diversity.

Despite empirical studies using these methods to measure response diversity, the lack of a common, standardised method to quantify response diversity has hindered comparisons of response diversity between studies, limiting the opportunity to systematically investigate whether response diversity is a fundamental mechanism dampening temporal variability across diverse communities. As such, empirical studies of response diversity require a standard and operationalizable method for measuring this understudied element of biodiversity.

Recently, Ross *et al*. (2023) developed a robust and flexible method to study how species identity and diversity shape ecological stability in the face of environmental change. The method is based on **species performance curves** (such as how temperature affects a species’ growth rate, see Glossary) and can account for nonlinear performance-environment relationships. Response diversity is then measured as the variation in the slope of the individual species’ performance curves. By capturing how species performances depend on environmental context, this new response diversity framework allows testing the diversity-stability relationship from a new perspective and represents a novel tool to mechanistically understand this relationship in a standardised way (Ross *et al*. 2023).

Nevertheless, this framework has a significant limitation in that it cannot capture response diversity in the context of multiple simultaneously changing (multifarious) **environmental drivers**. This is particularly limiting as human activities have caused an increase in the number and intensity of anthropogenic drivers acting on ecosystems (Bowler *et al*. 2020; IPCC-IPBES 2020). Ecosystems are now exposed to multiple environmental drivers simultaneously, which can undermine their biodiversity as well as their stability to a larger extent than when exposed only to a single environmental driver (Zelnik *et al*. 2018). Moreover, a multifarious context raises the potential for interactions between drivers, which may result in synergistic effects larger than, or antagonistic effects smaller than those expected from the sum of the individual stressors (Piggott *et al*. 2015). Multiple interacting drivers impact both biodiversity and stability (Pires *et al*. 2018; Polazzo & Rico 2021), and meta-analyses summarising available research on multiple driver impacts have highlighted that non-additive effects are common (Birk *et al*. 2020; Crain *et al*. 2008; Jackson *et al*. 2016). New methods which are better able to capture the response diversity of ecological communities in the context of multifarious environmental change are therefore urgently needed.

To date, there has been little, if any, exploration of response diversity in the context of multifarious environmental change. This is despite differences in environmental tolerances among species resulting in low redundancy across environmental contexts, suggesting that response diversity is key to stability even under multiple simultaneous environmental changes (Mori *et al*. 2013). Hence, an extension of empirical response diversity methods is needed to empirically test whether response diversity is a key determinant of stability in communities exposed to multifarious environmental change. Understanding the role of response diversity in driving the diversity-stability relationship in face of multiple environmental drivers may help the translation of experimental results and theoretical advances into information for policymakers, due to the real-world relevance of multifarious environmental change. Such information is important because the temporal variability of aggregate properties plays a central role in economic systems, food production, and other dimensions of human well-being (Armsworth & Roughgarden 2003; Cardinale *et al*. 2012; Renard & Tilman 2019).

Here, building on the method developed by Ross *et al*. (2023), we propose a new method to empirically quantify response diversity in the context of multifarious environmental change. First, we first (in Sections 3.1-3.3) illustrate this method using simulated data representing species performance curves in cases with and without interactions between different environmental drivers. Then (in Section 3.4), we investigate the role of the **direction of environmental change** in shaping response diversity when multiple drivers of environmental change fluctuate over time. Next (in Section 3.5) we address the case when the direction of the environmental change is unknown—that is, there is no information on how environmental conditions have changed or will change. Finally, in Section 4 we use numerical experiments to investigate how three aspects of species and their environment influence response diversity.

## 3. The principle and its application

In this section we describe a new method to quantify response diversity in the context of multiple environmental variables. We start by explaining the principle and the underlying mathematical concepts. Then, we describe how to calculate response diversity in two important and distinct situations: when the environmental conditions to which species are responding are known, versus when these environmental conditions are unknown. Finally, we discuss how quantifying response diversity when the environmental conditions are unknown can provide us with an estimation of the complete insurance capacity of a system.

### 3.1. The principle

The response diversity of a community can be measured as the diversity of species’ responses to environmental change (McCann 2016; Mori *et al*. 2013). Ross *et al*. (2023) proposed characterising species’ responses with the first derivative of their performance-environment function evaluated over an environmental gradient. By fitting individual species’ performance–environment relationship using **Generalised Additive Models** (GAMs), then taking the **first derivative** of the estimated penalised spline to determine the rate of change of species’ performance curves along the environmental axis, and finally measuring the variation of these first derivatives, the authors provide a flexible methodology to measure response diversity applicable when species responses are linear as well as nonlinear (Ross *et al*. 2023). For example, one can measure the response of each species’ intrinsic growth rate to temperature within a given community, quantify the strength and direction of these responses (e.g., as the first derivative of the response curve), and calculate the diversity of responses as a measure of response diversity (e.g., by calculating variation in the first derivatives among the species in a community). In cases where species responses are nonlinear, the first derivative and therefore response diversity is a function of the environmental state (Fig. 1).

**Figure 1.**
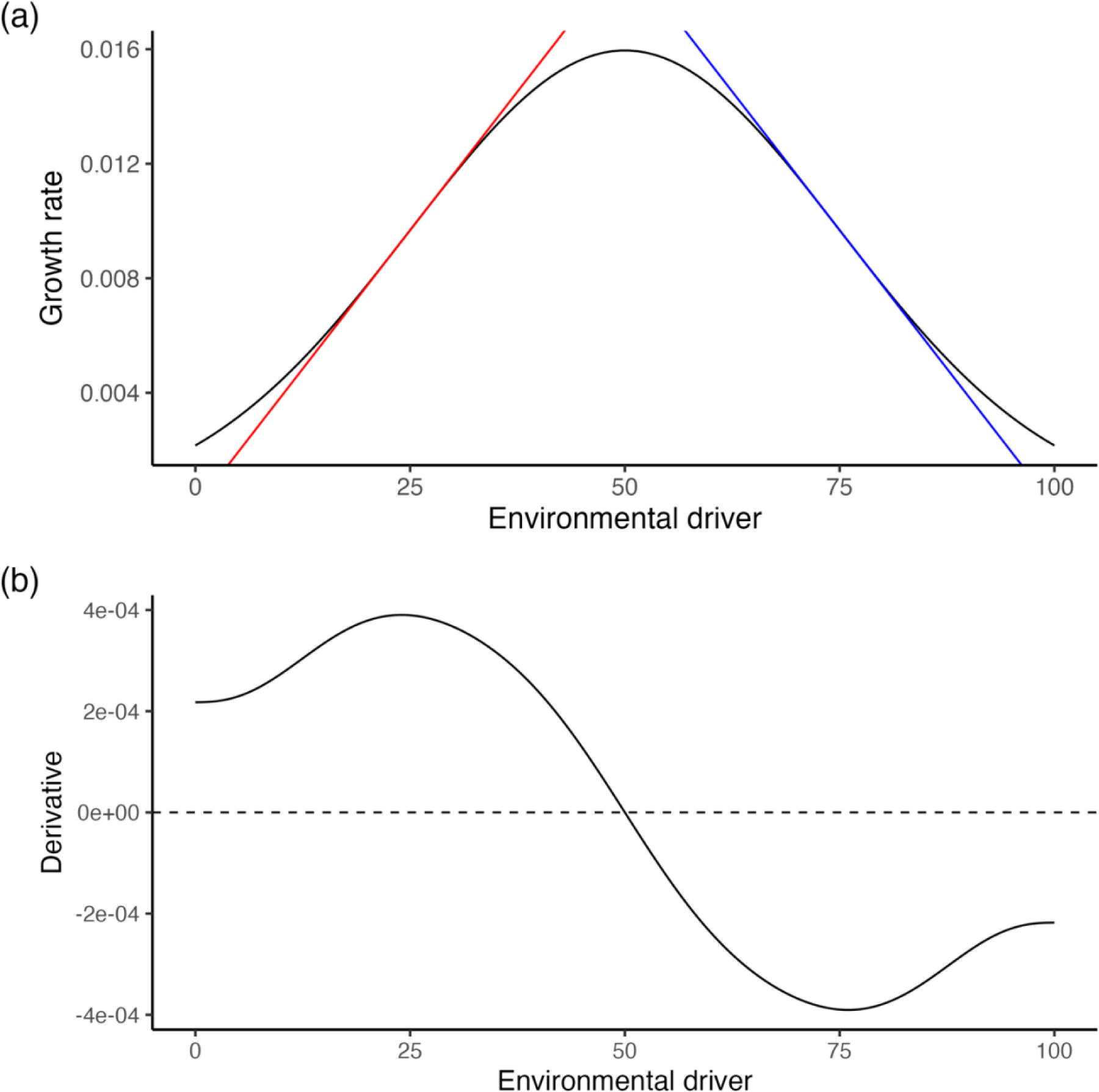
The first derivative of a function depends on the value of the environmental state. The value of the first derivative (slope of the curve, shown at two different values of the environmental gradient by the coloured lines) depends on where along the environmental gradient it is being measured. In (a) we show that not only the value of the first derivative is different depending on whether it is measured when the environmental driver assumes a value of 25 (red tangent line) or 75 (blue tangent line), but it has also an opposite sign. In (b) the first derivative of the curve is plotted along the environmental gradient.

Ross *et al*. (2023) proposed two metrics to quantify variation in species’ performance-environment relationships based on their first derivatives: **response dissimilarity** and **response divergence**. Response dissimilarity captures the total dissimilarity in species’ performance-environment relationships but does not consider whether relationships differ in their direction (that is, whether the performance of one species increases while that of another decreases). Response dissimilarity is thus lowest when all species respond identically, and highest when species responses differ in magnitude (without accounting for direction). In contrast, response divergence accounts for whether performance–environment responses differ in direction (positive or negative responses) (Ross *et al*. 2023). These metrics may be suitable for quantifying response diversity in the context of a single environmental driver but given that multiple environmental drivers change simultaneously in many natural systems (*e.g.*, Bowler *et al*. 2020), a methodological approach that can account for such environmental changes is needed.

Now imagine that the growth rate of a population depends on two environmental factors, such as temperature and salinity. Instead of a two-dimensional performance *curve* modelling performance against one environmental driver, we now have a three-dimensional **performance *surface*** to model performance against two drivers (Fig. 2). We can represent the dependency of growth rate on these two environmental drivers as *G* = *f*(*T*, *S*), where *G* is growth rate (our measure of performance), *T* is temperature, *S* is salinity, and *f* is a function describing the form of these relationships. It may be that the dependencies are linear, nonlinear, and with or without an interaction between temperature and salinity, hence our method must be able to accommodate these phenomena.

**Figure 2.**
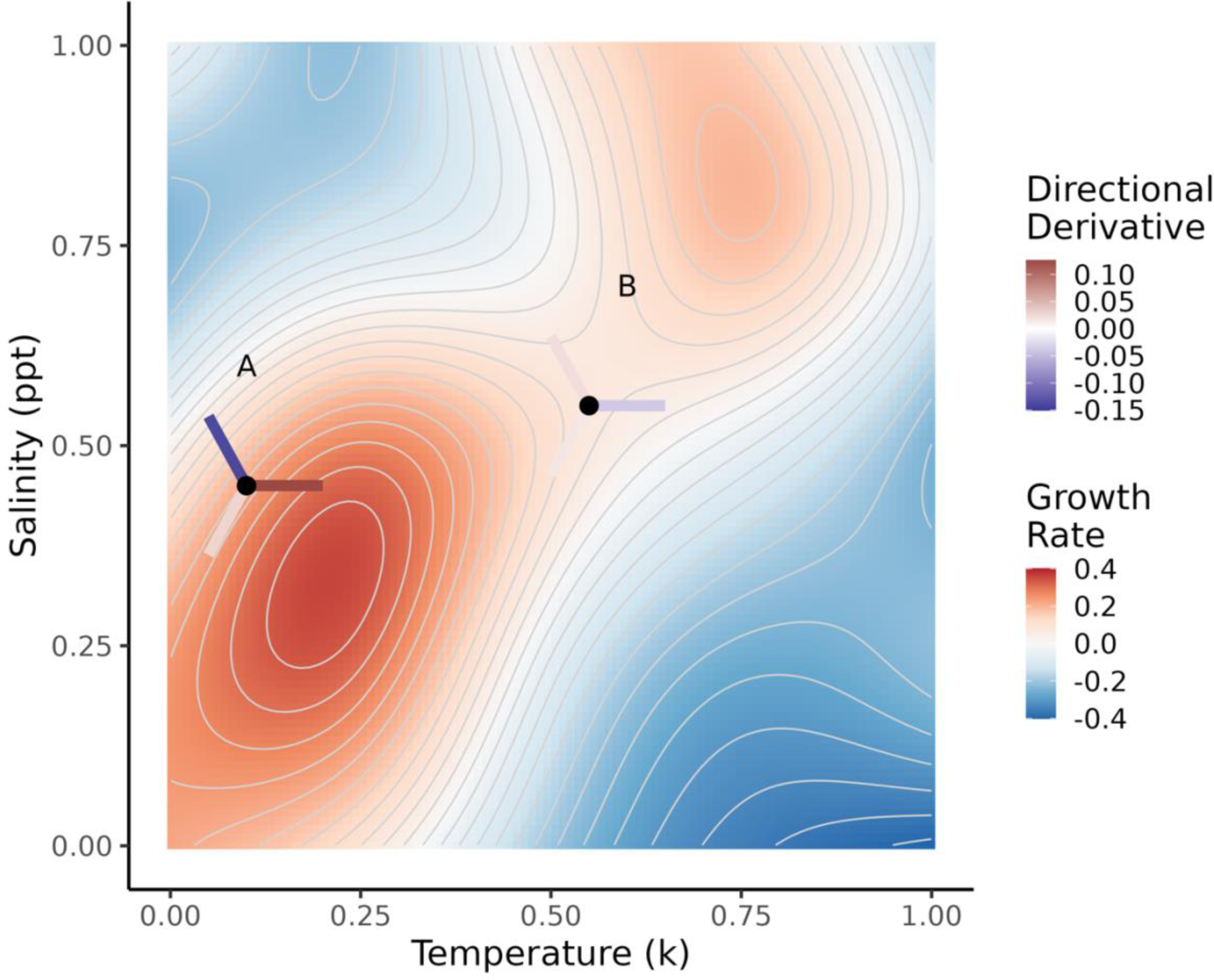
Illustration of a performance surface and some possible directional derivatives. The growth rate of a species is determined by changes in temperature and salinity. This growth rate-environment slope then depends on the location on the performance surface and the direction on the surface, and hence is termed the directional derivative. In the illustration, two cases are shown: Location A is on a “steep” side of the performance surface (i.e. of the hill). Depending on the direction from Location A the slope (i.e. directional derivative) can be positive (uphill), negative (downhill) or zero (traverse). In B, depending on direction, the growth rate-environment slope can, again, be positive, negative, or zero, but it will always be smaller than any directional derivatives from A, as B is on a “flatter” portion of the performance surface.

The solution we developed to account for these multiple environmental dependencies is a relatively straightforward extension from one to two dimensions of environmental change (and can in principle be extended further to many dimensions). Instead of calculating the first derivative (and measuring interspecific variation of these derivatives), we calculated the **directional (first) derivative**. A *directional* derivative is the slope on a performance surface at a particular location pointing in a particular direction. Imagine standing on a hill. You could be standing on a particularly steep part of the hill, or on a quite flat part, i.e., the slope depends on location. Furthermore, the slope you experience can depend on the direction you face. If you are standing on a side of the hill, the slope is zero when you face along a traverse but is positive when you face uphill.

Translating this into environmental space, such as variation in temperature and salinity (Fig. 2), the location on the surface is given by a particular value of temperature and salinity (**environmental location**). The direction on the surface is given by a vector containing the amount of change in temperature and amount of change in salinity. For example, the vector [1, 0] would be 1 unit change in temperature and zero in salinity. Whereas the vector [1/√2,1/√2] would be 1/√2 change in both temperature and salinity (and the vector would be of length 1, i.e., a **unit vector**). To find the directional derivative we must, therefore, specify both a location and a direction on the performance surface (see further details in Section 3.3).

### 3.2. Calculating the partial derivatives

The first step in calculating a directional derivative (i.e. the slope on a surface) is to calculate the partial derivatives of the surface. Mathematically, a partial derivative of a function of multiple variables is its derivatives with respect to one of these variables, with the remaining variables held constant. A partial derivative with respect to a specific variable can be conceptualized as the rate of change of the function in the direction of the variable considered. Coming back to the above example, where the growth rate of a species is affected by temperature and salinity, the partial derivative with respect to temperature represents the change in growth rate as temperature changes, when salinity is held at a constant value. Similarly, the partial derivative with respect to salinity holds temperature constant and measures the change in growth rate as a function of salinity at that fixed temperature. In other words, we analyse the effect of temperature or salinity change along a slice of the performance surface (Fig. 3).

**Figure 3.**
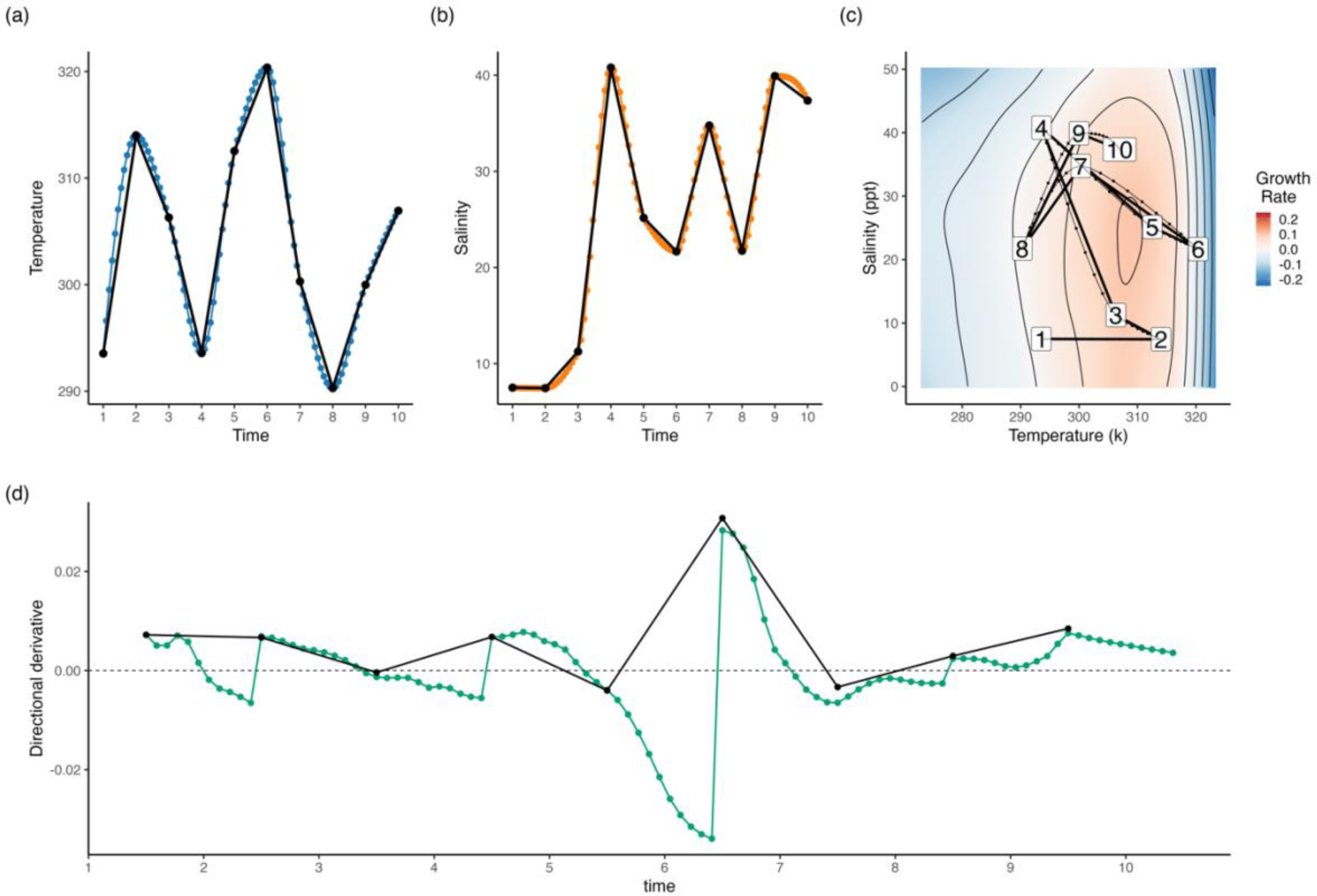
Calculating partial derivatives on a performance surface. (a and d) The performance surface for the growth rate (blue colours = negative growth rates, red colours = positive growth rates) of a focal species. (a) The black solid line represents the value of salinity at which the partial derivative with respect to temperature is then calculated. Holding salinity constant at 20 ppt, allows measuring: (b) a two-dimensional response curve reflecting the effect of temperature on the focal species’ growth rate when salinity is held constant at 20 ppt, and (c) the partial derivative with respect to temperature, when salinity is held constant at 20 ppt. (d) The black solid line represents the value of temperature at which the partial derivative with respect to salinity is calculated. Holding temperature constant at 300 K allows measuring: (e) a two-dimensional response curve reflecting the effect of salinity on the species’ growth rate when temperature is held constant at 300 K, and (f) the partial derivative with respect to salinity, when temperature is held constant at 300 K.

Following Ross *et al*. (2023), we use a GAM to fit species’ performance surfaces to multifarious environmental change using the mgcv (version 1.8.41; Wood 2011) package for R (R version 4.2.2; R Core Team 2022) and calculated the partial derivatives of the estimated surface using the gratia (R package version 0.8.1.46, https://gavinsimpson.github.io/gratia/; 13/12/2023 10:02:00 package for R. GAMs allow fitting performance surfaces of the response variable using tensor-product smooths. Tensor-product smooths are a mathematical way of representing nonlinear interactions between two or more predictors (environmental driver variables) and are a particularly suitable approach when predictors are measured on different scales.

### 3.3. Calculating directional derivatives

After calculating each of the partial derivatives, these are then summed in proportion to the amount of change in each of the environmental variables to give the directional derivative. “Proportional to the amount of change in each of the environmental variables” is akin to saying, “in the direction of environmental change”. Hence, a directional derivative describes the rate of change of a function at a particular location in a defined direction (*e.g.,* the change in the growth rate of a species as multiple environmental variables change). Calculating a directional derivative therefore requires knowledge of the direction of environmental change, and this is encoded in a unit vector. The direction can be arbitrary, but in an ecological context, the direction will be dictated, in most cases, by the temporal trajectory of change in the environmental conditions of interest.

Translating this into our example, to calculate the slope (directional derivative) at a location on the performance surface (a species’ response to changes in temperature and salinity), we need to know the location on the surface (e.g., this could be the current environmental conditions) and the direction of change in the environment (how the environmental drivers change over time). The direction of change in various environmental drivers can be derived from historic or projected environmental time series. However, in some cases, the direction of environmental change may be unknown, but one may nonetheless be interested in calculating response diversity. In the next section we will analyse these two cases separately.

For a mathematical perspective, consider for example a species performance surface with growth rate as the dependent variable (which determines the height of the surface) and temperature (*T*) and salinity (*S*) as the predictor variables. To calculate a directional derivative, we give a location on the surface, e.g., (*T*_0_, *S*_0_), and give the direction of environmental change in unit vector form û = 〈*U*_*T*_, *U*_*S*_ 〉. Furthermore, let the two partial derivatives be *f*_*T*_(*T*_0_, *S*_0_): the partial derivative of *f*(*T*_0_, *S*_0_) with respect to temperature (*T*), and *f*_*S*_ (*T*_0_, *S*_0_): the partial derivative of *f*(*T*_0_, *S*_0_) with respect to salinity (*S*). The directional derivative is then given by *f*_*T*_ (*T*_0_, *S*_0_)*U*_*T*_ + *f*_*S*_ (*T*_0_, *S*_0_)*U*_*S*_.

### 3.4. Known direction of environmental change

If the direction of environmental change is known, then calculating directional derivatives is straightforward and proceeds exactly as described above. For example, in Figure 4, we can see how temperature and salinity change over time (Figs. 4a and 4b). This allows us to plot the trajectory of environmental change on a species’ performance surface (Fig. 4c), which gives us a visual representation of the direction in which the directional derivatives are calculated when we follow the direction of environmental change. As temperature and salinity change over time on the species’ performance surface, the trajectory traverses an area of the surface where the species’ growth rate increases, and areas where it decreases (Fig. 4c). Intuitively, this results in changing values of the directional derivative over time depending on the trajectory of the environmental change; values are positive or negative in Figure 4d when passing over, respectively, positive, or negative growth rate areas of the performance surface (Fig. 4c).

**Figure 4.**
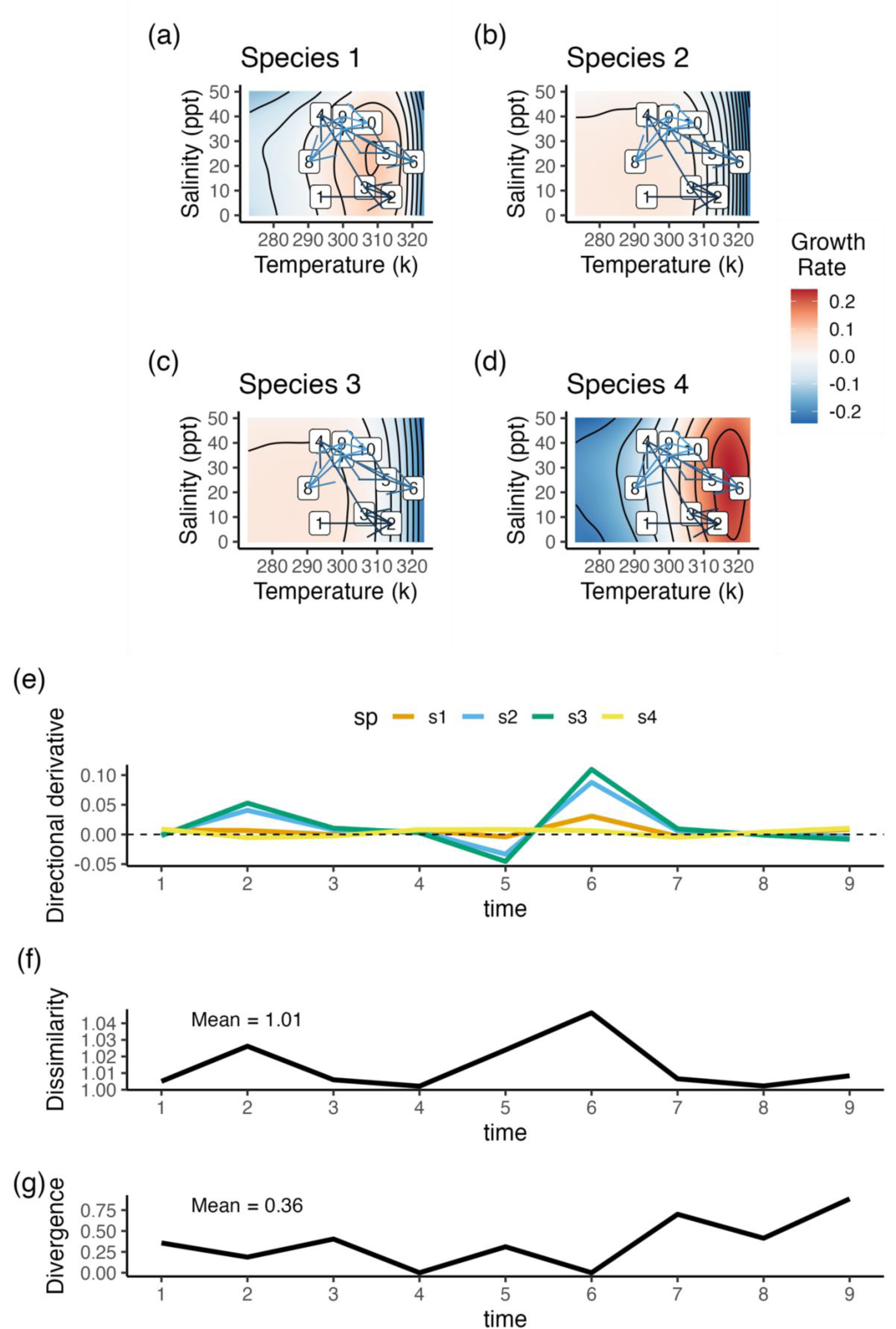
Calculation of directional derivatives for one species for a time series of environmental change. (a) Time series of temperature change. (b) Time series of salinity change. (c) Time series of temperature and salinity change overlayed on the species performance surface. Sequential numbers in white boxes represent positions on the performance surface at time points corresponding to (a) for the temperature axis and (b) for the salinity axis. Colours indicate the growth rate. (d) The time series of directional derivatives corresponding to the time series of environmental change in (a-c) and the species performance surface in (c). In (a) and (b) the colour lines and dots represent the interpolated time series of environmental change. Interpolation was done using cubic spline interpolation and sample size with n = 100. In (c), the thinner lines connecting the black dots represents the interpolated time series of temperature and salinity change on the species response surface. In (d), the coloured line represents the interpolated time series of directional derivatives over time. The interpolation was performed to estimate values that lie within two consecutive “measured” time points but were not explicitly measured. Performing the interpolation, we can estimate the smooth change in the directional derivative as the system moves from t =1, to t =2. Please note that the offset between “measured” points and the time on the x-axis is meant to represent the directional derivative as we go from t = 1 to t = 2.

In many real experimental and field studies, where time and resource limit the number of sampling one can perform, researcher will have only some time points measured for both the environmental conditions as well as the species’ traits (e.g. growth rate). This could result in only few points on the surface where the directional derivatives are being calculated. Let us consider the example of the directional derivative calculated at t =1 in Fig. 4d. Here we are calculating the directional derivative for a unit vector at (1) in the direction of (2) on the surface. This represents the instantaneous derivative (slope of the surface) at (1) in the direction of (2). Yet, the system does not just jump from (1) to (2) between t = 1 and t = 2, but rather it moves over the surface from (1) to (2). For didactical purpose, and to clearly deliver the principle, we use here interpolation to show that the slope of the surface is changing smoothly (not linearly) as we moved from one environmental location (i.e. the one at t = 1) to another (i.e. the one at t = 2).

In a community context, one can calculate the directional derivative over time in the direction of the environmental change for each individual species. Next, by applying the response dissimilarity and response divergence methods described by Ross *et al*. (2023), one can measure the variation in the directional derivatives of several species within a community to estimate response diversity (Fig. 5).

**Figure 5.**
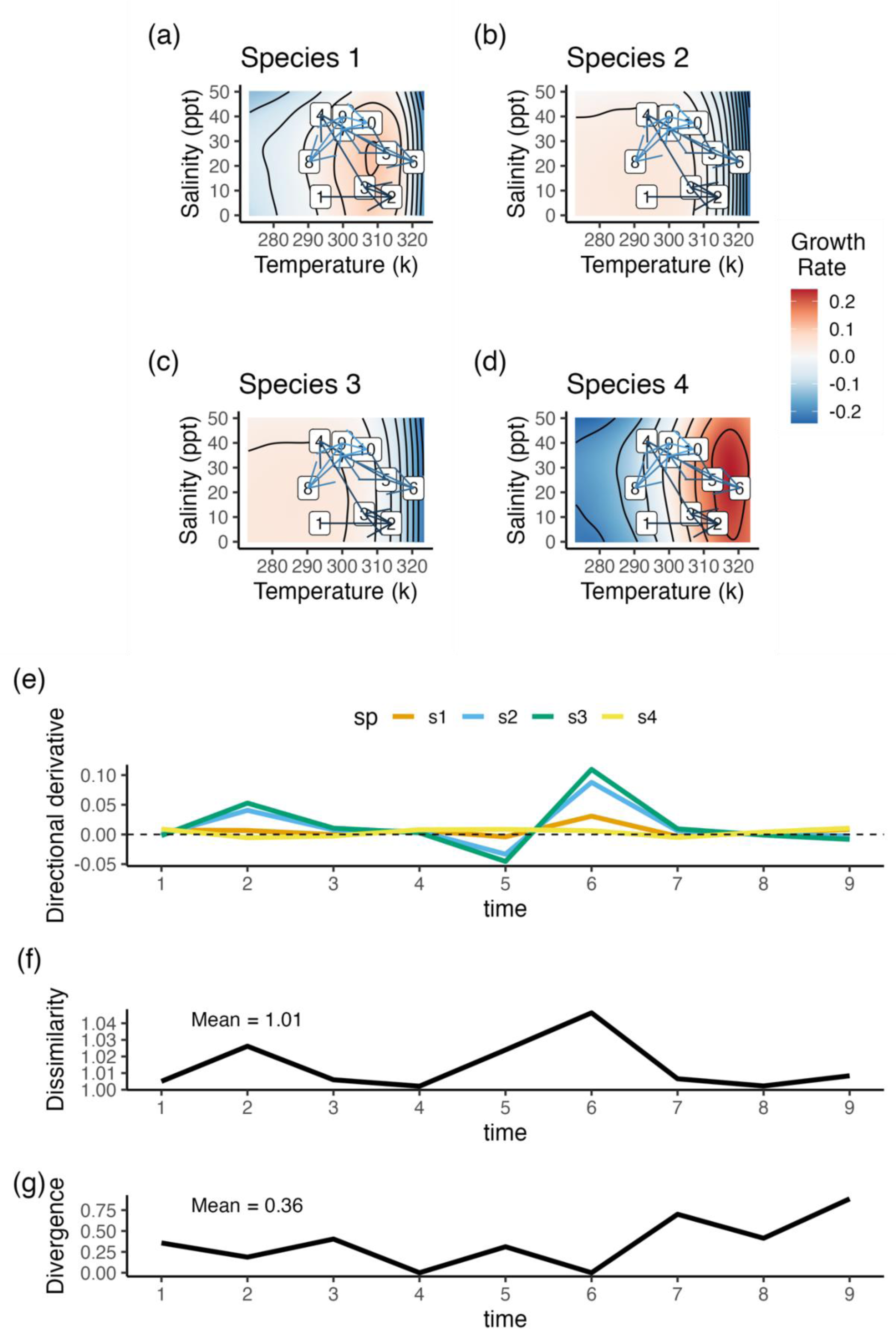
Response diversity calculation using performance surfaces for multiple species. (a-d) Performance surface of four species composing a community, with growth rates (colours, positive = red, negative = blue) that vary along salinity (Y-axes) and temperature (X-axes) gradients. Direction of environmental change is displayed on the surfaces as arrows and numbered time points (following Fig. 4). (e) Directional derivatives of each of the four species (a-d) over time. Response diversity is calculated from these multi-species directional derivatives as: (f) response dissimilarity, and (g) response divergence.

### 3.5. Unknown direction of environmental change

When applying response diversity to empirical data, the direction of change in environmental conditions may be unknown. For example, if dealing with future predictions, uncertainty may be too high to reliably forecast future environmental change.

In this section, we explore one way to calculate response diversity when the direction of the environmental change is unknown, by quantifying response diversity to all possible environmental changes and then taking the mean. Consider a single location (corresponding to a specific set of environmental conditions) on the performance surface of a species, whose growth rate is, again, determined by changes in temperature and salinity. Since we do not know how the environmental drivers are going to change, we can calculate the directional derivative for this specific location on the performance surface in many possible directions (Fig. 6).

**Figure 6.**
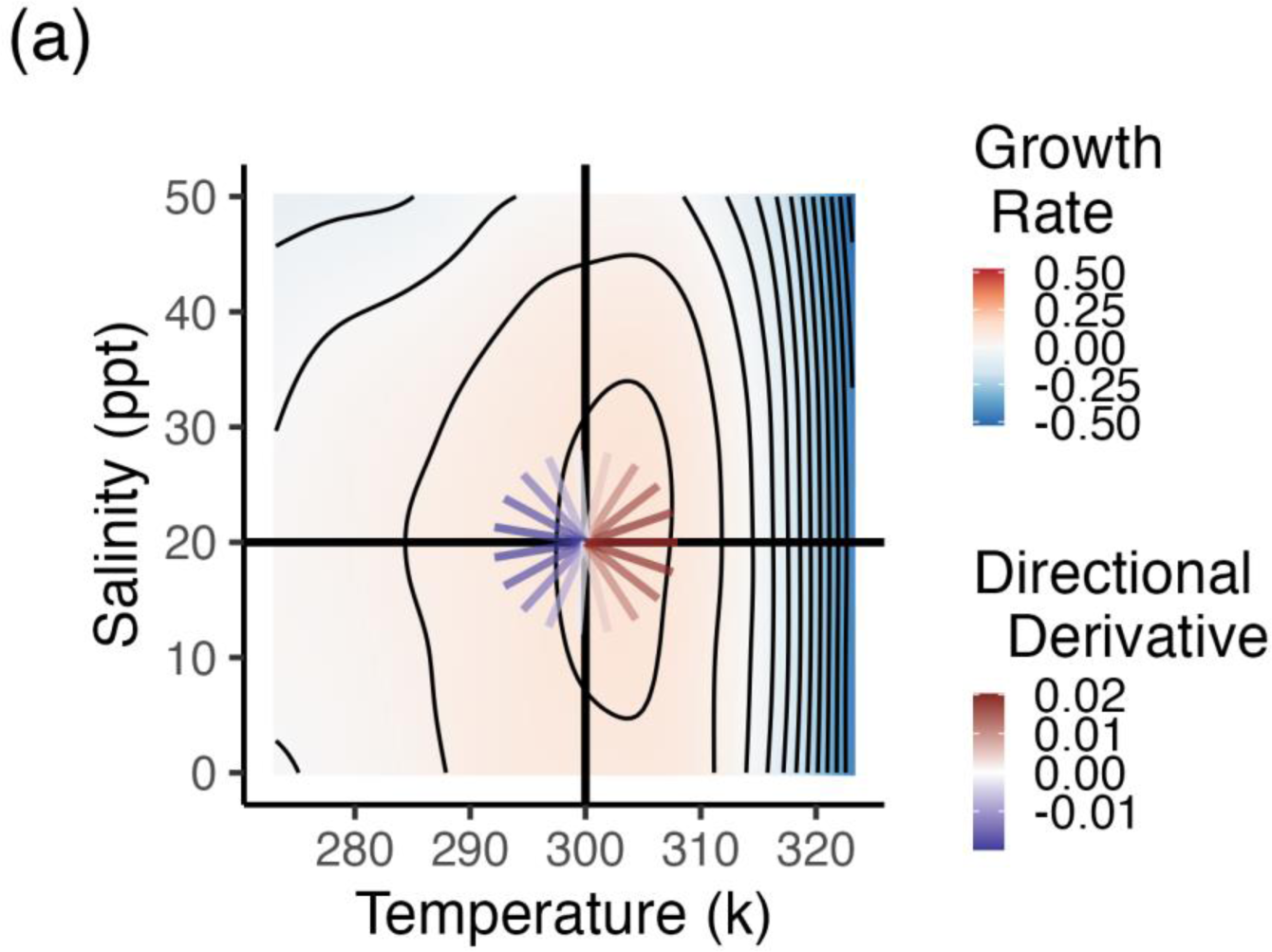
Directional derivatives calculated in many directions for a point on one species’ performance surface. From a specific point on the performance surface (salinity = 20 ppt and temperature = 300K), we can calculate the directional derivatives in many possible directions. Since the selected point is in the proximity of the maximum growth rate (i.e. top of the hill), all the directional derivatives calculated in the direction of the maximum (i.e. right side, “uphill”) are positive. Conversely, the directional derivatives calculated in the opposite direction of the maximum (left side, “downhill”) are negative.

Scaling up to a community context, we can do the same for each species in the community. Next, we calculate the variation (diversity) in the directional derivatives at each location in each direction. Finally, we calculate the diversity across all possible directions and locations, this gives us a **potential response diversity** value for a community (Fig. 7).

**Figure 7.**
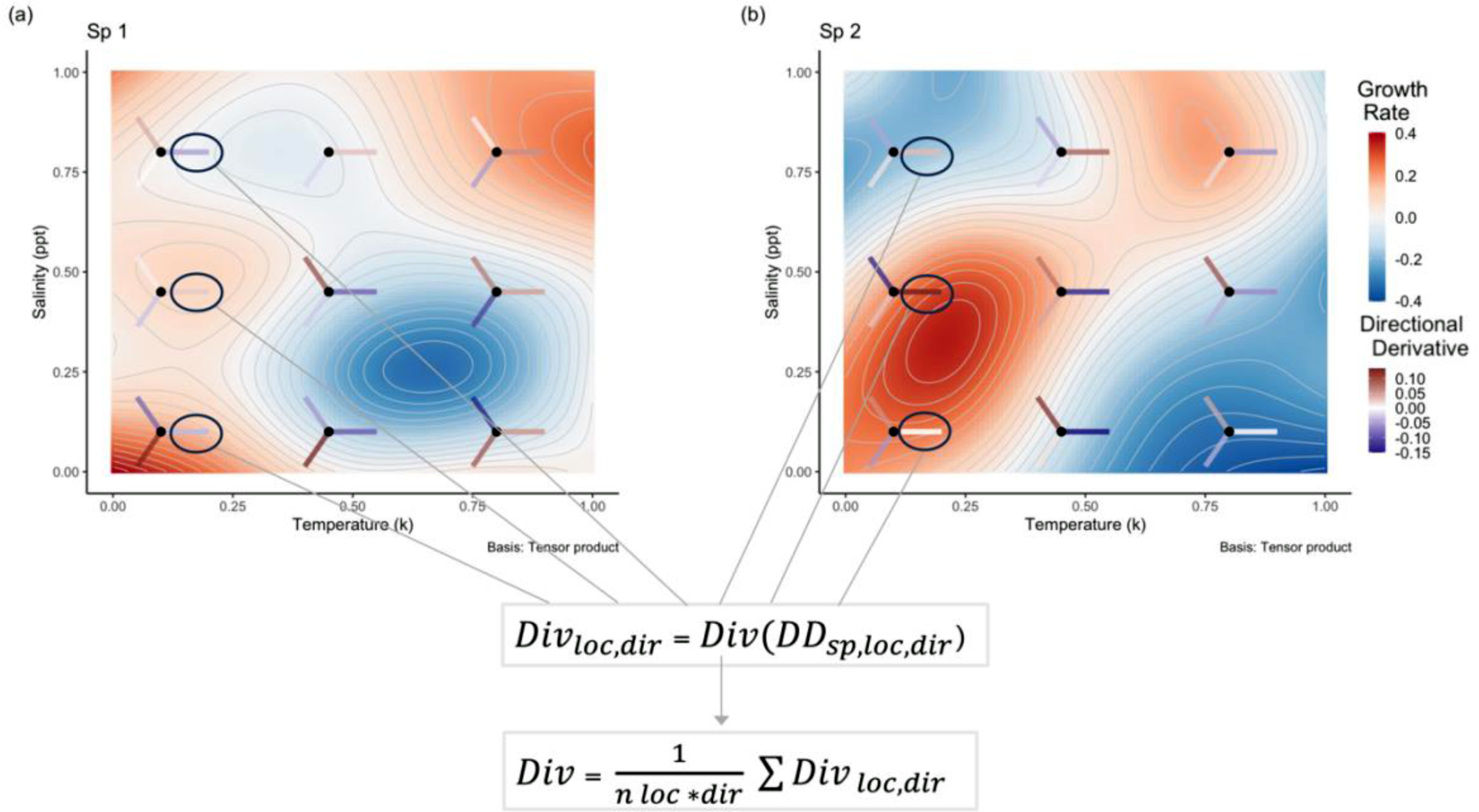
Illustration of the principle underlying the calculation of potential response diversity. Considering the difficulties related to displaying multiple 3D performance surfaces, each having multiple points with directional derivatives going in all possible directions, we focus here on representing only two species and nine points on each surface. For those points, we only display three directional derivatives to help visualise the calculation process, but please note that computationally this is done for every possible combination of locations and directional derivatives.

To be clear, the steps to calculate the potential response diversity can be summarised as follow:

1. Calculate every *DD*_*sp,loc,dir*_, that is the directional derivative (*DD*) of each species (*sp*) in each location (*loc*) in each direction (*dir*).
2. Calculate *Div*_*loc,dir*_ = *Div*(*DD*_*sp,loc,dir*_), that is, the diversity of the directional derivatives for a given location and direction across species.
3. Calculate 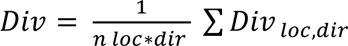, that is the average diversity across locations (*n loc* represents all the locations on the surface) and directions.

### 3.6. Discussion on potential response diversity

The framework proposed here for measuring response diversity when the direction of environmental change is unknown can be used to systematically measure response diversity to all potential environmental changes. The result of this analysis therefore represents potential response diversity, which captures the complete insurance capacity of a system under all possible environmental conditions. Potential response diversity can be seen as a comprehensive quantification of response diversity that does not depend on knowing the direction of the environmental change (e.g., does not require knowledge of future environmental conditions). By averaging across all possible conditions, no specific information on the direction of environmental change is needed to obtain a quantification of the insurance capacity of a community.

Even though here we quantify response diversity to two environmental drivers, our approach can be extended to cases where three or more drivers influence species traits. Its application is possibly very broad. Potential response diversity can be used to assemble communities with the highest possible response diversity to all potential directions of environmental change, perhaps providing an operationalizable way to promote stable ecosystem functioning through time (Mori *et al*. 2013). Assembling communities specifically designed to have high potential response diversity would allow ecosystem managers to promote communities and ecosystems able to maintain stable levels of functioning in face of environmental change. Assembling communities with different potential response diversity may also be used to test various ecological questions. For example, communities assembled along a gradient of potential response diversity could be used to directly test the link between response diversity and temporal stability, providing new insight into possible mechanistic links between response diversity and temporal stability in a way that gets beyond the context dependence of specific environmental change scenarios.

## 4. Investigating the drivers of response diversity

In this section we use numerical experiments to explore how three features of species and their environment affect patterns of response diversity. We term each of these three features a “treatment” in our numerical experiment. The three treatments were:

Treatment 1: The relative importance of the two environmental drivers. There were two levels of this treatment: both equally important; and one driver more important than the other one. The mathematical meaning and implementation are described below.

Treatment 2: The correlation pattern in the diversity of species’ responses to the two drivers. There were three levels: no correlation, positive correlation, and negative correlation (details below).

Treatment 3: The temporal means of each of the two environmental variables. There were three levels: low, medium, and high mean values (details below).

We selected these three treatments from numerous possible others because they are illustrative and interesting. These treatments are potential drivers of response diversity that have been previously discussed and that have good conceptual foundations (Laliberté *et al*. 2010; Mori *et al*. 2013; Ross *et al*. 2023). Moreover, these treatments are testable in controlled empirical studies (Leary & Petchey 2009). Nevertheless, numerous other treatments could be conceived. For example, other features of species’ performance to the environment (e.g. different shapes of species performance curves; co-tolerance and anti-tolerance to multiple drivers), and different scenarios of environmental change (e.g. different magnitude of fluctuations in the environmental variables; different correlation in temporal change of the environmental variables) may drive response diversity. While investigating these and other treatments would be interesting, we consider the three chosen treatments to be suitable and sufficient to illustrate drivers of response diversity in a multifarious context.

Before describing each treatment in detail, we first describe how we simulated species performance surfaces.

### 4.1. Simulating species performance surfaces

To test our new methods, we simulated species growth rate under the influence of two environmental drivers. Numerous mathematical functions have been used to represent how organismal performance changes with an environmental driver (Bernhardt *et al*. 2018; Kremer *et al*. 2017). Moreover, multiple mathematical functions have been used to represent an interactive effect of two or more environmental drivers on species performance (e.g. Thomas *et al*. 2017).

For its widespread application and ease of implementation, we used here the Eppley equation (Formula 1: Eppley, 1972). The Eppley equation captures the exponential relationship between maximum growth rate and temperature. It was derived from empirical data on phytoplankton and has become a fundamental component of our understanding of how primary productivity is constrained by temperature (Kremer *et al*. 2017). The Eppley equation depicts typical features of organismal responses along a thermal gradient, such as an exponential increase in growth rate moving along the environmental gradient until reaching the maximum grow rate, and a sharp decline in growth rate after that. This pattern is often termed a thermal performance curve, where peak growth rate occurs within the biologically relevant temperature range for the organism (DeLong *et al*. 2018).

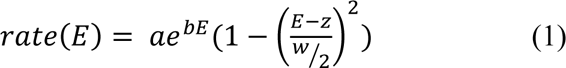

Here, *E* is the value of the environmental factor, *z* controls the location of the maximum value of the growth rate, *w* controls the range of *E* (environmental driver) over which the rate is positive, *a* is a scaling constant, and *b* controls the rate of increase of the growth rate towards the maximum rate as *E* increases.

The original Eppley function was developed to simulate species response to only one environmental variable. We thus adjust it here to allow for the effects of two simultaneous environmental drivers on species responses (formula 2).

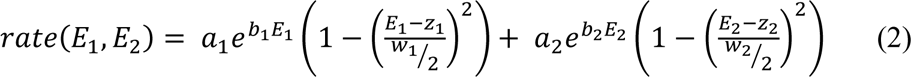

From the above formula, it is clear the effect of *E*_1_ and *E*_2_ is defined as additive. For example, the value of *E*_2_ does not affect the value of *E*_1_ at which the rate is maximised, and vice-versa (Figs. 8a and 8b). We can also introduce an interaction to this formula. We simulated all the scenarios and combinations described above for both additive and interactive environmental effects on species’ growth rate. However, we found no qualitative difference. We introduced an interaction by having the value of E1 at which the rate is maximised depend on the value of E2. This way of implementing an interactive environmental effect does not fundamentally change the shape of the performance curves, and thus the range of values the derivatives can assume.

**Figure 8.**
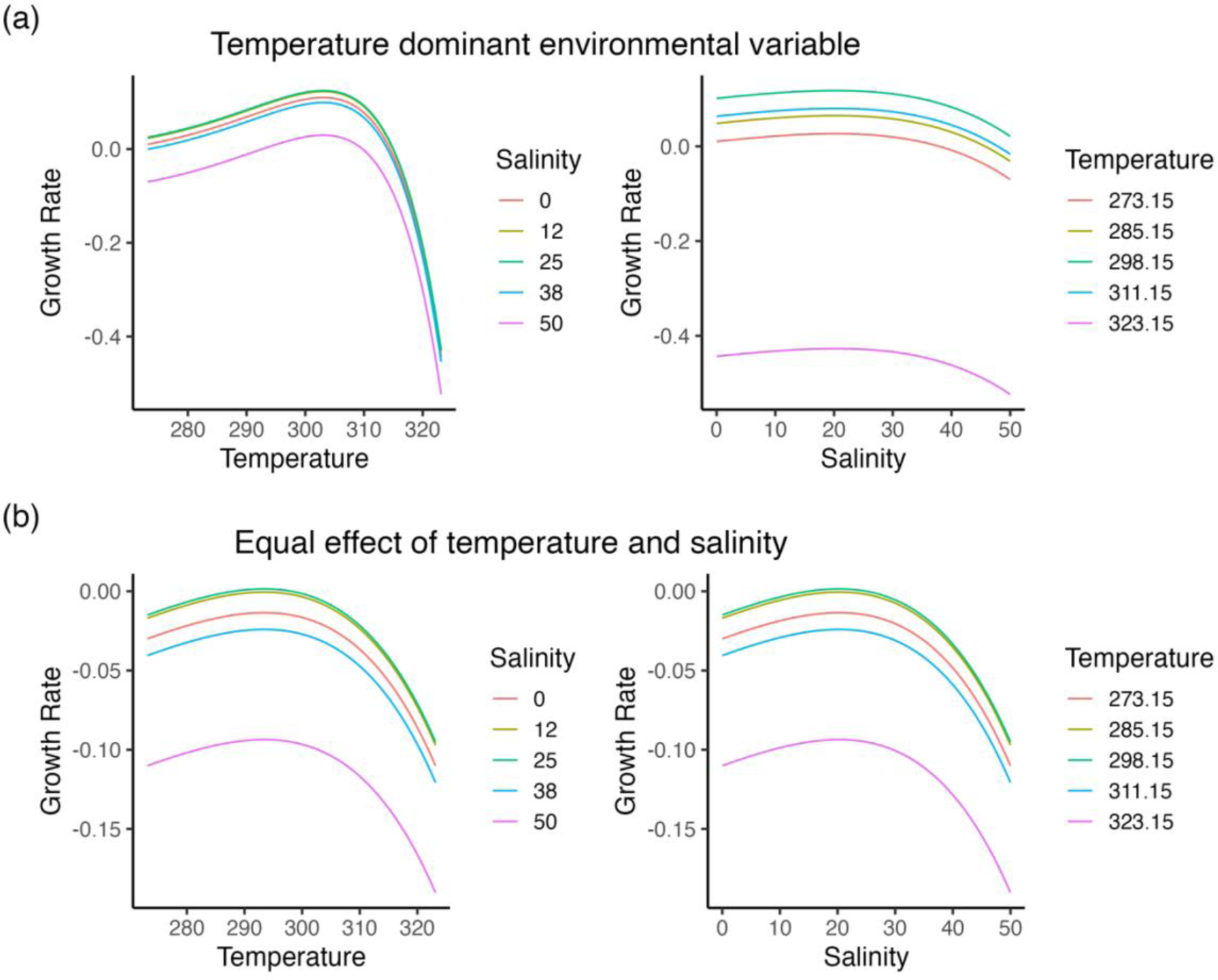
Simulated species growth rate responses to changes in temperature and salinity. (a) shows the case where temperature has a larger effect than salinity (compare the slope of the growth rate-temperature relationship in left panel to the weaker growth rate-salinity relationship in right panel). (b) shows the case where salinity and temperature have the same effect on species’ growth rate.

Accordingly, we report only the results obtained using the additive effect. For details on how we simulated the interactive environmental effect, please see the GitHub repository (https://github.com/opetchey/multifarious_response_diversity/releases/tag/v0.1).

### 4.2. Simulating treatment 1

Even when more than one environmental driver affects a species’ performance, often one environmental driver has a larger effect size than another (Schulhof *et al*. 2019). For example, the growth rate of a species might change more in response to temperature than salinity. Since temperature and salinity have different scales, we are concerned by the response of growth rate given the observed amount of change in temperature and salinity (Fig. 8).

To create treatment 1 (difference in the relative importance of the two environmental variables) we set different values for parameters of the temperature performance curves. To make temperature have a relatively large effect on species’ growth rate (Figs. 8a and 8b) we set *a*_1_ = 1e^-9^ (*a* is the overall scaling constant), *b*_1_ = 0.063 (b controls the rate of increase), *z*_1_ = 285 (z controls the location of the maximum response), *w*_1_ = 60 (*w* controls range of environmental conditions over which the response is positive). To make the case with equal importance (Figs. 8c and 8d) we set *a*_1_ = 1e^-3^, *b*_1_ = 0.02, *z*_1_ = 293, *w*_1_ = 10. In both cases the salinity parameters were *a*_2_= 1e^-3^, *b*_2_= 0.02, *z*_2_ = 20, *w*_2_ = 10.

### 4.3. Simulating treatment 2

Treatment 2 concerns the correlation pattern in the diversity of species’ responses to the two drivers. There were three levels: no correlation, positive correlation, and negative correlation. In the no correlation treatment level, we created communities that differed in the amount of interspecific variation in temperature optima (Fig. 9a, d, and g), but not in salinity optima (Fig. 9b). In this no-correlation level, diversity in responses to salinity was fixed at an intermediate value and thus did not correlate with the variation in diversity in temperature responses (Fig. 9c). We simulated three communities with low, medium, and high diversity in species responses to temperature, respectively, and all with medium diversity in salinity responses.

**Figure 9.**
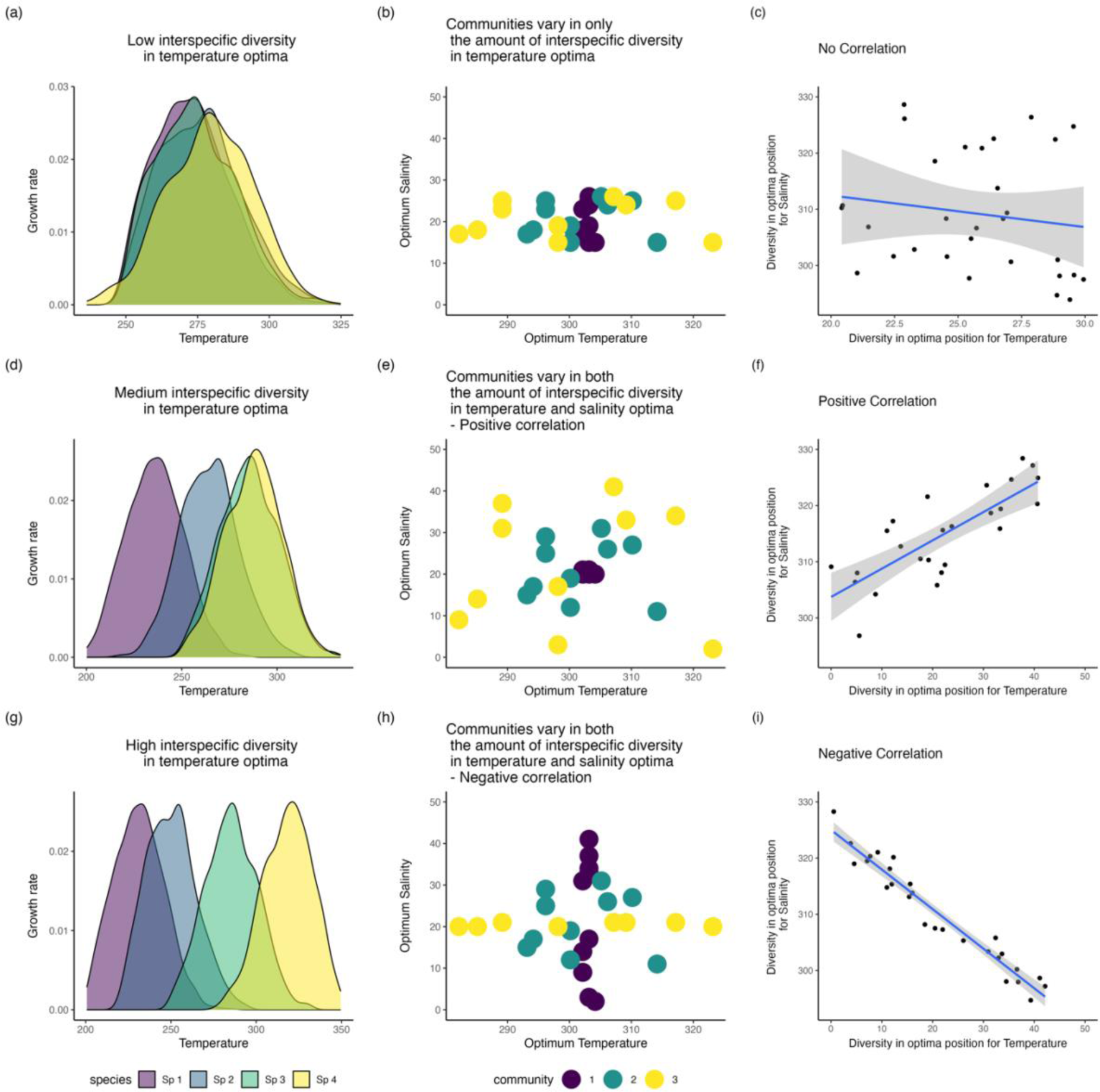
Treatment 2: Amount of and correlation in diversity of species’ responses to salinity and temperature. Each of (a), (d), and (g) show the performance curves of four species, with low (a), medium (d), and high (g) interspecific variation in temperature optimum. Panels (b), (e), and (h) illustrate different patterns of correlation in interspecific variation in temperature optima. Each coloured circle represents a species; the colour of the circle shows which community the species belongs to; the x-and y-coordinate shows the temperature and salinity optima of that species. Panel (b) illustrates the no correlation treatment. All three communities have a fixed and rather low amount of variation in the position of the salinity optimum (y axis), whereas the variation in the position of the temperature optimum increases from low in community 1 (purple), to medium in community 2 (green), to high in community 3 (yellow). Panel (e) illustrates the positive correlation treatment. Community 1 (purple) has low variation in the position of the optima for both salinity and temperature, community 2 (green) has intermediate variation in the position of optima for both temperature and salinity. Community 3 (yellow) has high variation in the position of the optima for temperature and salinity. Since the variation in the position of the optima gradually increase across the communities for both temperature and salinity, this the positive correlation treatment. Panel (h) illustrates the negative correlation treatment. Community 1 has high variation in the position of the optima for salinity, but low for temperature. Community 2 has intermediate variation in the position of the optima for both temperature and salinity, and community 3 has high variation in the position of the optima for temperature, but low for salinity. Since, in this scenario, when variation in optima position for salinity is high, variation in optima position for temperature is low and vice versa, we call this the negative correlation treatment. (c), (f), (i) exemplify the correlation between diversity in the optima position for salinity (y axis) and temperature (x axis).

In the positive correlation treatment level, the communities had low, medium, and high diversity in their responses to both temperature and salinity (Fig. 9e). I.e., the diversity of temperature and salinity responses were positively correlated across the three communities (Fig. 9f). In the negative correlation treatment level, there was one community with high diversity in salinity responses but low diversity in temperature responses, one community with intermediate diversity in response to both drivers, and one with low diversity in salinity responses but high diversity in temperature responses (Fig. 9h). This scenario is thus the negative-correlation scenario because response diversity to temperature and salinity were negatively related across the three communities (Fig. 9i).

### 4.4. Simulating treatment 3

Treatment 3 concerns the temporal means of each of the two environmental variables. There were three levels: low, medium, and high. First, we explain this treatment in the context of a single environmental variable (Fig. 10), which may have a low, medium, or high temporal mean (Fig. 10a-c). Importantly, this mean is low, medium, or high *relative to the species performance curves* (Fig. 10d, e).

**Figure 10.**
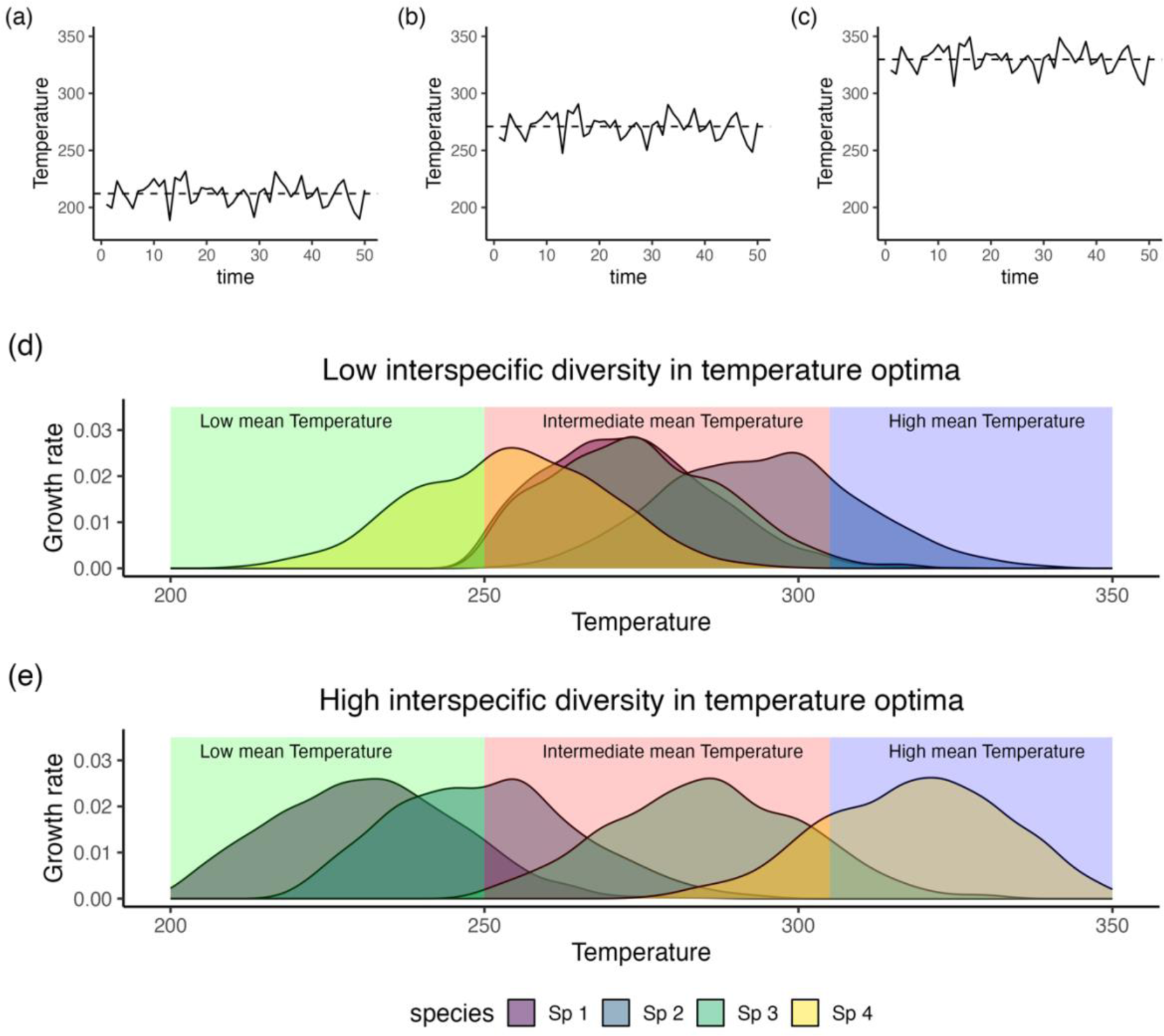
Variation in the mean value of the environment and its relationship with diversity of species’ optimum positions. (a) Temperature fluctuates with low mean values. (b) Temperature fluctuates with intermediate mean values. (c) Temperature fluctuates with high mean values. (d) Community with low diversity in species’ optimum growth rates as related to temperature. (e) Community with high diversity in optimum growth rates as related to temperature. In (d) and (e), shaded areas in different colours show the intervals in which temperature is fluctuating depending on its mean value scenarios.

Let us consider an example of a community composed of species with relatively little diversity in their performance curves, and optima close to the centre of the possible range of temperatures (Fig. 10d). When this community is exposed to temperatures in the centre of the range of possible temperatures, then there should be some response diversity (pink region in Fig. 10d). In contrast, when exposed to temperatures which are relatively low or high, then there should be very low response diversity (green and purple regions in Fig. 10d).

In a contrasting situation where species have very different optima for temperature (Fig 10e), the mean value around which the environmental variable fluctuates has less of an effect on response diversity since species optima (i.e. temperature at which their performance is maximized) are distributed across a wider range of temperatures. Thus, independently of the mean value around which temperature fluctuates, species will inevitably show different growth rates (i.e. they will show different responses to temperature).

When species are exposed to multiple environmental drivers simultaneously, the response diversity of a community will depend on the diversity in the responses to both environmental drivers as well as the mean value around which these drivers fluctuate. To quantify the effect of the mean value of the environmental drivers on multifarious response diversity, we manipulated the mean value around which temperature and salinity fluctuate.

### 4.5. Predictions

Treatment 1: When the two environmental drivers have equal relative importance, we expect that response diversity will be affected by features of performance curves to both drivers (e.g., amount of interspecific variation in species’ optima). When one environmental driver has high relative importance (and the other therefore low importance), we expect response diversity to be driven by patterns in only the responses to the dominant driver.

Treatment 2: We expect response diversity to be highest with positive correlation in diversity of species responses to the two environments. This should be especially so when both environments have equal relative importance (Treatment 1).

Treatment 3: We expect highest response diversity when the temporal mean of the environmental variables is at intermediate values, with respect to the location of the performance curves (Fig. 10). That is, we expect highest response diversity when the response curves of some species have their optimum above the mean environmental condition, and some below the mean.

### 4.6. Effects of the treatments 1 and 2 on potential response diversity

Since treatment 1 (relative driver importance) and treatment 2 (responses – driver correlation) can influence potential response diversity (unknown direction of environmental change), we here report the effects of these treatments only on potential response diversity.

Our simulations revealed that diversity in species’ responses to the environmental variables and correlation pattern in the diversity of species’ responses to the two drivers (treatment 2) were major drivers of potential response diversity (Fig. 11). However, the effects depended on whether temperature was a stronger environmental driver of species growth rates (Fig. 11a), or whether temperature and salinity equally affected growth rates (Fig. 11b; that is, they depended on treatment 1). When temperature was the dominant driver of species’ growth rate, potential response dissimilarity was highest in communities with high diversity in species’ responses to temperature, irrespective of whether diversity in species’ responses to salinity was high, intermediate, or low (Fig. 11a). When temperature and salinity equally influenced species’ growth rate though, potential response dissimilarity was highest when the diversity of responses to both temperature and salinity were high (Fig. 11b). In this case, when only either diversity in responses to salinity or to temperature was high, potential response dissimilarity took intermediate values. Potential response dissimilarity values were lowest when diversity in species’ responses to temperature and salinity was low.

**Figure 11.**
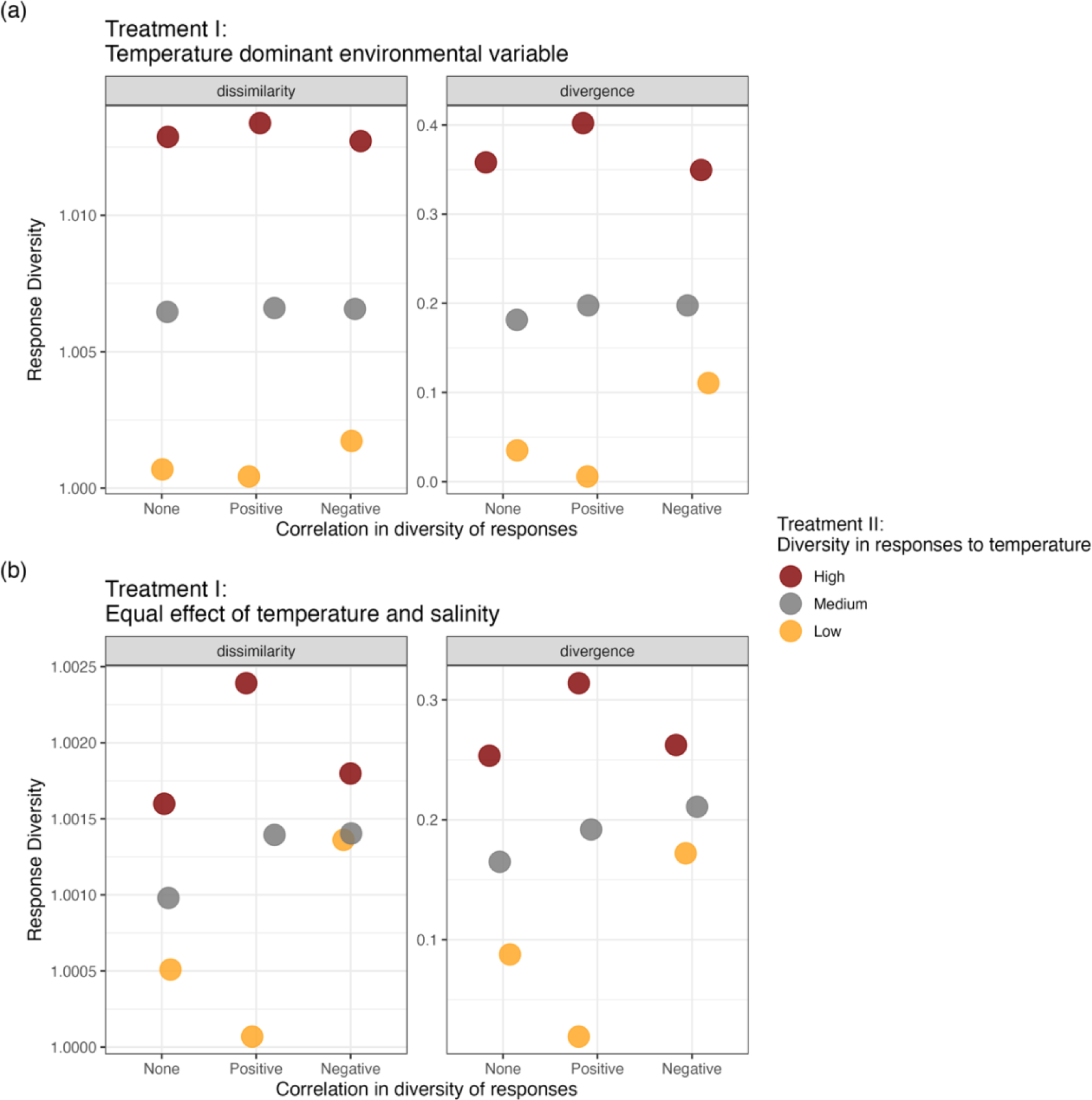
Effects of Factors I and II on potential response diversity. Potential response diversity measured as response dissimilarity and response divergence (see Ross et al. 2023) colour coded based on the different scenarios of diversity in species’ responses. (a) Shows potential response diversity for the case where temperature has a larger effect on species’ growth rate compared to salinity (Figs. 8a and 8b). (b) Shows potential response diversity for the case when temperature and salinity have the same effect on species’ growth rate (Figs. 8c and 8d). X axes show the three scenarios of factor II, that is the correlation between diversity in responses to temperature and salinity. Y axes show response diversity measured as dissimilarity (left side) and divergence (right side) as described by the facets.

Similarly, potential response divergence was highest in communities characterised by high diversity in species’ responses to temperature when temperature was the dominant environmental driver relative to salinity (Fig. 11a). The highest value of potential response divergence in this case emerged from the community with high diversity in optimum position for both temperature and salinity. Intuitively, the lowest potential response divergence value then emerged from the community with low diversity in optimum position for temperature and salinity. Potential response divergence was also high when the diversity in species’ responses to temperature was high, but diversity in responses to salinity was low (Fig. 11a).

When temperature and salinity had the same strength of effect on species’ growth rate, potential response divergence was highest when both diversity in species’ response to salinity and temperature were high (Fig. 11b). The second highest potential response divergence value was that for the community with high diversity in species responses to temperature and intermediate diversity in species responses to salinity. In contrast to the case where temperature was the dominant environmental driver, communities with high diversity in species’ responses to only one environmental driver had intermediate values of potential response divergence. Again, the lowest potential response divergence value was that for the community with low diversity in optimum growth rates for both temperature and salinity (Fig. 11b).

### 4.7. Effects of the treatments 1, 2, and 3 on response diversity

Here, we report the effects of the three treatments (treatment 1: relative driver importance; treatment 2: responses – driver correlation; treatment 3: temporal means of environmental variables) on response diversity measured when the direction of environmental change is known.

The mean value around which temperature and salinity fluctuated (treatment 3) had a large impact on response diversity, regardless of whether response diversity was measured as dissimilarity or divergence (Fig. 12). However, response dissimilarity and response divergence showed substantially different patterns (Figs. 12 and 13).

**Figure 12.**
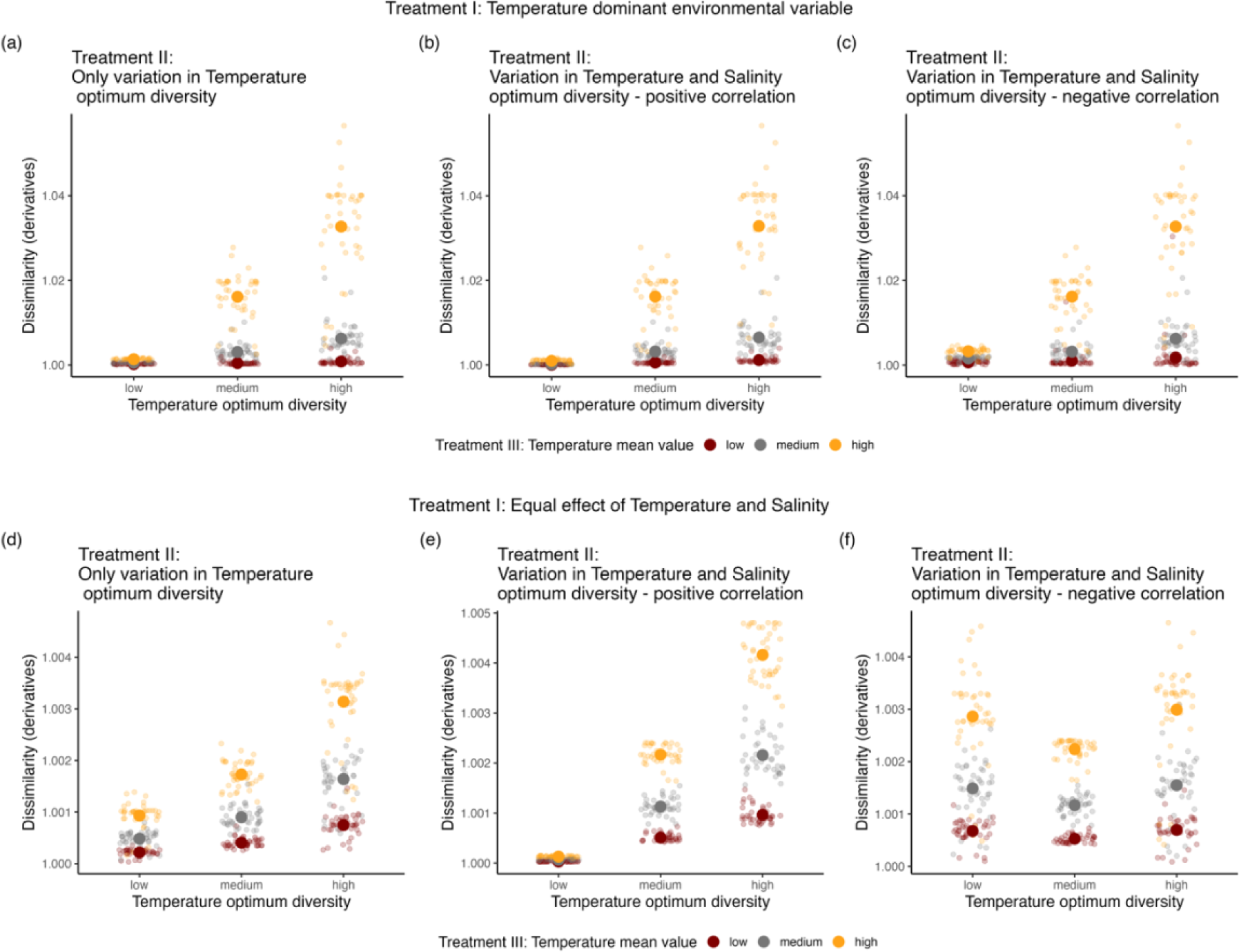
Effects of diversity in species’ responses on response diversity measured as dissimilarity. (a), (b), and (c) show how dissimilarity changes in the different scenarios of correlation between temperature and salinity optimum diversity depending on the mean value of the environment in the case where temperature is the dominant variable. (d), (e), and (f) show how dissimilarity changes in the different scenarios of correlation between temperature and salinity optimum diversity depending on the mean value of the environment in the case where temperature and salinity have an equal effect on species’ growth rate. For all panels, small dots represent the dissimilarity values measured for each community in our numerical experiments, whereas the big dots are the mean values for each of the treatment’s levels.

**Figure 13.**
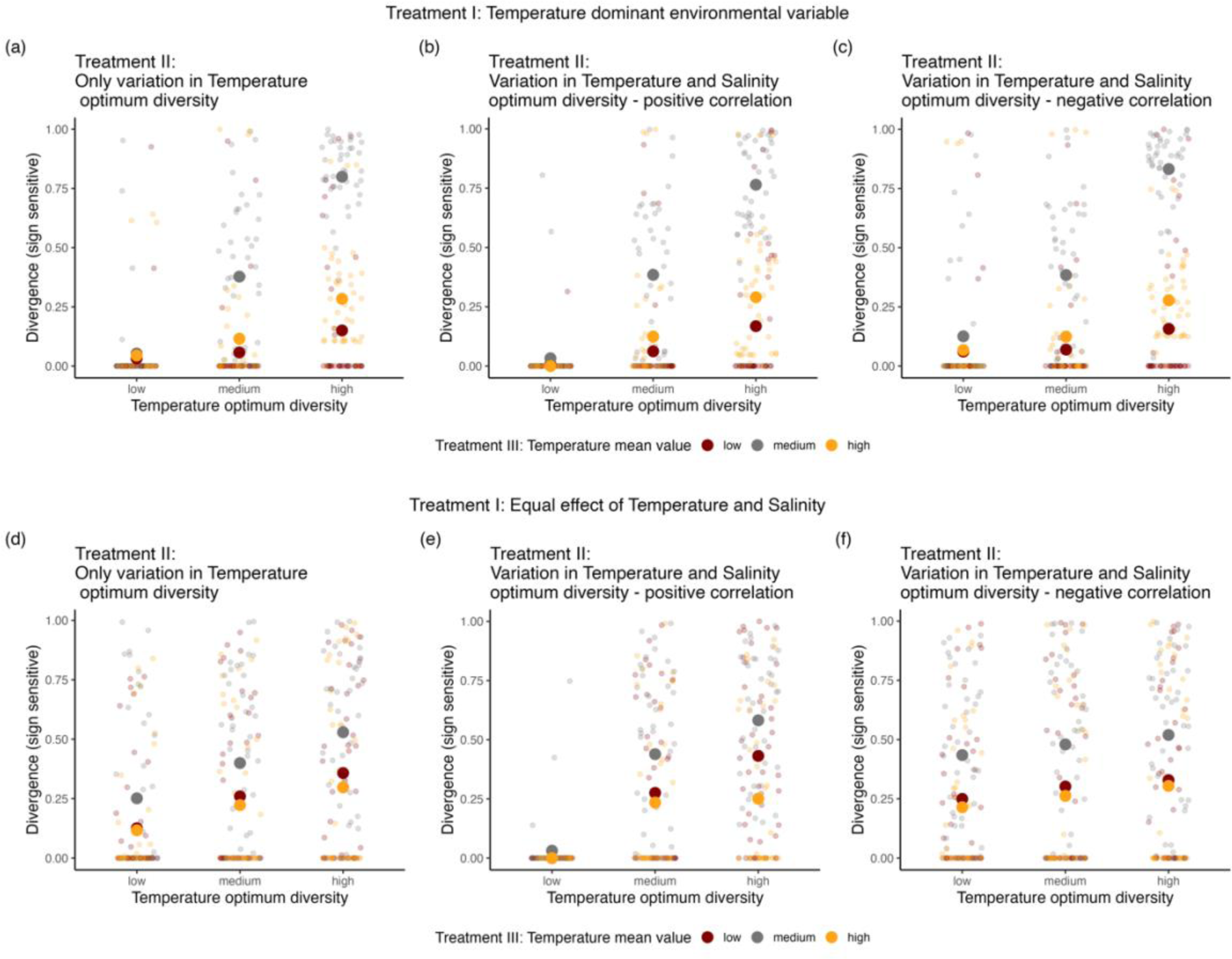
Effects of diversity in species’ responses on response diversity measured as divergence. (a), (b), and (c) show how divergence changes in the different scenarios of correlation between temperature and salinity optimum diversity depending on the mean value of the environment in the case where temperature is the dominant variable. (d), (e), and (f) show how divergence changes in the different scenarios of correlation between temperature and salinity optimum diversity depending on the mean value of the environment in the case where temperature and salinity have an equal effect on species’ growth rate. For all panels, small dots represent the dissimilarity values measured for each community in our numerical experiments, whereas the big dots are the mean values for each of the treatment’s levels.

Response dissimilarity increased with an increase in diversity of species responses to temperature when temperature fluctuated around intermediate and high mean values, when temperature was the dominant environmental driver. On the other hand, when temperature fluctuated around low mean values, response dissimilarity was always low when temperature dominated species’ responses (Fig. 12a - c).

Response dissimilarity was always highest for communities with high diversity in species responses to temperature when exposed to environmental conditions characterised by high mean values (yellow points in Fig 12). There was qualitatively no difference in response dissimilarity trends between the scenarios with different correlations between diversity in species responses to the environmental drivers when temperature was the dominant environmental driver (Figs. 12a–12c); that is, treatment 2 seemed to have little effect when species responses to temperature had a disproportionate influence of overall community responses to environmental conditions. When temperature and salinity had the same effect on species’ growth rate, response dissimilarity showed largely the same trends (Figs. 12d–12f).

One exception to this was under the negative correlation in diversity of species responses to the environmental variables. In this case we found similar response dissimilarity when the diversity of species’ responses to only one of the environmental variables was high, and the other low (Fig. 12f). That is, when temperature and salinity had an equal effect on species’ growth rate, treatment 2 (correlation between diversity in species’ responses to the environmental drivers) did have an effect, contrary to when temperature was the dominant environmental driver. In this case, having high diversity in response to either of the environmental driver led to similar level of response dissimilarity. We otherwise always found highest response dissimilarity when communities were exposed to temperature and salinity fluctuating around high mean values (yellow points in Figs. 12d–12f).

Response divergence also always increased with increasing diversity of species’ responses to temperature, but unlike for response dissimilarity, the highest values of response divergence were always associated with communities exposed to temperature and salinity fluctuating around intermediate, rather than high, mean values (grey points in Fig. 13). The scenario where species had low diversity in responses for both temperature and salinity showed the lowest divergence values, independent of the mean value of the environmental variables, and regardless of whether temperature was the dominant environmental driver or whether temperature and salinity had an equal effect (that is, regardless of treatment 1; Fig 13b and 13e).

When temperature was the dominant driver (treatment 1), response divergence showed the same pattern in all the correlation scenarios between diversity in responses to the environmental variables. That is, also for response divergence, Treatment 2 did not have an effect when temperature influenced disproportionately species’ growth rate compared to salinity. However, when temperature and salinity had an equal effect on species growth rate, treatment 2 had an effect. Indeed, as for response dissimilarity, when both environmental variables had an equal effect, having high diversity in responses to only one environmental variable was enough to produce high values of response divergence (Fig 13f). Conversely, when temperature was the dominant driver, communities with high diversity in responses to salinity, but low to temperature, had nonetheless low values of response divergence (Fig 13c).

## 5. Discussion

### 5.1. On the drivers of potential response diversity to multifarious environmental change (unknown direction of environmental change)

#### Treatment 1 & 2

We first investigated whether differences in the relative influence of environmental variables on species’ traits (treatment 1) and the correlation in the diversity of species responses to the drivers (treatment 2) can determine potential response diversity (unknown direction of environmental change). In this section, we only discuss the effects of the treatments on potential response diversity.

Generally, when temperature was the dominant stressor, response diversity was only driven by the diversity in species responses to temperature. This suggests that in case of driver dominance, response diversity largely behaves as it would to the single (dominant) driver (Ross *et al*. 2023). This is particularly relevant, as a recent meta-analysis found the net impacts of multiple environmental drivers are usually best explained by the effect of the stronger stressor alone in freshwater ecosystems (Morris et al. 2022). Hence, potential response diversity may be largely driven by the interspecific diversity in responses to a single, dominant, stressor in many real-world situations (Bray *et al*. 2019). Since potential response diversity can be seen as a measure of the insurance capacity of a community, in the case of a dominant environmental driver, interspecific diversity in responses to this single driver will largely determine the overall insurance potential in a community.

When no driver is dominant, then the correlation in diversity in species responses becomes vital in driving potential response diversity (treatment 2). Although driver dominance is common, at least in some systems (Bray *et al*. 2019; Morris *et al*. 2022), meta-analyses summarising multiple driver effects report that additive and synergistic interactions also occur (Birk *et al*. 2020; Crain *et al*. 2008; Jackson *et al*. 2016). Our analysis shows that when there is an equivalent effect of the environmental variables on species’ responses, perhaps unsurprisingly, the highest potential response diversity in this case was observed for communities with high diversity in responses to both environmental variables. This result confirmed the soundness of our new method for calculating potential response diversity, which responded meaningfully to changes in the amount of diversity in species’ responses. Having a reliable method to calculate potential response diversity and for understanding the underpinning drivers (treatments 1 and 2) of this important aspect of response diversity will allow estimation of the insurance capacity of a system when the direction of the environmental change is unknown. This new methodology for calculating potential response diversity, together with a deeper understanding of the underlying drivers, should provide ecosystem managers with a new to tool to assess and protect the stable delivery of ecosystem functions and services (Elmqvist *et al*. 2003; Folke *et al*. 2004).

We mostly focused on how to calculate potential response diversity using species’ entire performance surfaces. However, in many cases, we have some expectations regarding environmental change corresponding to a scenario (e.g. 2 degrees warming, higher nutrient load), rather than a totally unknown trajectory of future environmental change (IPCC 2019). A given environmental change scenario would correspond to a particular area on the performance surface, and potential response diversity can be calculated only for that area. This allows predicting how well a community may be “insured” against environmental change as represented under that specific environmental change scenario. This approach should therefore facilitate specific predictions about response diversity and, in turn, ecological stability in response to anticipated future environmental change scenarios.

Overall, we highlight here that potential response diversity is interdependent on the diversity in species responses to single environmental variables (treatment 2), and on the relative magnitude of response species show to each environmental variable (treatment 1).

### 5.2. On the drivers of response diversity to multifarious environmental change (known direction of environmental change)

#### Treatment 1 and 2

Here, we only discuss the effects of the treatments on response diversity measured when the direction of environmental change is known.

Our work based on simulated species responses to multifarious environmental change suggests that the diversity of species’ responses to each environmental variable individually is an important driver of potential response diversity (treatment 2). Critically, though, these differences in multifarious response diversity depend on whether there is a dominant environmental driver that has a disproportionate effect on species’ responses compared to the other(s) (treatment 1).

When species’ responses are more strongly influenced by one environmental variable compared to another, having high diversity in responses to the dominant environmental variable is a necessary condition to create high response dissimilarity or response divergence. Conversely, a community characterised by high diversity in responses to the subdominant driver (salinity), but low diversity in responses to the dominant driver (temperature), will always result in low response dissimilarity (Figs. 12a–12c) or response divergence (Figs. 13a–13c). Indeed, when temperature was the dominant environmental driver, response dissimilarity and response divergence were highest in communities with high diversity in responses to temperature (i.e. characterised by high diversity in temperature optimum diversity, Figs. 12a–12c and Figs. 13a–13c) but were low when only diversity for salinity was high. These results largely mirrored those for potential response diversity, suggesting that driver dominance (or its absence) is likely to determine all aspects of response diversity.

In contrast, when salinity and temperature had similar effects on species growth rate, either diversity in responses to salinity or to temperature could produce high response dissimilarity (Fig 12f) or response divergence (Fig. 13f). Hence, the effects of manipulating treatment 2 (correlation between diversity in species responses to the environmental drivers) depend directly on treatment 1 (relative driver importance).

#### Treatment 3

Whether response diversity changes depending on features of the environmental change drivers against which response diversity is measured is probably one of the least considered aspects of the already understudied and underappreciated sphere of response diversity (Laliberté *et al*. 2010; Mori *et al*. 2013).

To gain insight into the effects of the temporal means of the environmental variables on response diversity, we simulated three different conditions where the environmental variables fluctuate around low, intermediate, or high mean values. We anticipated highest response diversity when temperature and salinity fluctuate around an intermediate mean value in communities with high diversity in species’ responses to both environmental variables. This is because, when a community contains species with a high diversity of responses to both environmental variables, and temperature and salinity fluctuate around intermediate values, then some species will have their maximum growth rate at higher values (compared to the mean value around which the environment fluctuates), and other species’ maximum growth rates will fall at lower values. This situation should allow the highest possible response diversity among all scenarios, as there will be species performing extremely well, other extremely poorly, and others neither very well nor very poorly (Fig. 9).

However, this expectation was not met for the response dissimilarity. We found that response dissimilarity was consistently higher when temperature and salinity were fluctuating around high mean values, independently of whether temperature was the dominant environmental driver or not (treatment 1). In a previous study, Laliberté *et al*. (2010) found some rare, but drastic increases in response diversity at the high end of a land-use intensity gradient. At least in our study, this counterintuitive result is explained by the shape of species’ performance curves. Indeed, species’ performance curves showed the largest magnitude of change in growth rate in response to either temperature or salinity at the higher end of the environmental gradients (Figs. 8a and 8b). Thus, when diversity in species’ responses increases, even a less negative growth rate in the high end of the gradient for one of the two environmental variables results in a less negative directional derivative, which causes an increase in the value of response dissimilarity. Note that response dissimilarity does not consider the direction of the variation in the species’ environmental responses, only the absolute variation among responses (Ross *et al*. 2023). Hence, even if most of the responses take the same direction (negative growth rates, and thus negative derivatives), but one or few species in a community exhibit less negative growth with an environmental driver, this variation in responses is sufficient to produce a high response dissimilarity value.

These results force us to consider the importance of including the direction (positive or negative) of responses to the environmental variables when calculating response diversity. For example, Winfree & Kremen (2008) found that changes in bee species’ abundance with increasing proportions of native vegetation in the surrounding landscape were positively correlated in northern central California. Even though all species showed negative associations with native vegetation loss, the associations differed in degree, leading to response diversity. In such a case, even at the extreme of the gradient of native vegetation loss, we may find high response diversity, because species respond in the same direction (all declining in abundance), but some less strongly than others. When species responses only differ in the degree of their responses, but not in the direction, ecosystem functioning may not complete disappear (Winfree & Kremen 2008), but they may decline in their absolute value. That is, their temporal stability may be compromised. For instance, Hermann *et al*. (2023) found that all freshwater zooplankton species in an experimental community responded negatively to the exposure to a fungicide, but some less so than others. This led to a decline of total zooplankton abundance through time compared to unperturbed communities (Hermann *et al*. 2023). This reduction in total zooplankton abundance increased the temporal variability of the community aggregate property (total abundance), despite some difference in absolute species responses to the perturbation. Thus, response dissimilarity, or any other way of quantifying response diversity that only accounts for differences in the absolute species’ responses without considering the direction of these responses, can only inform about whether a community or ecosystem properties might be preserved, but is less informative about whether the temporal stability of these properties could be promoted. We suggest that if response diversity is to be understood and implemented to promote temporal stability of community aggregate properties and of ecosystem services, it is fundamental to consider not only the absolute difference in species’ responses, but also the direction of these responses.

Response divergence, on the other hand, met our predictions, and was highest when the environmental drivers fluctuated around intermediate mean values. When the environment was fluctuating around more extreme values, independently of whether it was doing so at the lower or higher end of an environmental gradient, response divergence was always lower than when environments fluctuated around intermediate values (Fig. 13). Previous studies have concluded that environmental change can reduce both functional redundancy and response diversity because of species loss (Laliberté *et al*. 2010; Mori *et al*. 2013). Our study shows that even without losing species (we did not simulate any species extinctions), the values of response diversity measured as response divergence, strongly depend on the value of the environmental drivers. This appears to be the case independently of the correlation in diversity in species’ responses to environmental drivers. Indeed, even when interspecific diversity in species responses to both environmental drivers was high, drivers fluctuating around high or low mean value determined a reduction in response divergence compared to when the environment fluctuated around intermediate values.

Overall, response divergence emerged as a somewhat more intuitive measure of response diversity compared to response dissimilarity; response divergence exhibited expected and consistent patterns in our simulations to diversity in species responses and to the variation in mean environmental values, without a strong dependence on the shape of species’ performance curves. Additionally, response divergence intrinsically considers the direction and the magnitude of the difference in species’ responses to environmental change (Ross *et al*. 2023). It is therefore likely to be the more appropriate of the two metrics considered here for quantifying response diversity if one wants to study the relationship between temporal stability of aggregate community and ecosystem properties and response diversity.

Finally, Mori *et al*. (2013) suggested that if aiming to conserve an ecosystem’s ability to deliver functions and services, the specific relationship between response diversity and anthropogenically driven environmental change is substantially more important than just measuring response diversity under unperturbed conditions. Here, we take a first step toward unveiling the mechanistic basis of the relationship linking response diversity to environmental change. We not only explore this relationship in general terms, but specifically in a multifarious environmental change context, which is widely recognised as the norm in real-world systems in the Anthropocene (Bowler *et al*. 2020).

## 6. Conclusions

Here, we proposed an empirically tractable method for quantifying response diversity in the context of multifarious environmental change that can be easily applied to experimental as well as observational studies. We showed how response diversity measured when multiple environmental drivers are simultaneously changing depends on the direction of the environmental change. Moreover, we proposed a way to calculate the potential response diversity of a system when the direction of the environmental change is unknown— that is, an estimate of the insurance capacity of a community under any possible environmental change scenario. Finally, we explored the drivers of response diversity in a multifarious environmental change context. In this regard, we showed how response diversity is determined by the diversity in species’ responses to each environmental driver, as well as by the correlation in species responses to these drivers. We also found that response diversity strongly depends on the mean value around which the environment fluctuates, and on the effects of each environmental variable on species’ traits. In sum, the framework we proposed herein makes an important step towards understanding the insurance capacity of ecological communities in the Anthropocene in the face of unprecedented and multifarious environmental change.

## 7. Glossary

**Directional (first) derivative:** please first read the Glossary entry for *Partial derivative*, since the directional derivative generalises the concept of a partial directional derivative. The directional derivative of a function is like a partial derivative of a function, except that it is the derivative in any specified direction of change in the variables of the function. For example, for the function *f(x*,*y*) one could measure the derivative in the direction of equal change in x and y (thus at 45 degrees to a partial derivative).

**Direction of environmental change:** When there are multiple environmental drivers changed, for example two, there can be different rates of change in these. These rates of change dictate the direction of environmental change. The direction can be encoded in a *unit vector* (see Glossary).

**Environmental driver**: any abiotic factor that causes measurable changes in properties of biological communities (Millennium Ecosystem Assessment 2005; Carpenter *et al*. 2006).

**Environmental location**: a location on a species performance surface, given by a specific value of each environmental driver. E.g., 20 Celsius and 0.25 ppt salinity give a location on a temperature – salinity surface.

**First derivative:** the first order derivative of a function represents the rate of change of variable with respect to another variable (i.e. the change in a species’ growth rate as temperature increases).

**Generalised additive model (GAM)**: a flexible regression model that allows for non-linear relationships between the dependent variable and independent variables (Hastie & Tibshirani 1987).

**Partial derivative:** a partial derivative of a function of multiple variables is the rate of change of the value of a function (i.e., the first derivative or slope) with respect to variation in only one of the dependent variables. For the function *f(x*,*y*) the partial derivative with respect to x would be the derivative for variation in *x* but no variation in *y*.

**Potential response diversity:** a way to measure response diversity when the direction of environmental change is unknown. Potential response diversity captures the insurance capacity of a system under all possible environmental conditions. Potential response diversity can be seen as a general and comprehensive quantification of response diversity that does not depend on the direction of the environmental change.

**Response diversity:** the variety of responses to the environment exhibited by different species in a community (Elmqvist *et al*. 2003; Mori *et al*. 2013).

**Response dissimilarity**: a response diversity metric calculated based on pairwise Euclidean distances (i.e. dissimilarity) in performance environment relationships between all pairs of species in the community (see Leinster & Cobbold 2012). The metric captures the absolute dissimilarity in species responses to the environment but does not consider the sign of the dissimilarity (it is sign-insensitive).

**Response divergence**: a response diversity metric that accounts for whether performance– environment responses diverge in direction (it is sign-sensitive; Ross *et al*. 2023). Higher values represent a greater balance of positive and negative responses around zero (where zero indicates no relationship between performance and an environmental driver).

**Species performance curve:** a measure of some aspect of a population’s *performance*, such as its intrinsic rate of increase or biomass change along an environmental gradient (Ross et al. 2023). Note that “curve” implies only one dimension of environmental change, e.g., change in only temperature.

**Species performance surface:** the same as a species performance curve, except with two or more dimensions of environmental change (e.g., salinity *and* temperature).

**Unit vector**: A vector of length 1. For example, these are unit vectors: [0, 1], [1, 0], [1/√2,1/√2]. They are used for encoding information about direction of change in environmental variables.

## Acknowledgments

S.R.P-J.R. was supported by subsidy funding to the Okinawa Institute of Science and Technology Graduate University (OIST). R.L. was supported by the Swiss National Science Foundation (SNF) Project 310030_188431.

## 8. Author contributions

FP: Conceptualisation, Formal analysis, Investigation, Methodology, Writing - original draft, Project administration, Software, Writing - review & editing.

RL: Providing original data, Writing - review & editing.

FP: Conceptualisation, Supervision, Methodology, Writing - review & editing.

SRP-JR: Conceptualisation, Writing - review & editing.

GLS: Software, Writing - review & editing

OLP: Conceptualisation, Formal analysis, Funding acquisition, Investigation, Methodology, Project administration, Resources, Software, Supervision, Writing - review & editing.

## Notes

https://github.com/opetchey/multifarious_response_diversity/releases/tag/v0.1

### Competing Interest Statement

The authors have declared no competing interest.

