## Appendix1 for "Response diversity in the context of multifarious environmental change"

1 Introduction
2 The principle
3 A simulated empirical example
4 Partial derivatives
5 Directional derivatives
6 Response diversity calculation
7 Empirical example
8 Partial derivatives for a single species
9 Directional deriviatives

Code

### Appendix for: Response diversity in the context of multifarious environmental change.

Francesco Polazzo, Romana Limberger, Frank Pennekamp, Samuel Ross, Gavin Simpson, Owen Petchey

11 December, 2023

### 1 Introduction

Researchers have previous suggested that the response diversity of a community be measured by the diversity of responses to environmental change. For example, one can measure the response of each of the species' intrinsic growth rate to temperature, quantify the strength and direction of these responses (e.g., as the first derivative of the response curve), and calculate the diversity of responses (e.g., by calculating variation in the first derivatives among the species in a community). When responses are nonlinear, the response diversity will be a function of the environmental state (i.e. the first derivative is a function of the value of the environmental state). So far we demonstrated this approach for quantifying response diversity in the context of a single environmental factor, but given that multiple environmental factors can change simultaneously, we need an approach that works in that context.

### 2 The principle

*To learn about the mathematical principles watch these youtube videos:*

- Surfaces and Partial Derivatives (https://www.youtube.com/watch?v=k4wNIZr8GU4)
- 54. Slope of the Surface in Any Direction - Directional Derivative, and Properties of the Gradient (https://www.youtube.com/watch?v=wfjipWmyRYg)

Imagine that the growth rate of a population depends on two environmental factors, e.g. temperature and salinity. We can represent the dependency as $G = f(T, S)$, where $G$ is growth rate, $T$ is temperature, and $S$ is salinity. It may be that the dependencies are linear, nonlinear, and with an interaction between temperature and salinity, hence our approach needs to be able to accommodate this phenomena.

The response of growth rate to change in temperature and salinity is the gradient / slope of this surface, with units of growth rate [per time] per temperature [degrees C] per salinity [parts per thousand]. Because the slope (first derivative) of the surface can (when dependencies are nonlinear) vary across the surface (location on the surface), and can vary in different directions on the surface, to calculate a slope we must specify the current environment (location on the surface) and the direction of change in the environment. The location on the curve is the current environmental condition, $(T_0, S_0)$, and the direction of environmental change is the unit vector $\hat{u} = \langle U_T, U_S \rangle$.

Put another way, we calculate a directional derivative at a point on the response surface. We can write this as $D_{\hat{u}} f(T_0, S_0)$ and can calculate it as $f_T(T_0, S_0)U_T + f_S(T_0, S_0)U_S$, where $f_T$ is the partial derivative of $f(T, S)$ with respect to $T$ and $f_S$ is the partial derivative of $f(T, S)$ with respect to $S$.

Efficient evaluating in $n$ dimensions can be done by taking the dot product of the partial derivatives at the location and the direction unit vector: $D_{\hat{u}} f(T_0, S_0) = \triangledown f \cdot \hat{u}$ where, $\triangledown f = \langle f_T, f_S \rangle$. (In R, the dot product of `a` and `b` is `sum(a*b)`)

Figure 2.1 is an illustration of the principle of directional derivatives on a surface.

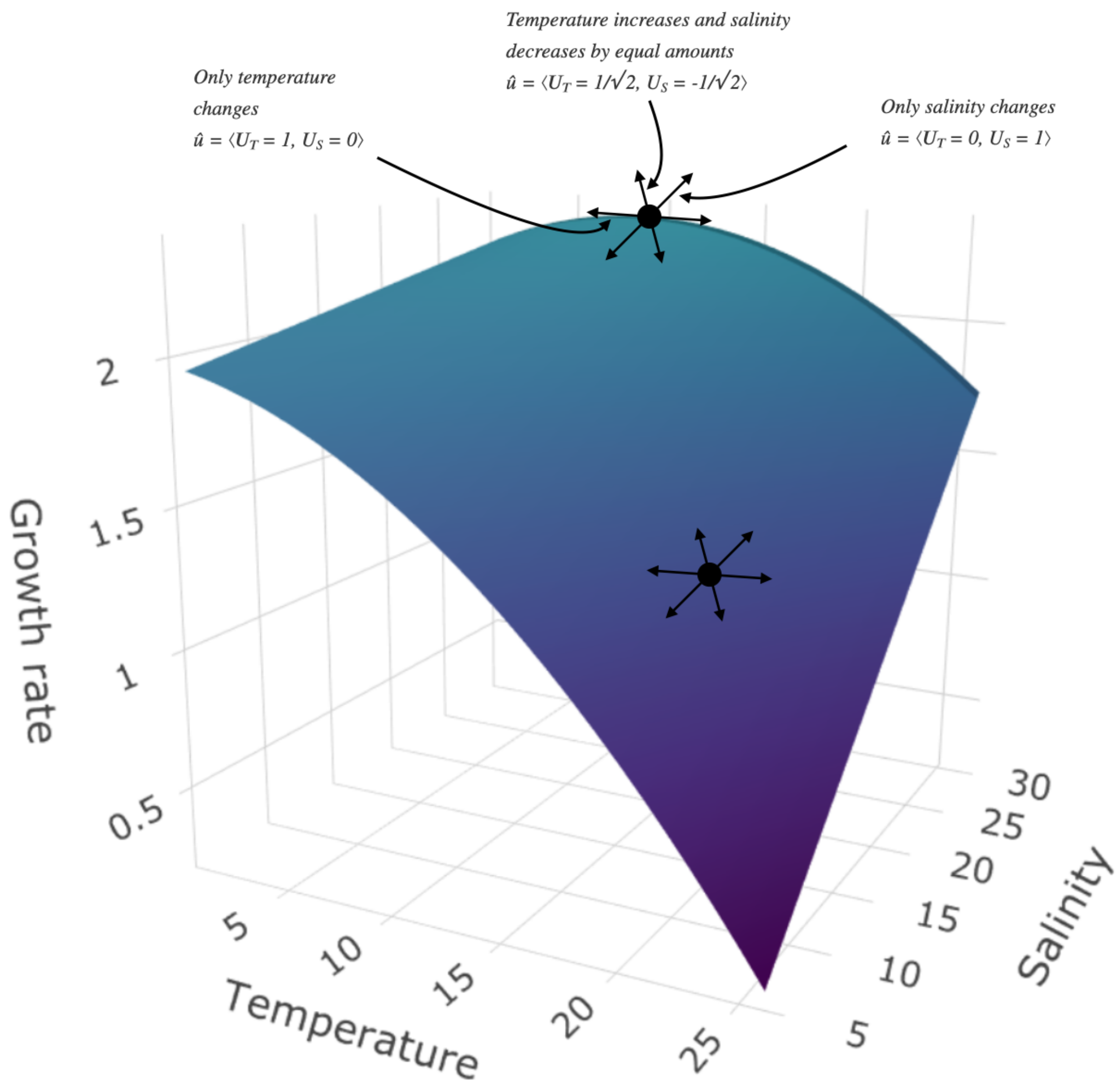


Figure 2.1: This figure needs considerable improvement! It is currently a keynote illustration.

### 3 A simulated empirical example

Numerous mathematical functions have been used to represent how organismal performance changes with an environmental driver citation require (). Moreover, multiple mathematical functions have been used to represent an interactive effect of two or more environmental drivers on species performance e.g. Thomas et al 2017 (https://onlinelibrary.wiley.com/doi/full/10.1111/gcb.13641). *We have to decide if we are going to make simulations with different of these functions. This might increase our confidence that the method for calculating response diversity is robust to variation in types of response curve. If we decide not to, then we should likey argue that we think its robust.*

#### 3.1 Simulating performance curves

Let us use the Eppley performance curve, which was used, for example, in this paper Bernhardt et al. 2018 (https://royalsocietypublishing.org/doi/10.1098/rspb.2018.1076).

With one environmental variable, the performance (i.e., rate) is given by:

- $\mathrm{rate(E)} = \mathrm{ae}^{\mathrm{bE}}(1 - (\frac{\mathrm{E-z}}{\mathrm{w/2}})^2)$
- $\mathrm{E}$ is the values of the environmental factor.
- $\mathrm{z}$ controls location of maximum.
- $\mathrm{w}$ controls range of $\mathrm{E}$ over which the rate is positive.
- $\mathrm{a}$ scaling constant.
- $\mathrm{b}$ controls rate of increase towards the maximum rate, as $\mathrm{E}$ increases.

Adding a second environmental variable gives:

$\text{rate}(E_1, E_2) = a_1 e^{b_1 E_1}(1 - (\frac{E_1 - z_1}{w_1/2})^2) + a_2 e^{b_2 E_2}(1 - (\frac{E_2 - z_2}{w_2/2})^2)$

In this case, it is clear the effect of $E_1$ and $E_2$ is defined as being additive. For example, the value of $E_2$ does not affect the value of $E_1$ at which the rate is maximised ($z_1$), and vice-versa (see also Figure **??**)

*Including an interaction*. One way to do this is to make the value of $E_1$ at which the rate is maximised depend on the value of $E_2$:

$\text{rate}(E_1, E_2) = a_1 e^{b_1 E_1}(1 - (\frac{(E_1 + z_{int21} * E_2 - z_1)}{w_1/2})^2 + a_2 e^{b_2 E_2}(1 - (\frac{E_2 - z_2}{w_2/2})^2$

When $z_{int21} = 0$ then this equation becomes the previously mentioned additive one. When $z_{int} \neq 0$ then the value of $E_1$ at which the rate is maximised is a function of the value of $E_2$. We used this method for adding an interaction due to its simplicity. Other methods could be used, and if also or otherwise used could add confidence about the robustness of the method for calculating response diversity.

### 3.2 Simulating multiple species' performance curves

#### 3.2.1 No interacting environmental effects

First we create (or import) a table of parameter values of each species, with species in the rows and parameters in the columns. In the following example, only values of the $z$ parameters differ among the species (which determine the location of the maximum rate).

**Show** 10 **entries** **Search:**

| | a1 | b1 | z1 | w1 | a2 | b2 | z2 | w2 | z_int21 | sd_rate |
|---|---|---|---|---|---|---|---|---|---|---|
| | A | A | All | A | A | A | All | A | Al | All |
| 1 | 1e-9 | 0.063 | 270.084014022723 | 60 | 0.001 | 0.02 | 22.434567604214 | 10 | 0 | 0 |
| 2 | 1e-9 | 0.063 | 277.091162335128 | 60 | 0.001 | 0.02 | 16.4699217258021 | 10 | 0 | 0 |
| 3 | 1e-9 | 0.063 | 290.880527682602 | 60 | 0.001 | 0.02 | 10.0129270693287 | 10 | 0 | 0 |
| 4 | 1e-9 | 0.063 | 286.016520692501 | 60 | 0.001 | 0.02 | 24.2431184416637 | 10 | 0 | 0 |
| 5 | 1e-9 | 0.063 | 294.497439691331 | 60 | 0.001 | 0.02 | 16.1726777348667 | 10 | 0 | 0 |
| 6 | 1e-9 | 0.063 | 291.006739914883 | 60 | 0.001 | 0.02 | 25.7363655837253 | 10 | 0 | 0 |
| 7 | 1e-9 | 0.063 | 281.938274691347 | 60 | 0.001 | 0.02 | 26.7779199965298 | 10 | 0 | 0 |
| 8 | 1e-9 | 0.063 | 294.50001725927 | 60 | 0.001 | 0.02 | 15.4671661788598 | 10 | 0 | 0 |
| 9 | 1e-9 | 0.063 | 296.871775598265 | 60 | 0.001 | 0.02 | 23.1785696651787 | 10 | 0 | 0 |
| 10 | 1e-9 | 0.063 | 272.563425118569 | 60 | 0.001 | 0.02 | 20.3631447255611 | 10 | 0 | 0 |

Showing 1 to 10 of 10 entries Previous 1 Next

For convenience we then convert the table of parameters into a list-column (https://dcl-prog.stanford.edu/list-columns.html). We can then easily make performance curves of each of the species, and put those into a list-column in the same table.

Code

Here are some examples of the species' performance curves (only with additive effects of $E_1$ and $E_2$).

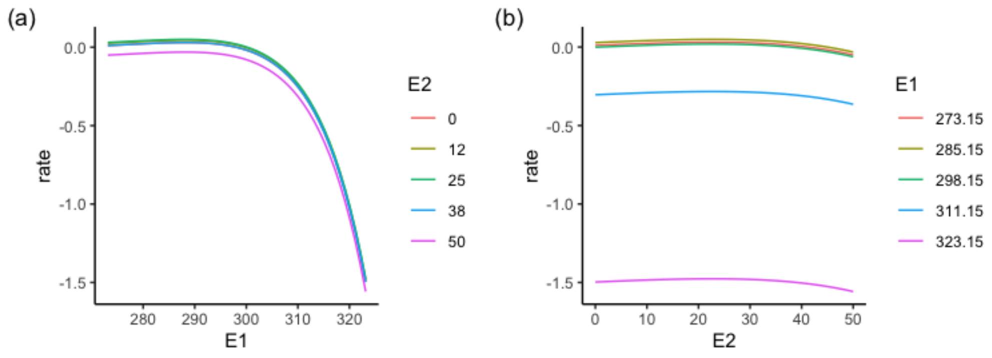


Figure 3.1: Performance curves for a species with maximum growth at **low** values of $E_1$. Without interacting environmental effects.

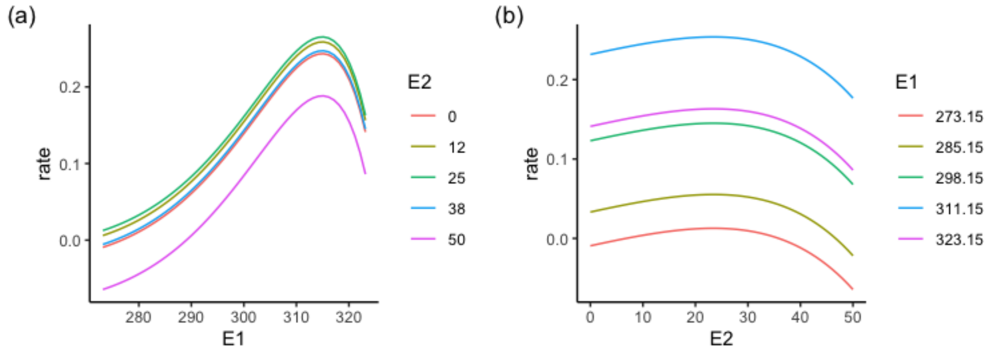


Figure 3.2: Performance curves for a species with maximum growth at **high** values of $E_1$. Without interacting environmental effects.

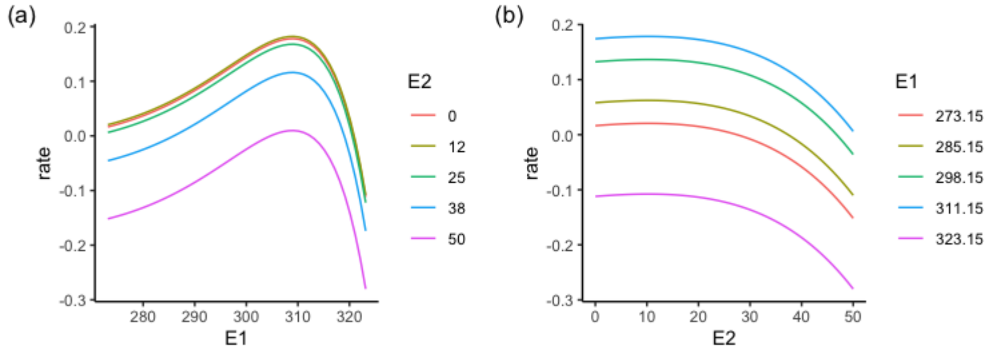


Figure 3.3: Performance curves for a species with maximum growth at **low** values of $E_2$. Without interacting environmental effects.

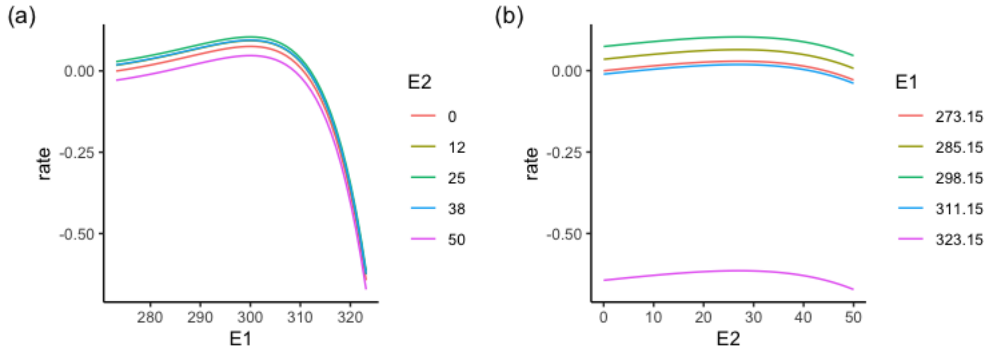


Figure 3.4: Performance curves for a species with maximum growth at **high** values of $E_2$. Without interacting environmental effects.

#### 3.2.2 Interacting environmental effects

And now with interacting environmental effects…

**Show** 10 **entries** **Search:**

| | a1 | b1 | z1 | w1 | a2 | b2 | z2 | w2 | z_int21 | sd_rate |
|---|---|---|---|---|---|---|---|---|---|---|
| | | | All | | | | All | | All | A |
| 1 | 1e-9 | 0.063 | 288.354849403258 | 60 | 0.001 | 0.02 | 27.06988717895 | 10 | -0.201642219638534 | 0 |
| 2 | 1e-9 | 0.063 | 271.357648316771 | 60 | 0.001 | 0.02 | 12.6090735010803 | 10 | -0.228507629167015 | 0 |
| 3 | 1e-9 | 0.063 | 286.300132074393 | 60 | 0.001 | 0.02 | 21.7647998407483 | 10 | -0.217486977906434 | 0 |
| 4 | 1e-9 | 0.063 | 272.106599032413 | 60 | 0.001 | 0.02 | 21.3384760916233 | 10 | -0.1396200451061 | 0 |
| 5 | 1e-9 | 0.063 | 284.935023745056 | 60 | 0.001 | 0.02 | 25.3184265876189 | 10 | -0.18356941957818 | 0 |
| 6 | 1e-9 | 0.063 | 279.786555566825 | 60 | 0.001 | 0.02 | 27.2984294313937 | 10 | -0.180088470986822 | 0 |
| 7 | 1e-9 | 0.063 | 295.299732657149 | 60 | 0.001 | 0.02 | 11.0968862567097 | 10 | -0.210827960997092 | 0 |
| 8 | 1e-9 | 0.063 | 288.789769064169 | 60 | 0.001 | 0.02 | 13.9972907165065 | 10 | -0.229017128062291 | 0 |

| | a1 | b1 | z1 | w1 | a2 | b2 | z2 | w2 | z_int21 | sd_rate |
|---|---|---|---|---|---|---|---|---|---|---|
| 9 | 1e-9 | 0.063 | 274.479032030795 | 60 | 0.001 | 0.02 | 27.274556402117 | 10 | -0.242164493335918 | 0 |
| 10 | 1e-9 | 0.063 | 280.406717893202 | 60 | 0.001 | 0.02 | 18.3316139038652 | 10 | -0.180328044210025 | 0 |

Showing 1 to 10 of 10 entries Previous 1 Next

For convenience we then convert the table of parameters into a list-column (https://dcl-prog.stanford.edu/list-columns.html). We can then easily make performance curves of each of the species, and put those into a list-column in the same table.

Code

Here are some examples of the species' performance curves (with interacting effects of $\mathrm{E}_1$ and $\mathrm{E}_2$).

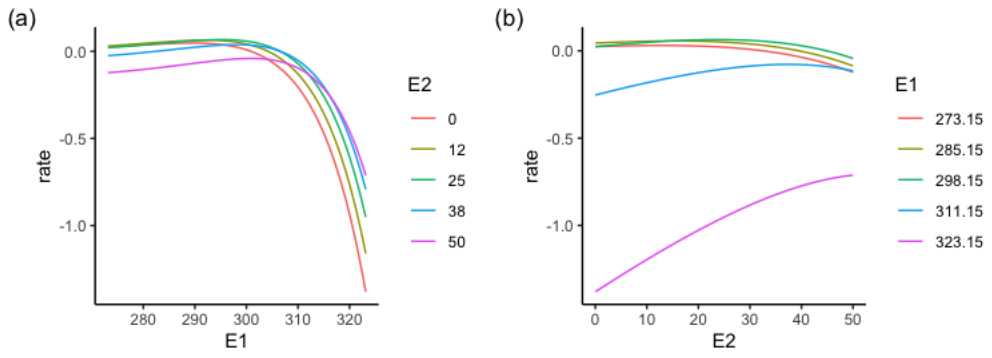


Figure 3.5: Performance curves for a species with maximum growth at **low** values of $\mathrm{E}_1$. With interacting environmental effects.

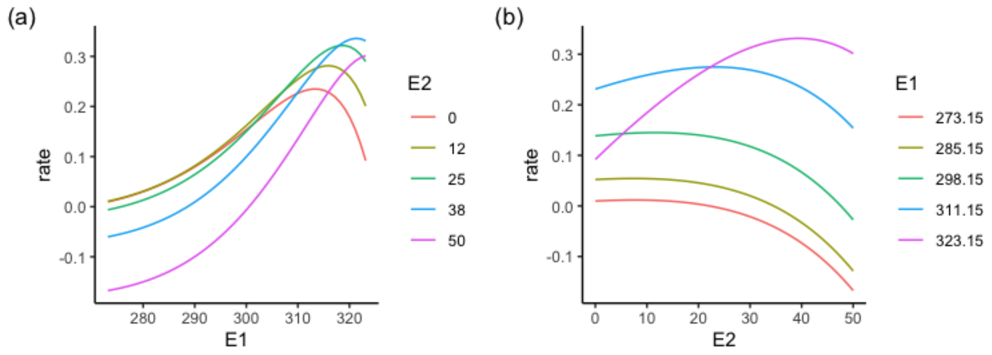


Figure 3.6: Performance curves for a species with maximum growth at **high** values of $\mathrm{E}_1$. With interacting environmental effects.

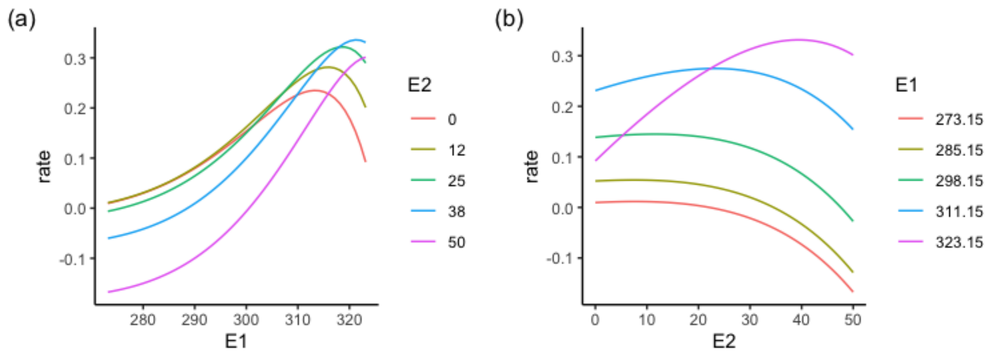


Figure 3.7: Performance curves for a species with maximum growth at **low** values of $\mathrm{E}_2$. With interacting environmental effects.

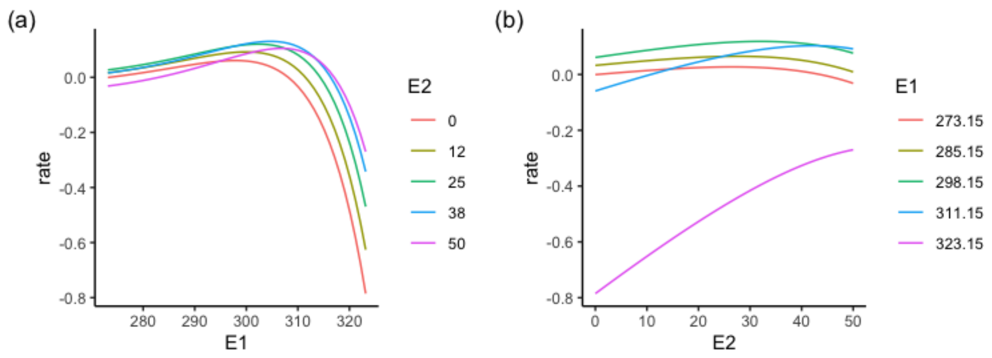


Figure 3.8: Performance curves for a species with maximum growth at **high** values of $E_2$. With interacting environmental effects.

### 3.3 Fitting GAMs to noisy rate observations

Try with and without an interaction. Therefore make two species, one with no interaction `z_int = 0` and the other with `z_int = 0.1` . All other parameters are the same. Note that noise is added to the rate observations.

Bottom line is that the gam picks up an interaction when we have included one in the parameters used to generate the rates, and does not pick one up when we have not. This confirms that our more mechanistic thinking and methods are matching our statistical thinking and methods, and confirms that each are promising, so far.

GAM notes:

- Owen watched this youtube video Introduction to Generalized Additive Models with R and mgcv (https://youtu.be/sgw4cu8hrZM) by Gavin Simpson (author of the `gratia` package that we've been using to calculate derivatives of GAMs.). By the way, there is a part of this video about Model Selection, about Confidence Intervals, and about p-values for smooths (about 1h:57m to 2h:26) which I think is less useful for our purposes.
- In the video mentioned above at about 54m:30s (https://youtu.be/sgw4cu8hrZM?t=2730), there is the recommendation to estimate the penalisation parameter ( $\lambda$) via restricted maximum likelihood, which is done by giving the the argument `method = "REML` in the `gam()` call (the default in `mgcv` version 1.8-36 is `method="GCV.Cp"` ).
- Default smoother is a *low rank thin plate spline* ( `bs = "tp"` )
- `s(E1, E2) will fit a *bivariate isotropic thin plate spline*. *Isoptropic* means there is a single smoothness parameter for the smooth. It is sensitive to the scale of $E1$ and $E2$.
- Tensor products ( `te(E1, E2` ) have separate marginal basis, and therefore separate smoothness parameters. They are invariant to scales of $E1$ and $E2$. Therefore we should be using tensor products.
- Specifying a model with `te(E1, E2` ) does not then allow inspection of the main effects and interaction term–there is essentially only one smoother which contains the main effects and interaction. So it is difficult to know if the interaction term is required. An alternative is to specify the model as `s(E1) + s(E2) + ti(E1, E2)` , in which the third term is then only the interaction part and allows a decomposition into the different effects. Useful for examining if the interaction is required.
- To use cubic splines use `bs = "cr"` .
- There are many smoothers in `mgcv`
- Choosing *k* (the *basis complexity*, which is the maximum wiggliness) is a bit of an art. The penalty then shrinks the used basis complexity, reducing the effective degrees of freedom. Must check that large enough *k* is provided; use `gam.check()` . Only cost to large $k$ is computational effort.

**Show** 10 **entries** **Search:**

| | a1 | b1 | z1 | w1 | a2 | b2 | z2 | w2 | z_int21 | sd_rate |
|---|---|---|---|---|---|---|---|---|---|---|
| | All | All | All | All | All | All | All | All | All | All |
| 1 | 1e-9 | 0.063 | 285 | 60 | 0.001 | 0.02 | 20 | 10 | 0 | 0.02 |
| 2 | 1e-9 | 0.063 | 285 | 60 | 0.001 | 0.02 | 20 | 10 | -0.2 | 0.02 |

Showing 1 to 2 of 2 entries Previous 1 Next

#### 3.3.1 Without interaction

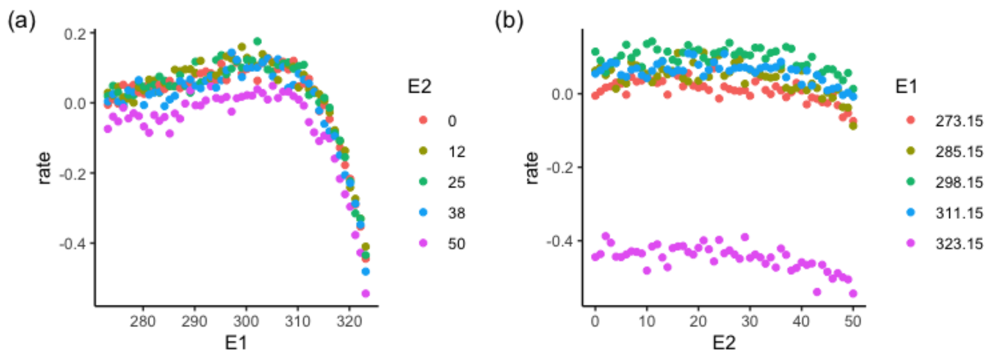


Figure 3.9: Performance curves for a species *without* interacting environmental effects and with some noise in the rate.

```
##
## Family: gaussian
## Link function: identity
##
## Formula:
## rate ~ ti(E1) + ti(E2) + te(E1, E2)
##
## Parametric coefficients:
##             Estimate Std. Error t value Pr(>|t|)
## (Intercept) 0.015428   0.000499   30.92   <2e-16 ***
## ---
## Signif. codes:  0 '***' 0.001 '**' 0.01 '*' 0.05 '.' 0.1 ' ' 1
##
## Approximate significance of smooth terms:
##                edf Ref.df       F p-value
## ti(E1)    3.999310  4.000 14148.5  <2e-16 ***
## ti(E2)    3.936725  3.997   608.8  <2e-16 ***
## te(E1,E2) 0.001897 16.000     0.0   0.992
## ---
## Signif. codes:  0 '***' 0.001 '**' 0.01 '*' 0.05 '.' 0.1 ' ' 1
##
## R-sq.(adj) =  0.958   Deviance explained = 95.8%
## -REML = -5820.5  Scale est. = 0.00064775  n = 2601
```

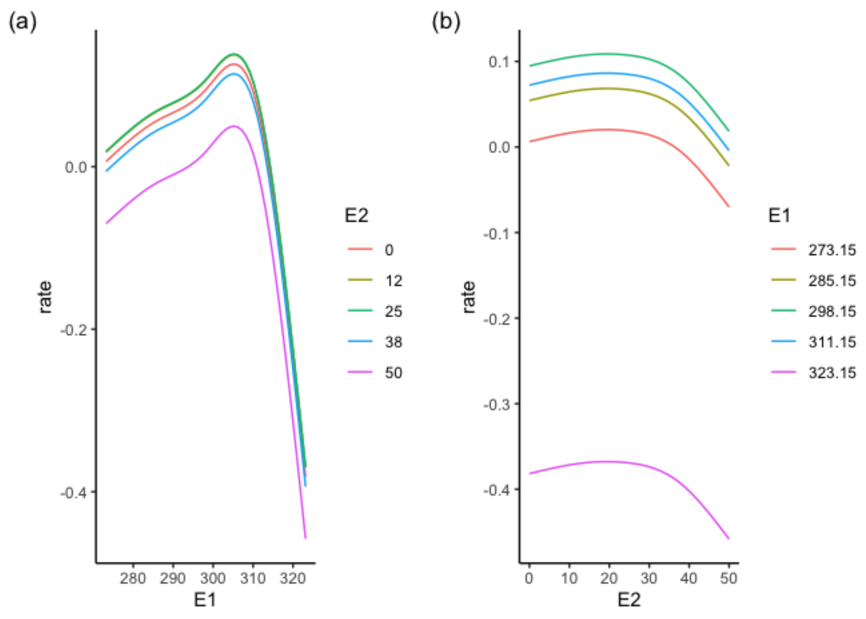

```
##
## Family: gaussian
## Link function: identity
##
## Formula:
## rate ~ s(E1) + s(E2)
##
## Parametric coefficients:
##              Estimate Std. Error t value Pr(>|t|)
## (Intercept) 0.0154285  0.0003952   39.04   <2e-16 ***
## ---
## Signif. codes:  0 '***' 0.001 '**' 0.01 '*' 0.05 '.' 0.1 ' ' 1
##
## Approximate significance of smooth terms:
##         edf Ref.df       F p-value
## s(E1) 8.970  9.000 10199.8  <2e-16 ***
## s(E2) 7.146  8.187   475.4  <2e-16 ***
## ---
## Signif. codes:  0 '***' 0.001 '**' 0.01 '*' 0.05 '.' 0.1 ' ' 1
##
## R-sq.(adj) =  0.974   Deviance explained = 97.4%
## -REML = -6407.3  Scale est. = 0.00040624  n = 2601
```

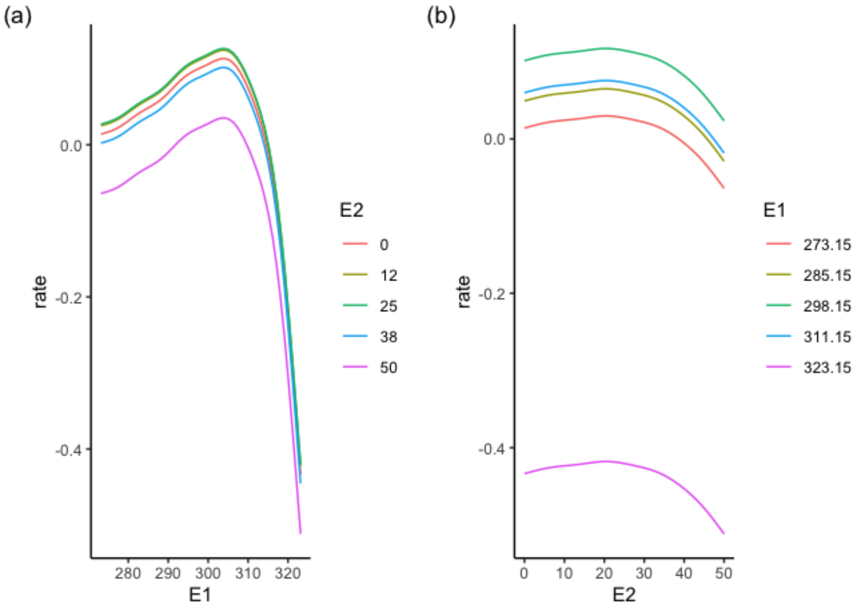


#### 3.3.2 With interaction

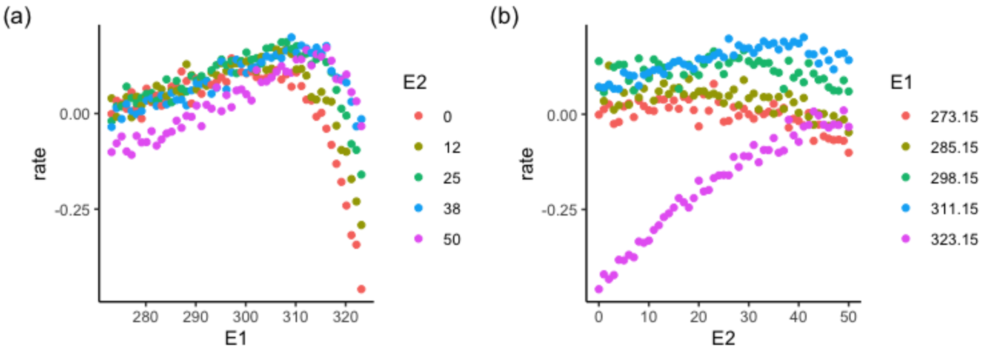


Figure 3.10: Performance curves for a species *with* interacting environmental effects and with some noise in the rate.

```
##
## Family: gaussian
## Link function: identity
##
## Formula:
## rate ~ ti(E1) + ti(E2) + te(E1, E2)
##
## Parametric coefficients:
##             Estimate Std. Error t value Pr(>|t|)
## (Intercept) 0.0656082  0.0004677   140.3   <2e-16 ***
## ---
## Signif. codes:  0 '***' 0.001 '**' 0.01 '*' 0.05 '.' 0.1 ' ' 1
##
## Approximate significance of smooth terms:
##             edf Ref.df      F p-value
## ti(E1)     3.991  3.999 1899.8  <2e-16 ***
## ti(E2)     3.991  4.000  302.6  <2e-16 ***
## te(E1,E2) 15.614 16.000  489.1  <2e-16 ***
## ---
## Signif. codes:  0 '***' 0.001 '**' 0.01 '*' 0.05 '.' 0.1 ' ' 1
##
## R-sq.(adj) =  0.924   Deviance explained = 92.5%
## -REML = -5943.3  Scale est. = 0.00056892  n = 2601
```

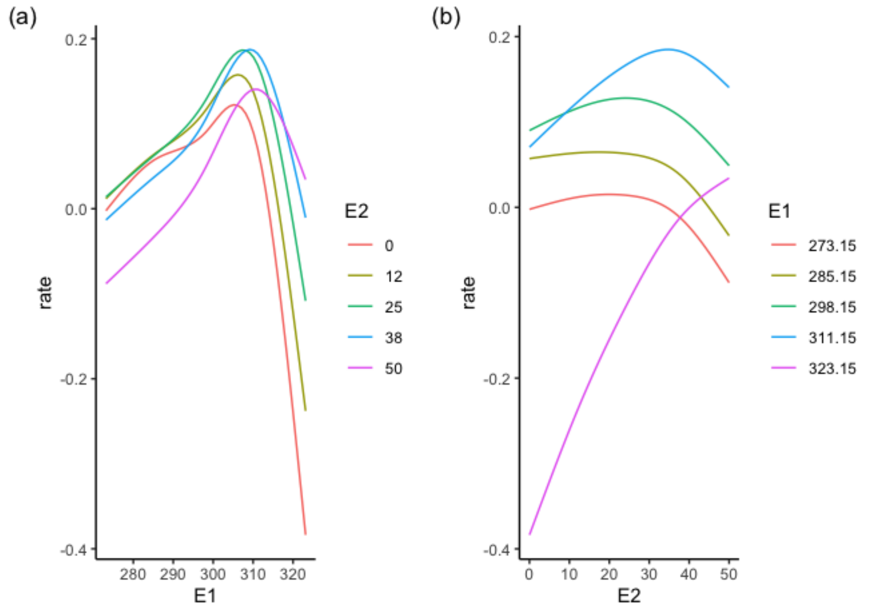


```
##
## Family: gaussian
## Link function: identity
##
## Formula:
## rate ~ s(E1) + s(E2)
##
## Parametric coefficients:
##             Estimate Std. Error t value Pr(>|t|)
## (Intercept) 0.0656082  0.0009042   72.56   <2e-16 ***
## ---
## Signif. codes:  0 '***' 0.001 '**' 0.01 '*' 0.05 '.' 0.1 ' ' 1
##
## Approximate significance of smooth terms:
##         edf Ref.df     F p-value
## s(E1) 8.561  8.945 672.0  <2e-16 ***
## s(E2) 4.421  5.434 101.2  <2e-16 ***
## ---
## Signif. codes:  0 '***' 0.001 '**' 0.01 '*' 0.05 '.' 0.1 ' ' 1
##
## R-sq.(adj) =  0.717   Deviance explained = 71.8%
## GCV = 0.0021379  Scale est. = 0.0021264  n = 2601
```

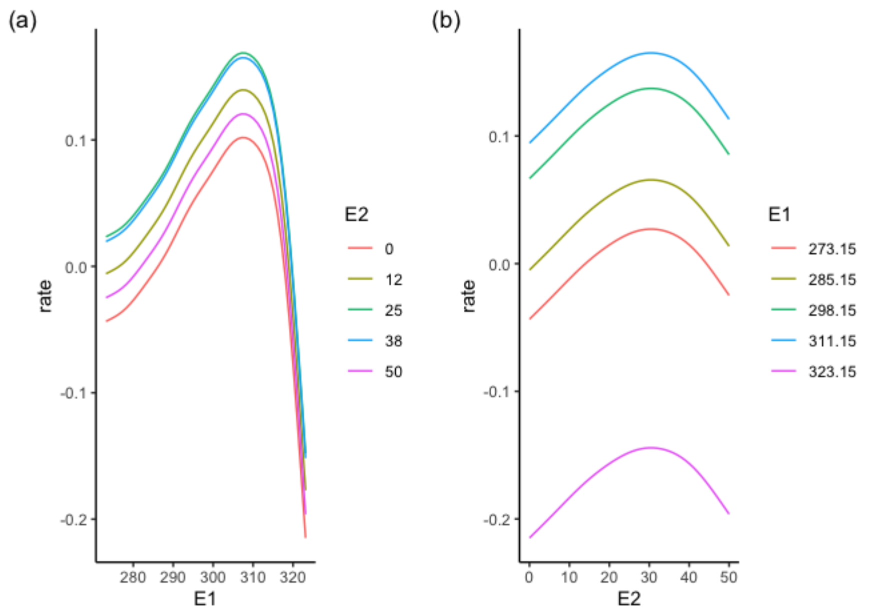


### 4 Partial derivatives

First step in calculating directional derivatives is estimating the two partial derivatives $f_{E1}(E1_0, E2_0)$ and $f_{E2}(E1_0, E2_0)$ (please review the section The principle if necessary).

#### 4.1 Getting the partial derivatives

Partial derivatives. Draw response surface for sp 1 and calculate partial derivatives at a specific location (E1 = 300, E2 = 20). To calculate the partial derivative with respect to E1, E2 must be held constant.

Visualising the partial effect of E1 at a fixed level of E2.

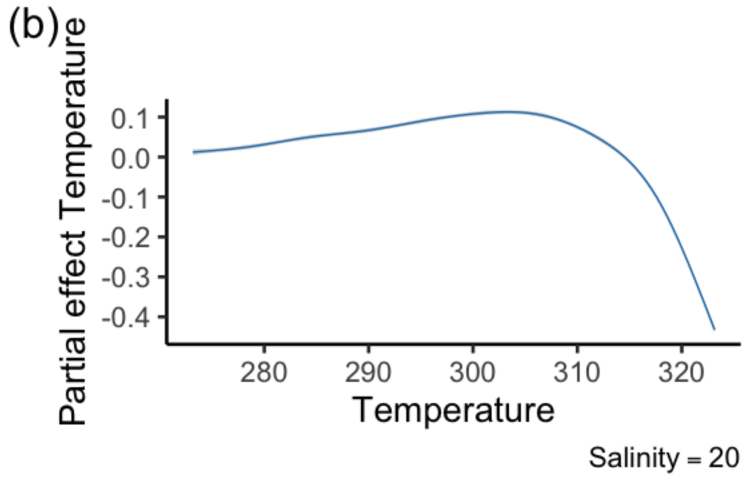


Figure 4.1: Partial effect of E1 on the growth rate of sp 1 when E2 is held constant at E2 = 20.

Partial derivative with respect to E1 when E2 is constant at 20.

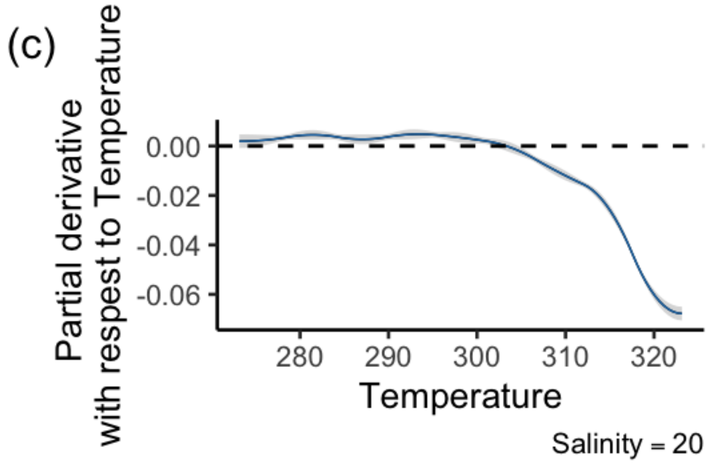


Figure 4.2: Partial derivative with respect to E1 when E2 is constant at 20.

Partial derivatives with respect to E2 (E1 held constant)

Partial effect of E2 on the growth rate of sp 1 when E1 is held constat at E2 = 300

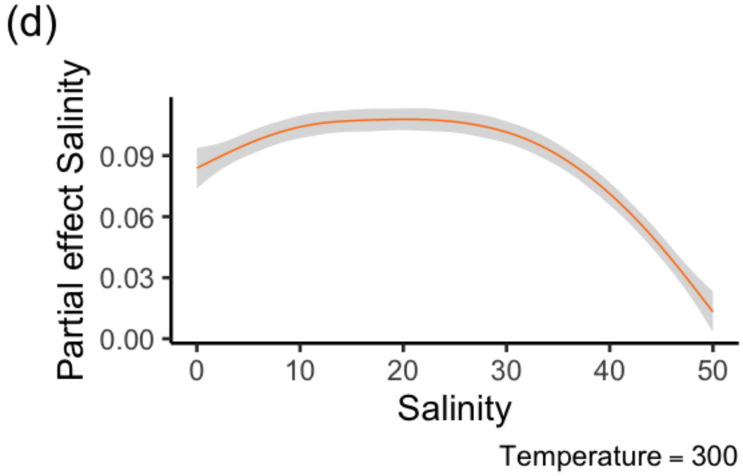


Figure 4.3: Partial effect of E2 on the growth rate of sp 1 when E1 is held constant at E1 = 300

Partial derivative with respect to E2 when E1 is constant at 300

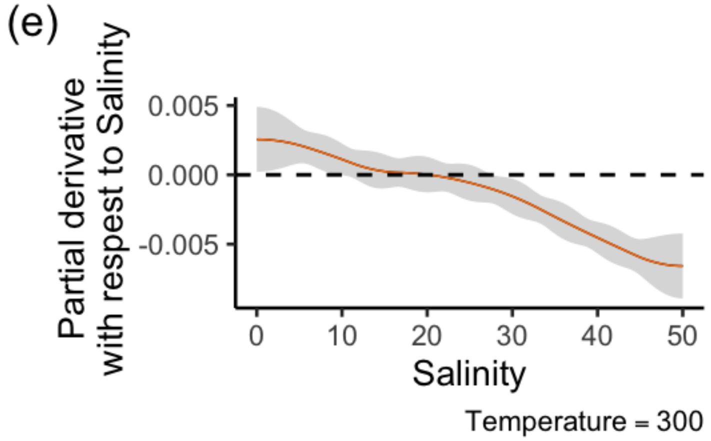


Figure 4.4: Partial derivative with respect to E1 when E2 is constant at 20.

Plot the two partial derivatives and relative effects

### 5 Directional derivatives

#### 5.1 No direction of environmental change specified

#### 5.1.1 One point

We start showing how directional derivatives can be calculated even when the direction of the environmental change is unknown. This may be the case when we want to calculate response diversity for future scenarios, and the future direction of environmental change is thus not known. Or we may have data for a species or a community at only one environmental location (E1 = x, E2 = y). It is therefore important to be able to measure directional derivatives when the direction of the environmental change is unknown, as this can provide useful information on response diversity nonetheless, for instance, by taking the mean of the slopes calculates in all directions.
Measuring response diversity when the direction of environmental change is unknown may represent a way to systematically measuring response diversity to all possible environmental changes. This is represents, in our view, an absolute measure of overall response diversity, since it captures the complete insurance capacity of a system under all possible environmental conditions. We thus put some emphasis on this approach here.

Here, we calculate, for a specific point (E1 = 300, E2 = 20), directional derivatives in all directions.

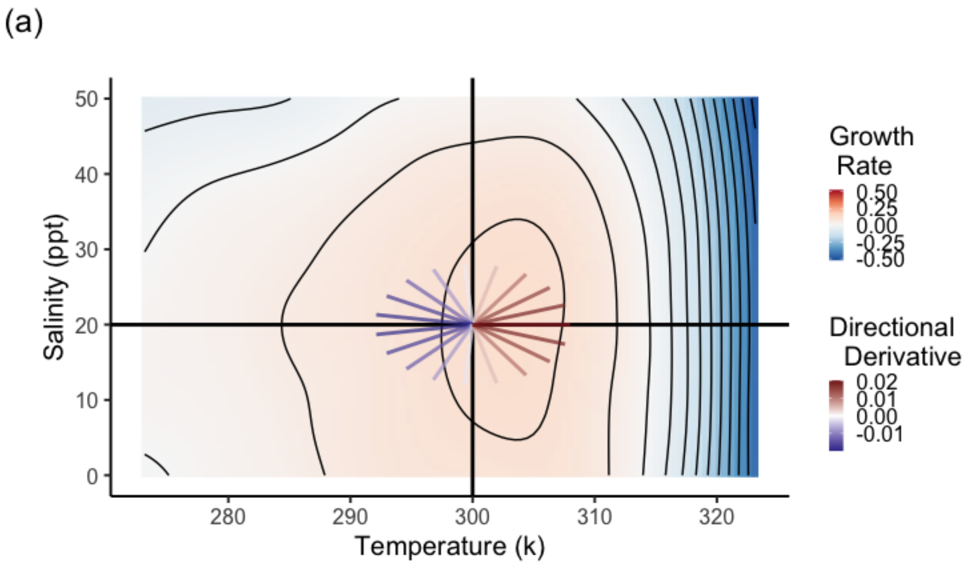


Figure 5.1: Directional derivatives calculated in all possible direction for a specific point on the response surface of sp1. Clearly, the slope of the directional derivative depends on the direction (red positive, blue negative). Note: the size of the radius was only chosen for representation purposes, and does not have any implication. The slope of the segments departing from the point have each their fixed slopes independently of the size of the radius.

#### 5.1.2 Several points

We can measure all possible directional derivatives also for several points on the surface. This might be the case when we know that a species or a community occurs at multiple locations on the surface (multiple environmental conditions), but we do not know the direction of change.

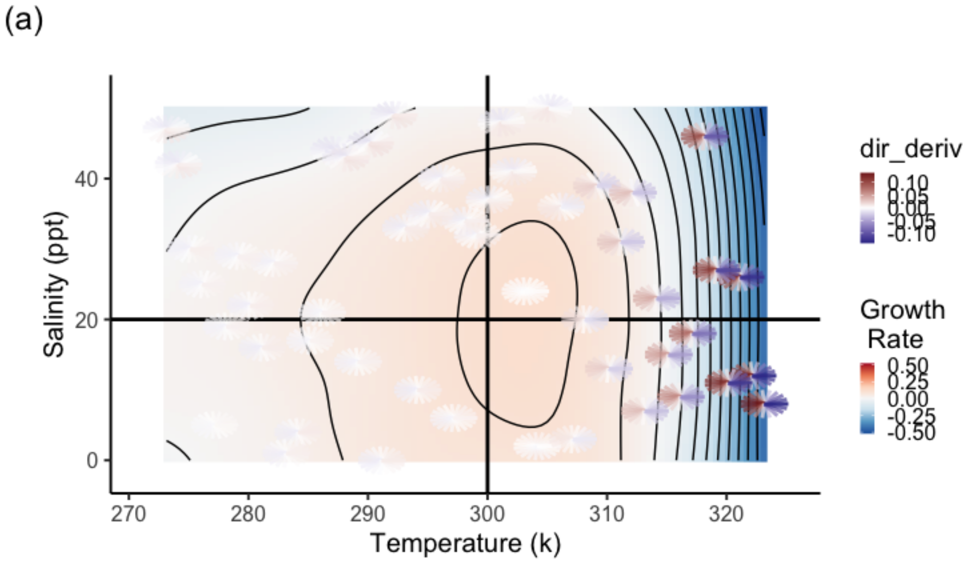


Figure 5.2: Directional derivatives calculated in all possible direction for several points on the response surface of sp1. Clearly, the slope of the directional derivative depends on the direction (red positive, blue negative).

##### 5.1.3 Grid of points

Finally, we might do the same for a grid of points on the surface. We may want to do that when we do not have information on where a species or a community is living within the surface, but we know the range of values of E1 and E2.

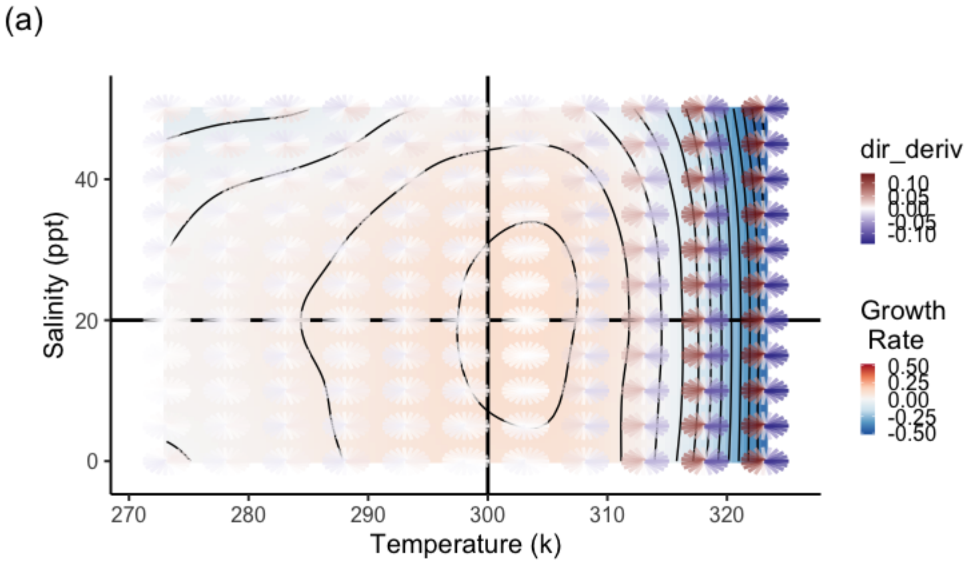


Figure 5.3: Directional derivatives calculated in all possible direction for a grid of points on the response surface of sp1. Clearly, the slope of the directional derivative depends on the direction (red positive, blue negative).

### 6 Response diversity calculation

Environmental variables may show different correlations between each other. The increase in one environmental variable may be directly correlated with the increase of another one (positive correlation), or vice versa, the increase in one driver may be correlated to a decrease in the other one (negative correlation). Yet, two environmental variables may change over time, or space, completely independently.
We may imagine that these different types of relationships between two environmental variables could determine specific trends in response diversity.

To explore this hypothesis, we calculate now response diversity for two communities (one with additive effect, and one including an interactive environmental effect) composed of 4 spp in 4 different cases: 1. Unknown direction of the environmental change 2. Direction of env change is given by the time series,and E1 and E2 change over time independently 3. Direction of env change is given by the time series,and E1 and E2 change over time with positive correlation 4. Direction of env change is given by the time series,and E1 and E2 change over time with negative correlation

We want to see if any consistent trend appears in the two communities when E1 and E2 have different correlations.

Steps:

i. Simulate spp performance curves with the modified Eppley function with and without interactive effect.

ii. Fit response surface for each sp (done with GAMs)

iii. Data wrangling and partials derivatives calculations

#### 6.1 Community 1 - without interactive effect

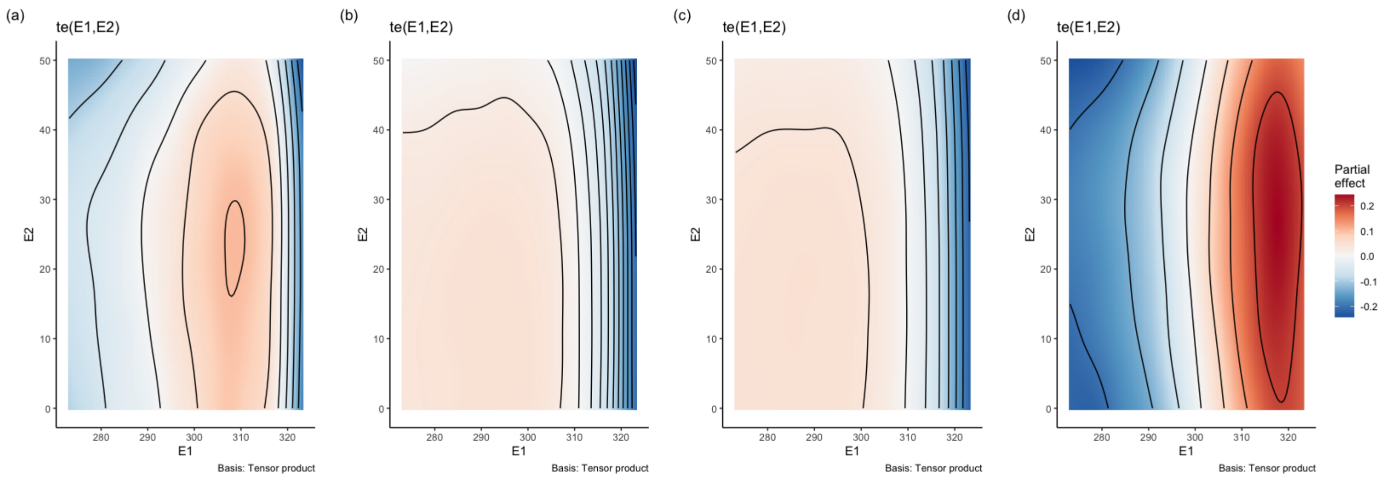


Figure 6.1: Response surface of the three species composing community 1. (a) Sp4. (b) Sp6. (c) Sp 11

#### 6.2 Community 2 - interactive effect

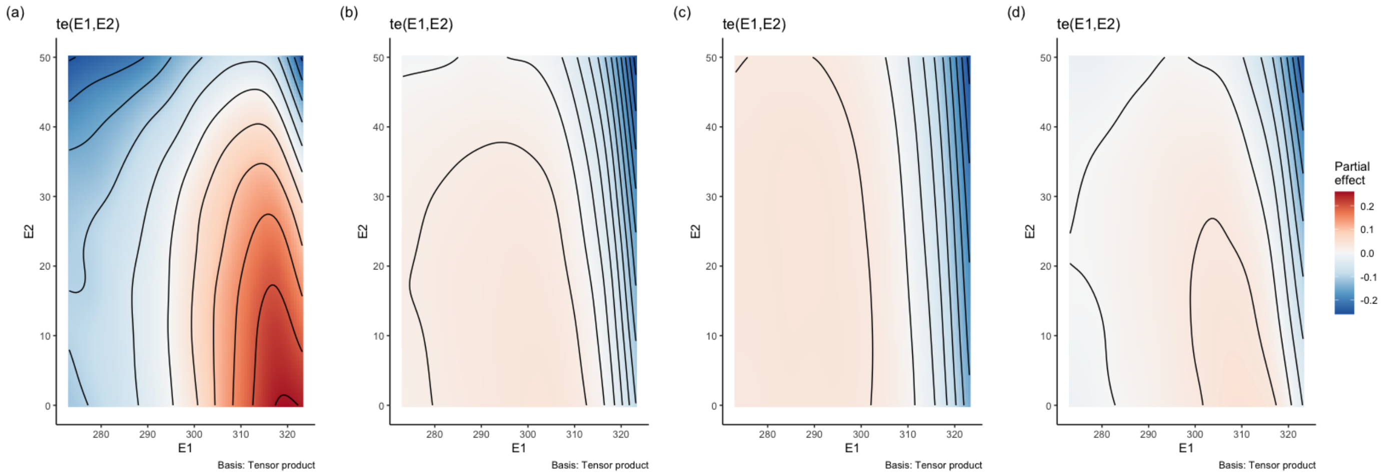


Figure 6.2: Response surface of the three species composing community 1. (a) Sp2. (b) Sp5. (c) Sp 13

#### 6.3 Unknown direction of the environmental change

Table showing the calculated response diversity for one of the two communities when the direction of the environmental change is unknown. In this case, we calculated response diversity for a community in all possible directions across the surface, which represents in our opinion the most sensible way to measure the absolute response diversity of a specific community.

| divergence | dissimilarity | community |
|---:|---:|---:|
| 0.2693448 | 1.011636 | 1 |
| 0.2473904 | 1.010359 | 2 |

##### 6.3.1 E1 and E2 change independently over time

This example mimics a situation where the two environmental variables change over time completely independently. This is a common situation in field studies, where multiple drivers of environmental change are not correlated one another.

In this case the direction of the environmental change is given by the change of E1 and E2 over time.

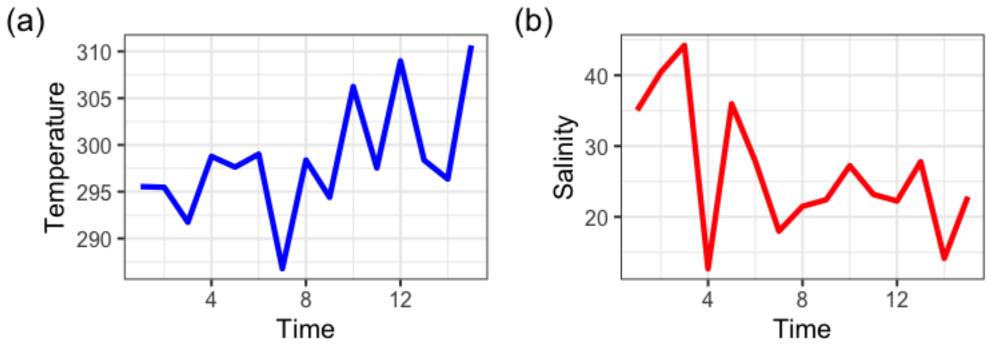


Figure 6.3: Time series of E1 and E2 changing independently over time.

##### 6.3.2 Response surfaces with change in environmental conditions

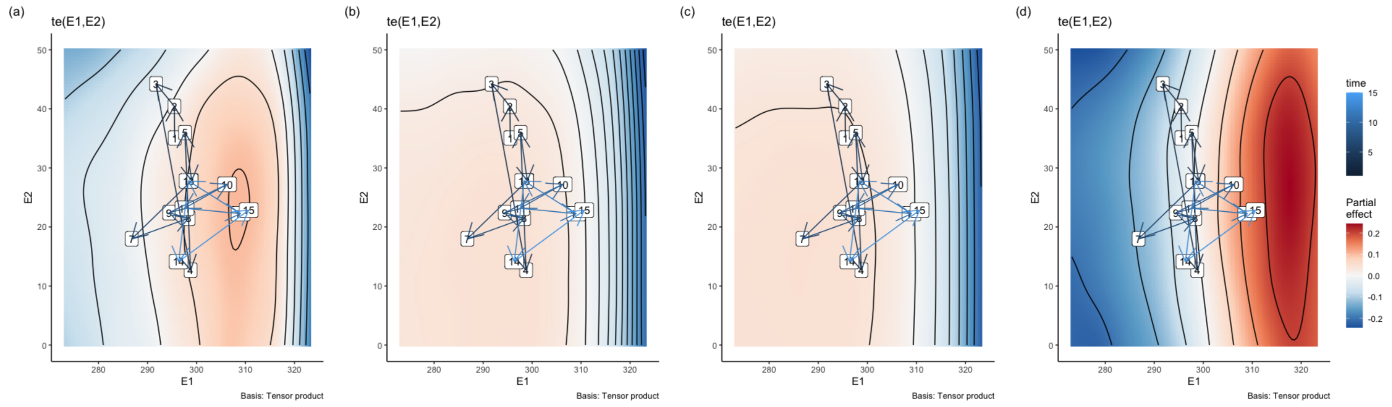

Figure 6.4: Response surface of the three species composing community 1. (a) Sp4. (b) Sp6. (c) Sp 11. The numbers on the response surfaces show the environmental location in the time steps of the time series and the arrows connect the time steps.

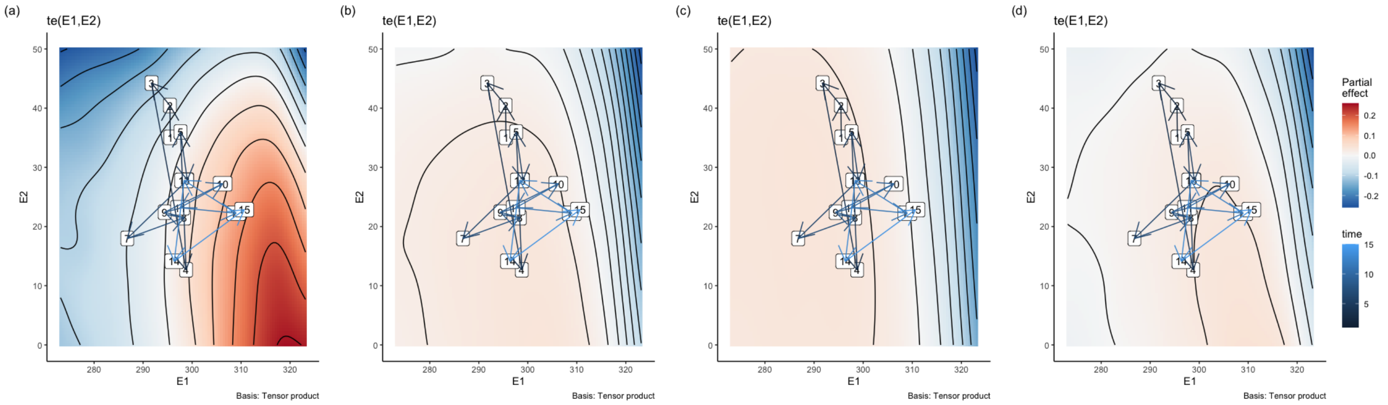


Figure 6.5: Response surface of the three species composing community 2. (a) Sp2. (b) Sp5. (c) Sp 13

Table showing the calculated response diversity for one of the two communities when the two environmental variables change independently over time (only first 6 rows shown).

| time | E1_ref | E2_ref | s1 | s2 | s3 | s4 | rdiv | sign | Med |
|---|---|---|---|---|---|---|---|---|---|
| 1 | 295.5499 | 35.06010 | -0.0025569 | -0.0043734 | -0.0031682 | -0.0019315 | 1.000992 | 0.0000000 | 1.004178 |
| 2 | 295.4694 | 40.43676 | -0.0076105 | -0.0038792 | -0.0010685 | -0.0079641 | 1.003054 | 0.0000000 | 1.004178 |
| 3 | 291.7350 | 44.23112 | 0.0059942 | 0.0082445 | 0.0069416 | 0.0049920 | 1.001339 | 0.0000000 | 1.004178 |
| 4 | 298.7794 | 12.68533 | 0.0010253 | -0.0008450 | 0.0011959 | 0.0006765 | 1.000809 | 0.8280425 | 1.004178 |
| 5 | 297.6343 | 35.97065 | 0.0033619 | 0.0040078 | 0.0024411 | 0.0033568 | 1.000588 | 0.0000000 | 1.004178 |
| 6 | 299.0214 | 27.84749 | -0.0045781 | 0.0049459 | 0.0077936 | -0.0073761 | 1.006887 | 0.9724763 | 1.004178 |

Plot response diversity over time

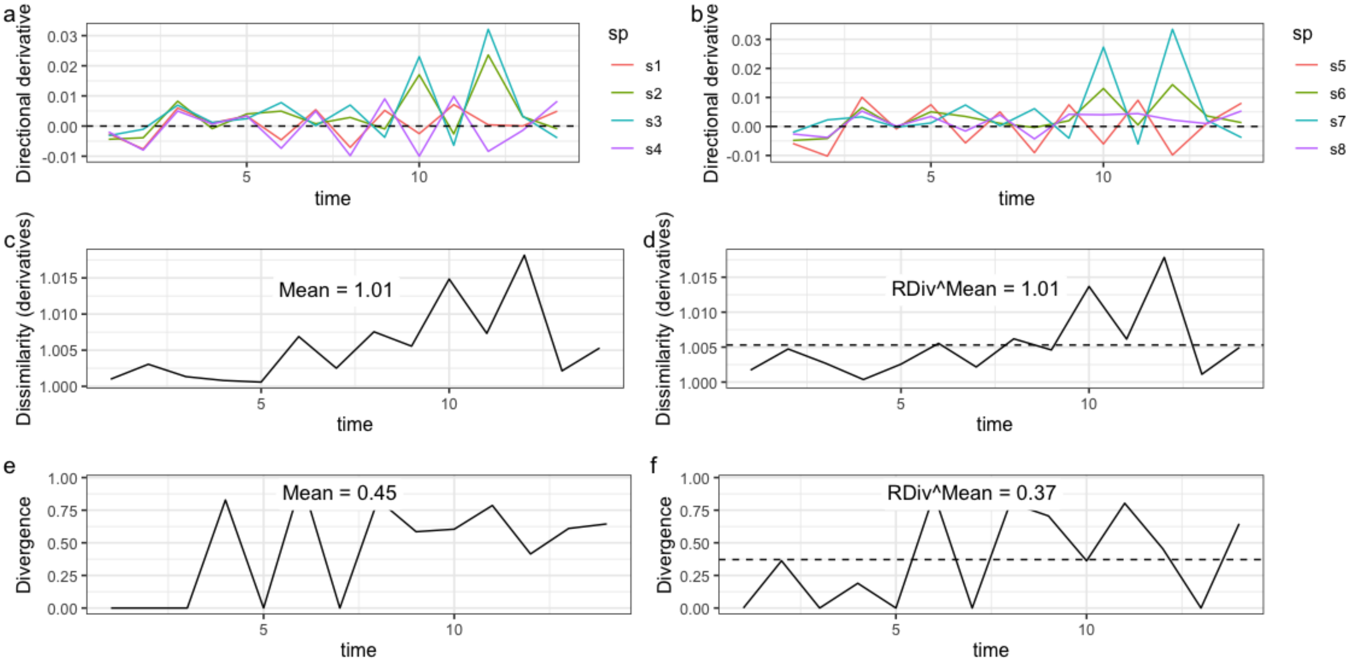


Figure 6.6: Directional derivatives and response diversity with known direction of env change. E1 and E2 change independently over time. a and b: Species directional derivatives over time. c and d: Response diversity measured as similarity-based diversity metric. e and : Response diversity measured as divergence (sign sensitive).

#### 6.4 E1 and E2 change with negative correlation

This example mimics a situation where the two environmental variables change over time with negative correlation. This is common in field studies, where one environmental variable (e.g. CO2 concentration in oceans) increases, while another (e.g. pH) decreases e.g. Shirayama & Thornton (2005) (https://agupubs.onlinelibrary.wiley.com/doi/full/10.1029/2004JC002618).

Creating a time series with E1 and E2 changing over time with negative correlation.

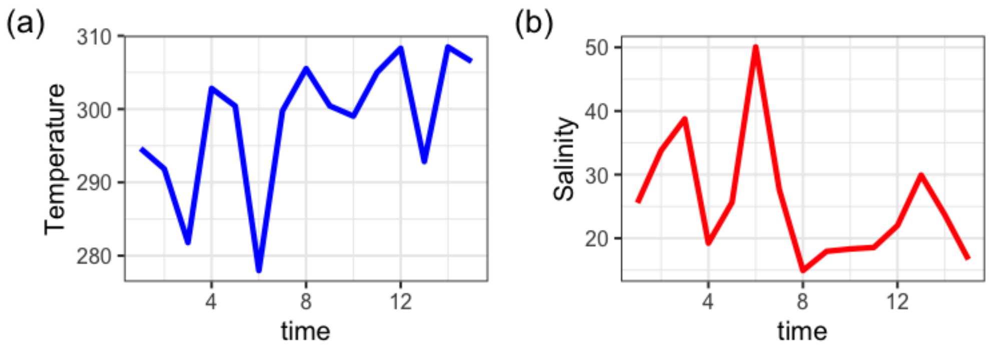


Figure 6.7: Time series of E1 and E2 changing with negative correlation over time.

#### 6.4.1 Response surfaces with change in environmental conditions

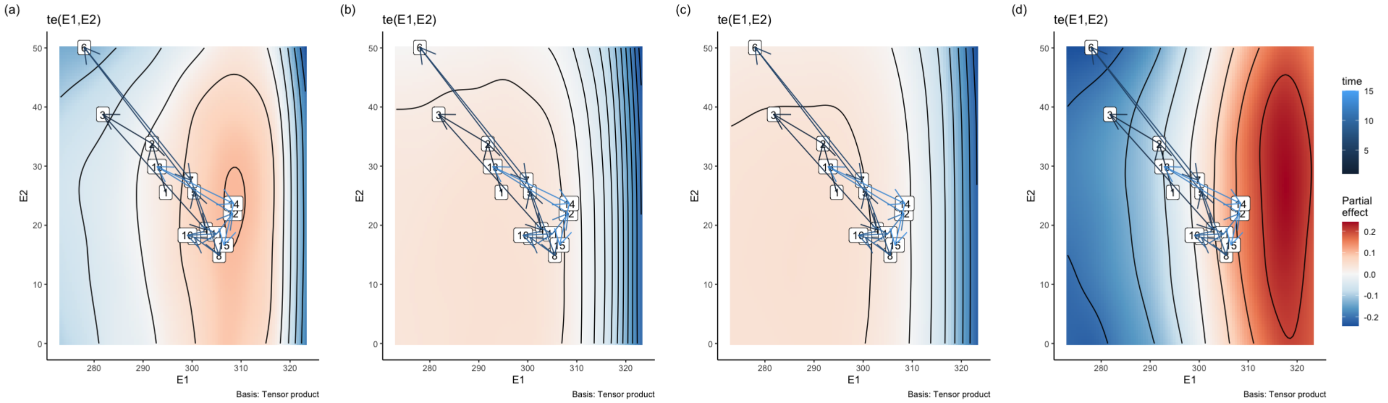


Figure 6.8: Response surface of the three species composing community 1. (a) Sp4. (b) Sp6. (c) Sp 11. The numbers on the response surfaces show the environmental location in the time steps of the time series and the arrows connect the time steps.

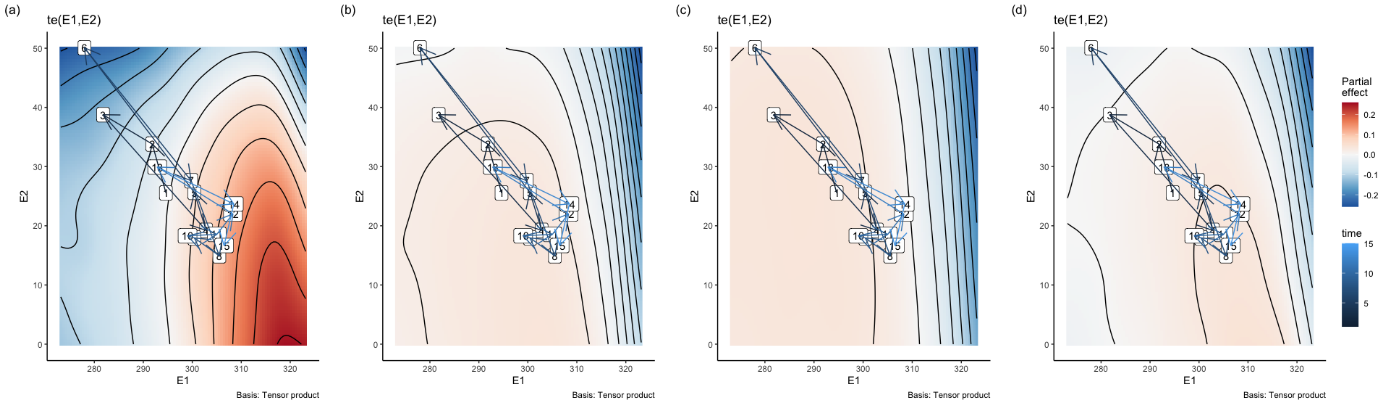


Figure 6.9: Response surface of the three species composing community 2. (a) Sp2. (b) Sp5. (c) Sp 13

Table showing the calculated response diversity for one of the two communities when the two environmental variables change with negative correlation over time (only first 6 rows shown).

| time | E1_ref | E2_ref | s1 | s2 | s3 | s4 | rdiv | sign | Med |
|---|---|---|---|---|---|---|---|---|---|
| 1 | 294.6225 | 25.55104 | -0.0017809 | -0.0012320 | -0.0007742 | -0.0023943 | 1.000676 | 0.0000000 | 1.004724 |
| 2 | 291.8397 | 33.79961 | -0.0055741 | -0.0023475 | -0.0009544 | -0.0080189 | 1.003054 | 0.0000000 | 1.004724 |
| 3 | 281.7902 | 38.76077 | 0.0045087 | 0.0057514 | 0.0042607 | 0.0043408 | 1.000580 | 0.0000000 | 1.004724 |
| 4 | 302.8219 | 19.22634 | -0.0013861 | 0.0018754 | 0.0048419 | -0.0030857 | 1.003383 | 0.7784758 | 1.004724 |
| 5 | 300.4154 | 25.64585 | -0.0054512 | 0.0020723 | 0.0052478 | -0.0071079 | 1.005579 | 0.8494616 | 1.004724 |
| 6 | 277.9783 | 50.06561 | 0.0066401 | 0.0091631 | 0.0071840 | 0.0046479 | 1.001762 | 0.0000000 | 1.004724 |

Plot response diversity over time for the two communities

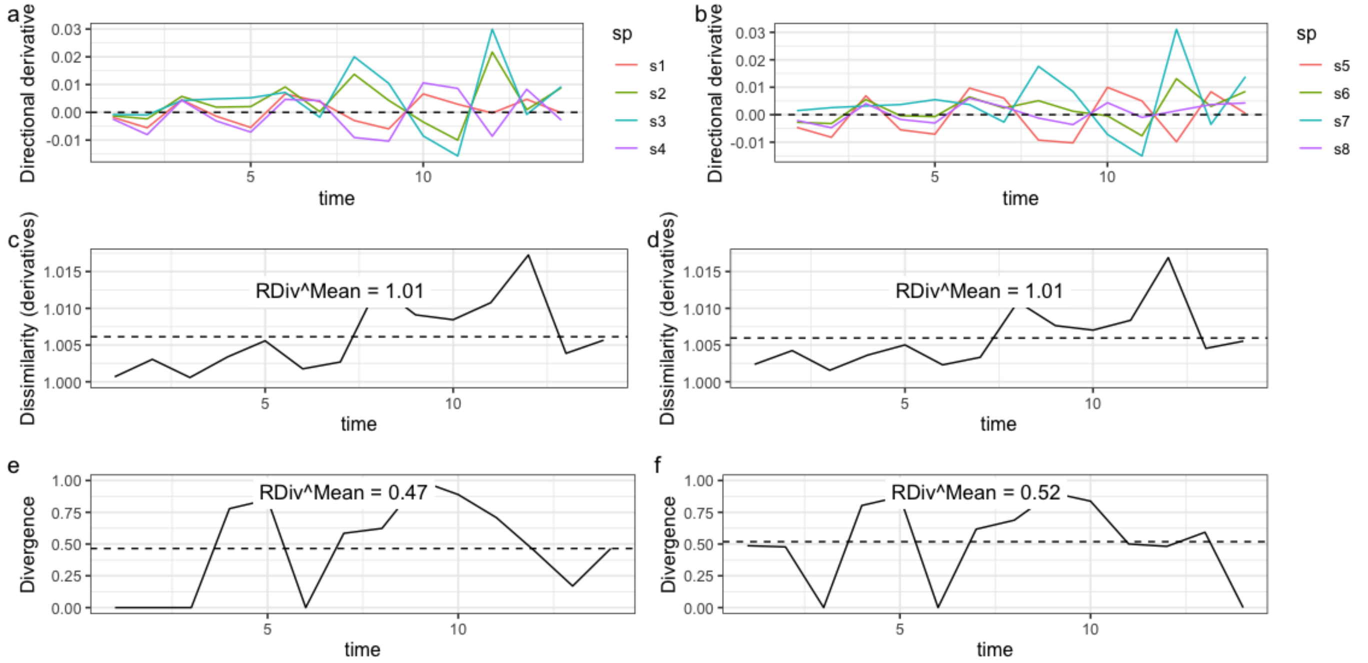


Figure 6.10: Directional derivatives and response diversity with known direction of env change. E1 and E2 change with negative correlation over time. a and b: Species directional derivatives over time. c and d: Response diversity measured as similarity-based diversity metric. e and : Response diversity measured as divergence (sign sensitive).

### 6.5 E1 and E2 change with positive correlation

Finally, two environmental variables can show positive correlation over time. A typical example is given by the positive correlation between air temperature and UV radiation e.g. Häder at al. 2015 (https://pubs.rsc.org/en/content/articlehtml/2015/pp/c4pp90035a).

Let us create a time series with E1 and E2 changing over time with positive correlation

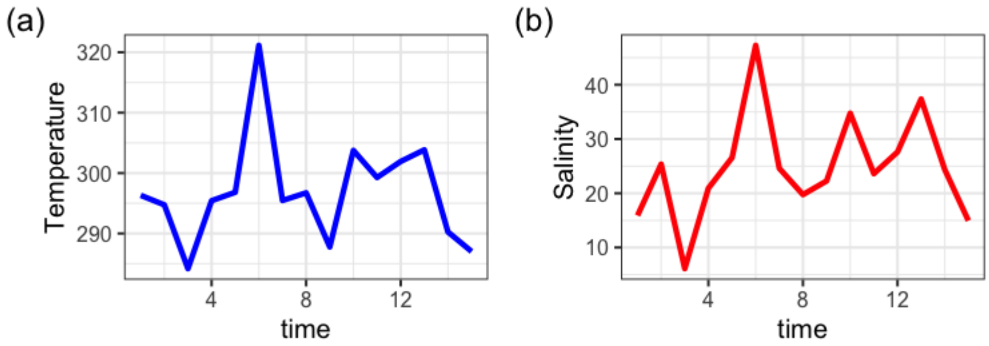


Figure 6.11: Time series of E1 and E2 changing with positive correlation over time.

#### 6.5.1 Response surfaces with change in environmental conditions

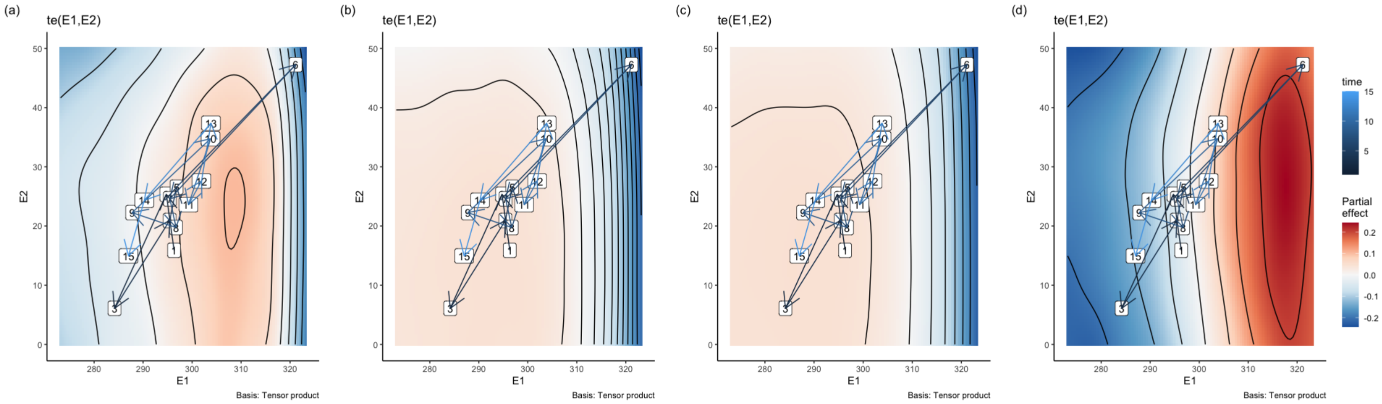


Figure 6.12: Response surface of the three species composing community 1. (a) Sp4. (b) Sp6. (c) Sp 11. The numbers on the response surfaces show the environmental location in the time steps of the time series and the arrows connect the time steps.

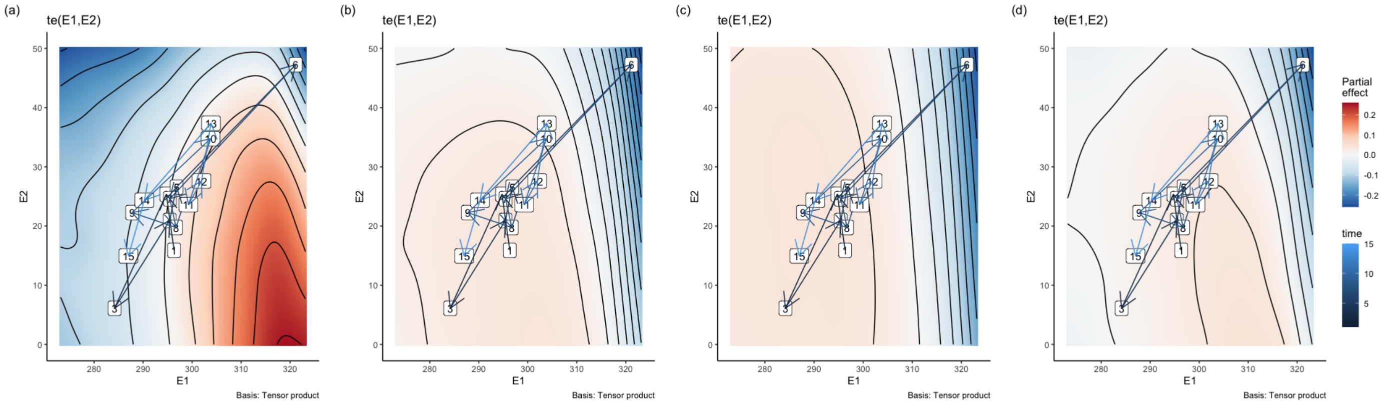


Figure 6.13: Response surface of the three species composing community 2. (a) Sp2. (b) Sp5. (c) Sp 13

Table showing the calculated response diversity for one of the two communities when the two environmental variables change with positive correlation over time (only first 6 rows shown).

| time | E1_ref | E2_ref | s1 | s2 | s3 | s4 | rdiv | sign | Med |
|---:|---:|---:|---:|---:|---:|---:|---:|---:|---:|
| 1 | 296.3194 | 15.893276 | -0.0008065 | -0.0004111 | 0.0007374 | -0.0006908 | 1.000614 | 0.9552337 | 1.003235 |
| 2 | 294.7370 | 25.358621 | -0.0023788 | 0.0019470 | 0.0036368 | -0.0051287 | 1.003831 | 0.8297969 | 1.003235 |
| 3 | 284.1721 | 6.102644 | 0.0017780 | 0.0020120 | 0.0020059 | 0.0043266 | 1.000957 | 0.0000000 | 1.003235 |
| 4 | 295.4237 | 20.931492 | 0.0013011 | -0.0008613 | -0.0021201 | 0.0033618 | 1.002327 | 0.7735041 | 1.003235 |
| 5 | 296.8193 | 26.556820 | 0.0043070 | -0.0033500 | -0.0059004 | 0.0074102 | 1.005955 | 0.8865661 | 1.003235 |
| 6 | 321.1482 | 47.260206 | 0.0332335 | 0.0889998 | 0.1072674 | 0.0070665 | 1.044901 | 0.0000000 | 1.003235 |

Plot response diversity over time

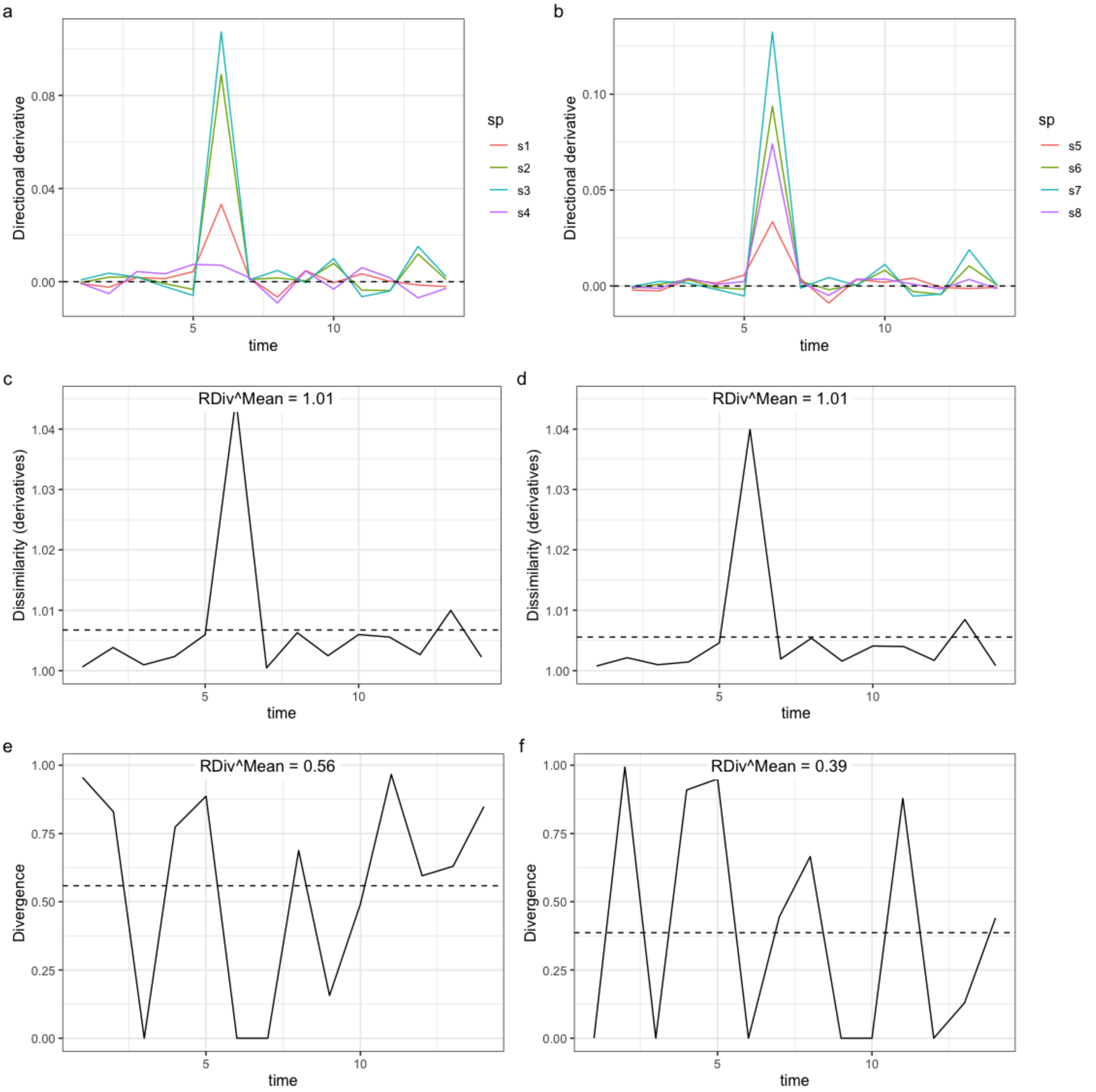


Figure 6.14: Directional derivatives and response diversity with known direction of env change for community 1 and 2. E1 and E2 change with negative correlation over time.a and b: Species directional derivatives over time. c and d: Response diversity measured as similarity-based diversity metric. e and : Response diversity measured as divergence (sign sensitive).

Now, we visualize the relationship between different correlations between the two environmental variables and response diversity.

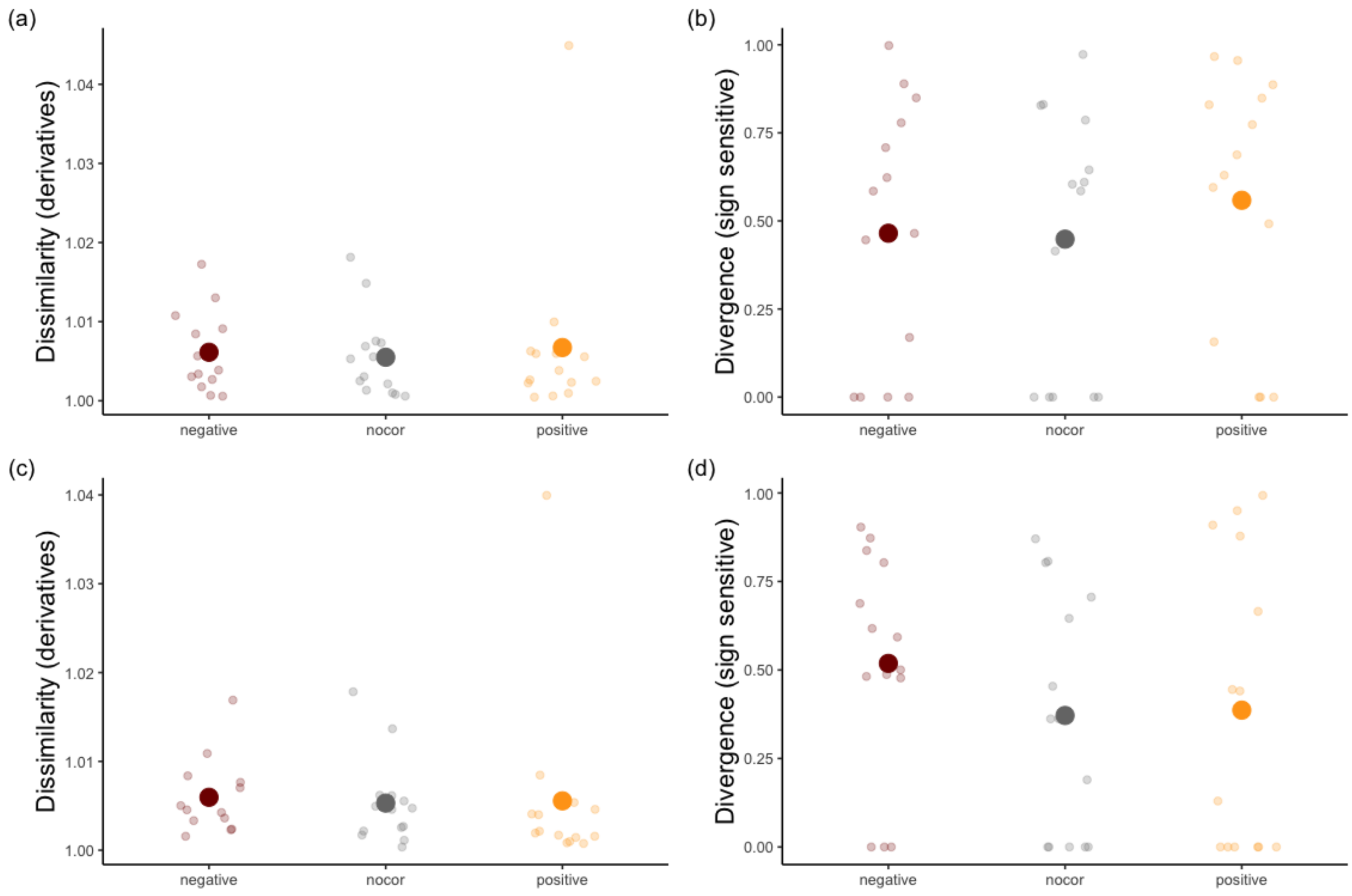


Figure 6.15: Correlation types and response diversity. a and c: correlation types and response diversity measured as dissimilarity in the first derivatives (sign insensitive) for community 1 and 2 respectively. c and d. correlation types and response diversity measured as divergence in the first derivatives (sign sensitive) for community 1 and 2 respectively

We can rule out the hypothesis that different types of relationships between two environmental variables could determine specific trends in response diversity.

### 7 Empirical example

We use data coming from an experiment where individual ciliates species have been exposed to a gradient of nutrient, light, and their combinations in a factorial design. We first show how to calculate the partial derivatives, then we calculate the directional derivatives based on a simulated time series (in the original experiment, the level of the treatments have been kept constant throughout the expt duration). Finally, we assemble random composed communities and calculate response diversity for each of them.

(The original data and metadata are available at https://doi.org/10.5281/zenodo.8383688 (https://doi.org/10.5281/zenodo.8383688))

##### 7.0.1 Load data set and look at species responses

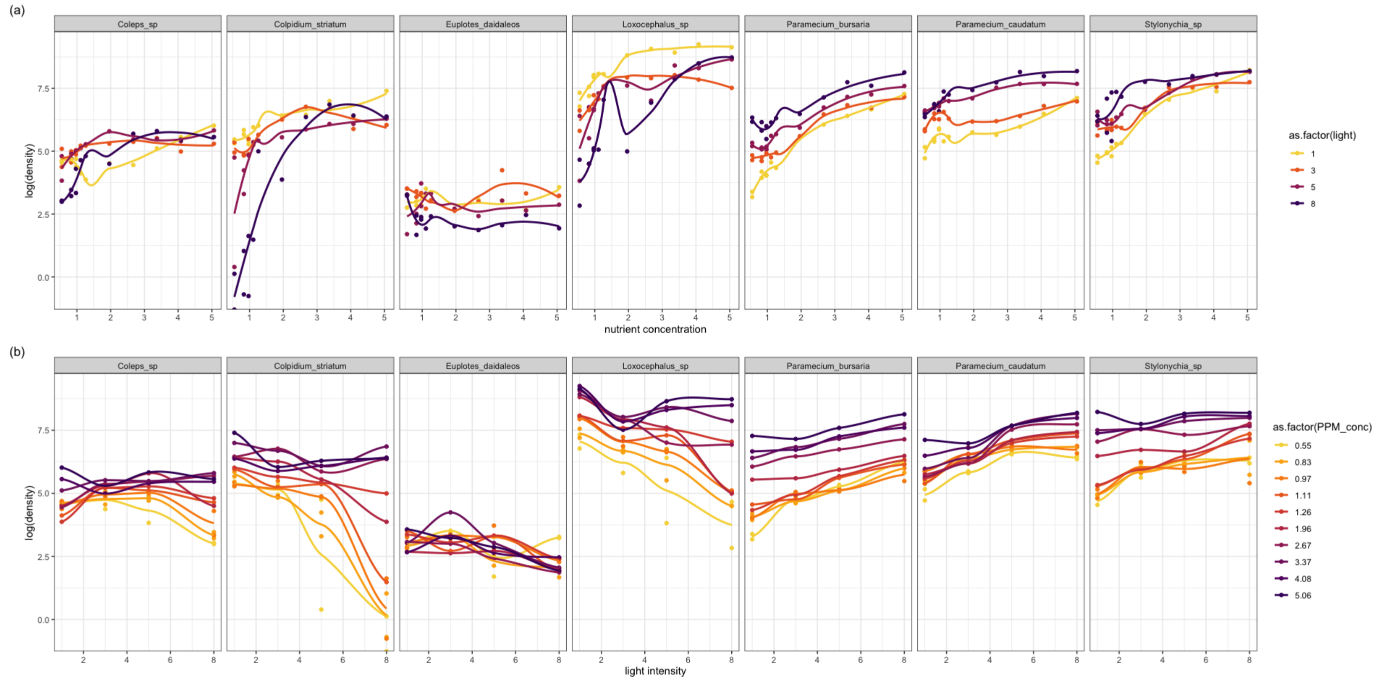


Figure 7.1: Species responses to the environmental drivers. a. Species responses to nutrient concentrations. b. Species responses to light intensity

#### 7.0.2 Fittig GAMs on empirical data

| species | E1 | E2 | predicted |
|---|---|---|---|
| Loxocephalus_sp | 0.55 | 1.00 | 1040.202 |
| Loxocephalus_sp | 0.55 | 1.02 | 1033.719 |
| Loxocephalus_sp | 0.55 | 1.04 | 1027.237 |
| Loxocephalus_sp | 0.55 | 1.06 | 1020.756 |
| Loxocephalus_sp | 0.55 | 1.08 | 1014.277 |
| Loxocephalus_sp | 0.55 | 1.10 | 1007.801 |

#### 7.0.3 Plotting surface for a sp

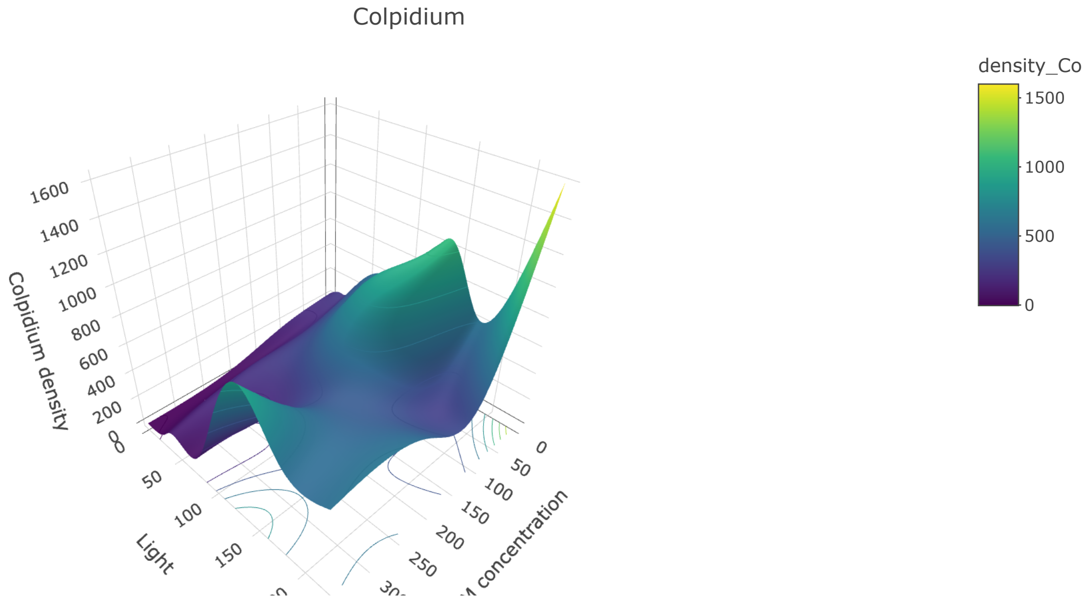


Figure 7.2: Response surface fitted with GAM. High non-linearity.

### 8 Partial derivatives for a single species

#### 8.1 E1 - Nutrients

First, we calculate the partial derivative with respect to nutrient concentration keeping light intensity constant at 5.

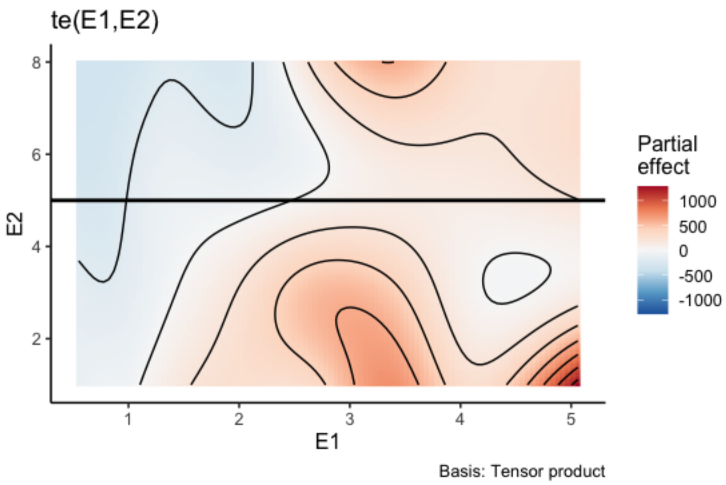


Figure 8.1: Response surface of Colpidium. The two solid lines show at which level of nutrients and light each partial derivative is going to be calculated.

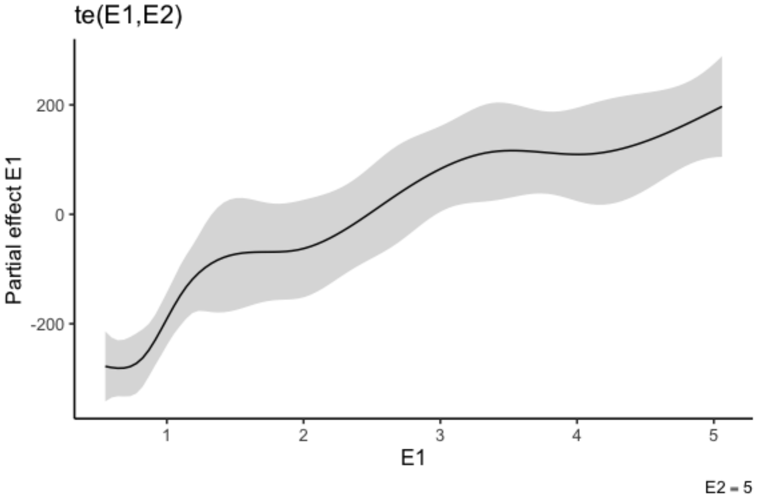


Figure 8.2: Partial effect of nutrient concentration on the growth rate of Colpidium when light intensity is held constant at 5.

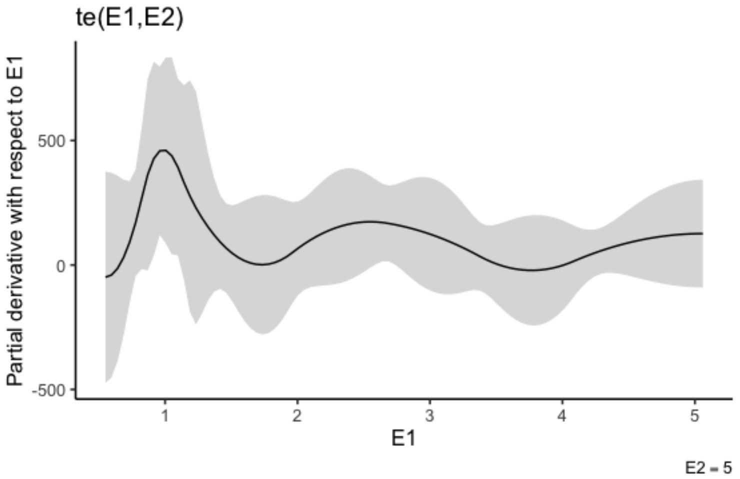


Figure 8.3: Partial derivative with respect to nutrient when light intensity is constant at 5.

#### 8.2 E2 - Light intensity

Second, we calculate the partial derivative with respect to light intensity keeping nutrient concentration constant at 2.67.

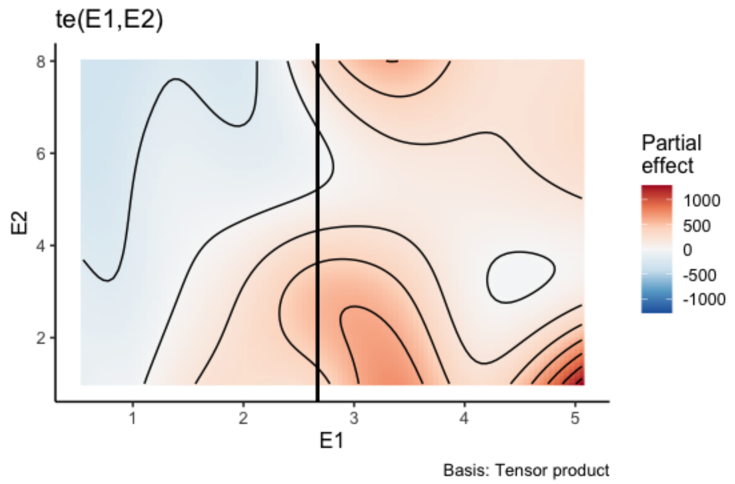


Figure 8.4: Response surface of Colpidium. The two solid lines show at which level of nutrients and light each partial derivative is going to be calculated. Not sure we get the gray areas…

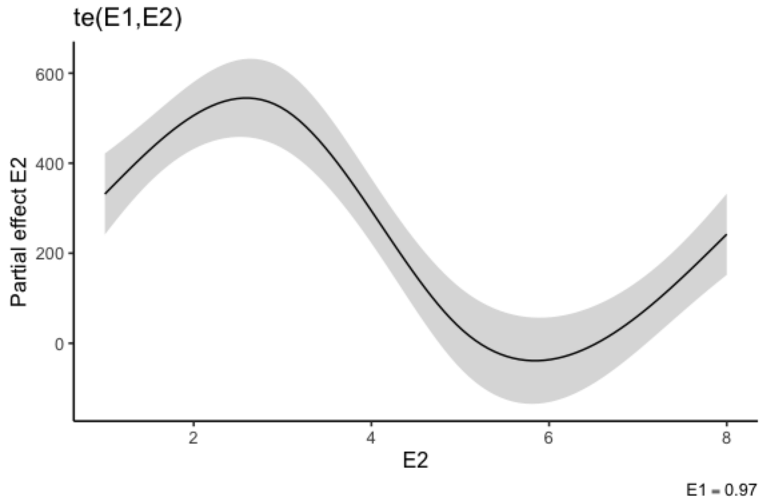


Figure 8.5: Partial effect of light intensity on the growth rate of Colpidium when nutrient concentration is held constant at 2.67.

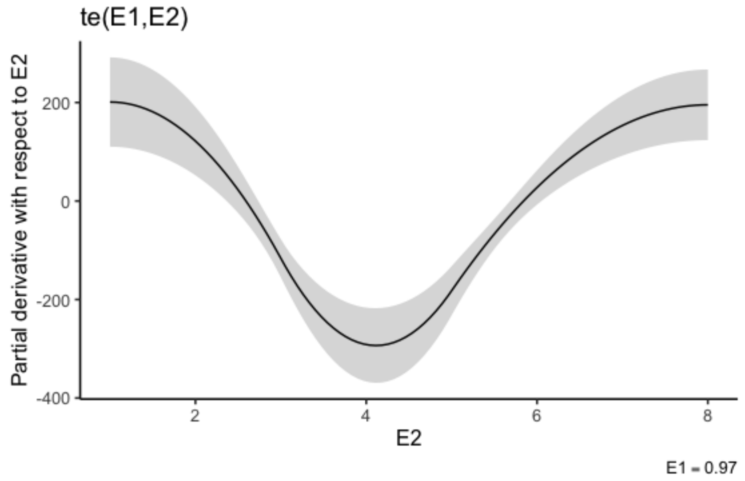


Figure 8.6: Partial derivative with respect to nutrient when light intensity is constant at 5.

##### 8.2.1 Plot surface and partial derivatives

Plot the two partial derivatives and relative effects

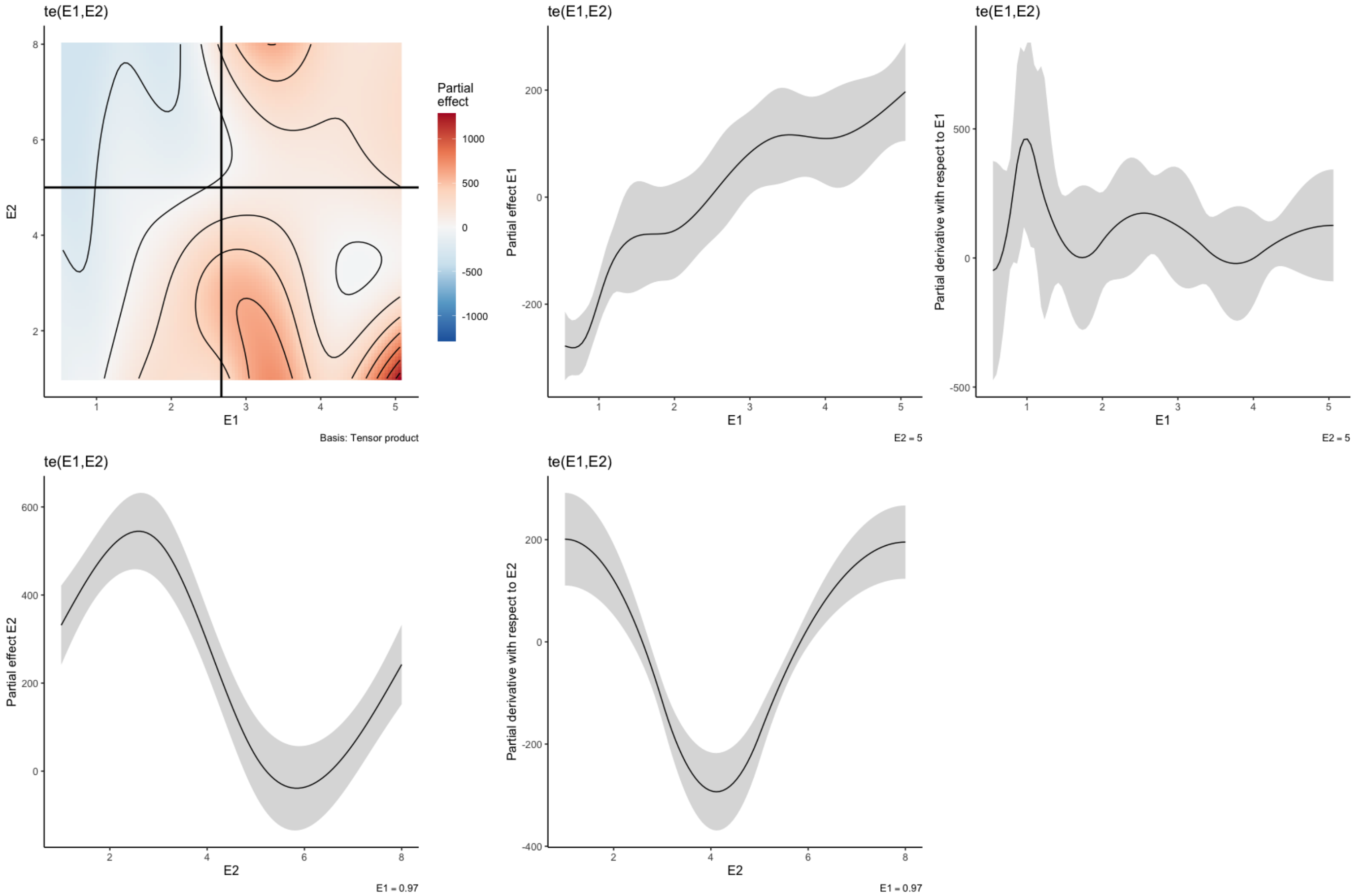


Figure 8.7: Summary plot of Colpidium (a) response surface of Colpidium (b) Partial effect of nutrient concentration on the density of Colpidium when light intensity is held constant at 5. (c) Partial derivative with respect to nutrient concentration when light intensity is held constant at 5. (d) Partial effect of light intensity on the growth rate of Colpidium when nutrient concentration is held constant at 2.67. (e) Partial derivative with respect to light intensity when nutrient concentration is held constant at 2.67.

### 9 Directional deriviatives

To calculate the directional derivatives for all spp used in the experiment, we first create a time series with nutrient concentration and light intensity changing randomly over time, we fit GAMs individually for each species, and then we calculate partial derivatives.

Time series of nutrient concentration and light intensity changing over time.

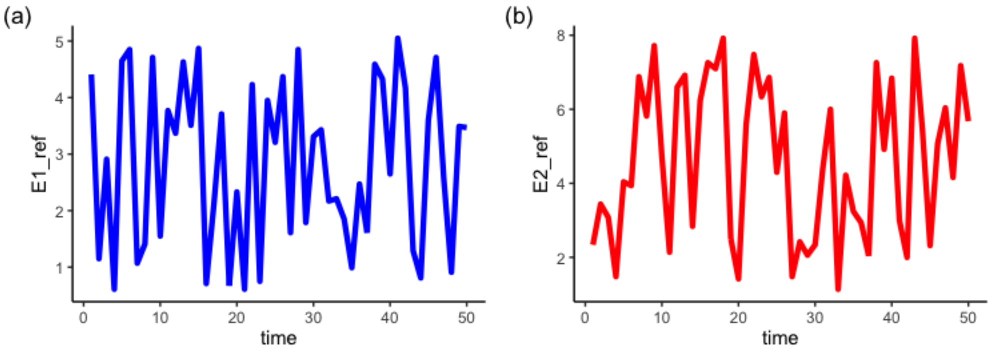


Figure 9.1: Time series of (a) nutrient concentration and (b) light intensity changing over time.

Table with calculated partial derivatives for each sp at different times (only first 6 rows shown).

| sp | time | E1_ref | E2_ref | pd_E1 | pd_E2 |
|---|---|---|---|---|---|
| Loxocephalus_sp | 1 | 4.41 | 2.34 | -697.5280 | -3111.3121 |
| Loxocephalus_sp | 2 | 1.15 | 3.44 | 1404.3671 | -280.5791 |
| Loxocephalus_sp | 3 | 2.91 | 3.08 | 149.0230 | -1170.1751 |
| Loxocephalus_sp | 4 | 0.61 | 1.48 | 3098.1877 | -390.3620 |
| Loxocephalus_sp | 5 | 4.65 | 4.04 | 231.7739 | 1889.1895 |

| sp | time | E1_ref | E2_ref | pd_E1 | pd_E2 |
|---|---|---|---|---|---|
| Loxocephalus_sp | 6 | 4.85 | 3.94 | 121.2058 | 1987.8819 |

#### 9.1 Calculating response diversity for a specific community composition

First, we need to calculate the directional derivatives in the direction of the env change.

| sp | time | E1_ref | E2_ref | pd_E1 | pd_E2 | nxt_value_E1 | nxt_value_E2 | del_E1 | del_E2 | unit_vec_mag | uv_E1 | uv_E2 | dir_deriv |
|---|---|---|---|---|---|---|---|---|---|---|---|---|---|
| Loxocephalus_sp | 1 | 4.41 | 2.34 | -697.5280 | -3111.3121 | 1.15 | 3.44 | -3.26 | 1.10 | 3.4405813 | -0.9475143 | 0.3197134 | -333.8104 |
| Loxocephalus_sp | 2 | 1.15 | 3.44 | 1404.3671 | -280.5791 | 2.91 | 3.08 | 1.76 | -0.36 | 1.7964409 | 0.9797149 | -0.2003962 | 1432.1064 |
| Loxocephalus_sp | 3 | 2.91 | 3.08 | 149.0230 | -1170.1751 | 0.61 | 1.48 | -2.30 | -1.60 | 2.8017851 | -0.8209052 | -0.5710645 | 545.9117 |
| Loxocephalus_sp | 4 | 0.61 | 1.48 | 3098.1877 | -390.3620 | 4.65 | 4.04 | 4.04 | 2.56 | 4.7828025 | 0.8446930 | 0.5352510 | 2408.0759 |
| Loxocephalus_sp | 5 | 4.65 | 4.04 | 231.7739 | 1889.1895 | 4.85 | 3.94 | 0.20 | -0.10 | 0.2236068 | 0.8944272 | -0.4472136 | -637.5663 |
| Loxocephalus_sp | 6 | 4.85 | 3.94 | 121.2058 | 1987.8819 | 1.07 | 6.88 | -3.78 | 2.94 | 4.7887368 | -0.7893522 | 0.6139406 | 1124.7674 |

Then we can calculate response diversity for an hypothetical community containing all the species tested i this experiment.

| time | E1_ref | E2_ref | Coleps_sp | Colpidium_striatum | Euplotes_daidaleos | Loxocephalus_sp | Paramecium_bursaria | Paramecium_caudatum | Stylonychia_sp | |
|---|---|---|---|---|---|---|---|---|---|---|
| 1 | 4.41 | 2.34 | -55.108269 | -223.002238 | -1.4385086 | -333.8104 | -407.08776 | -260.61916 | -575.88740 | 6. |
| 2 | 1.15 | 3.44 | 99.105500 | 345.818087 | 0.6358374 | 1432.1064 | 71.03804 | 87.73018 | 417.73480 | 5. |
| 3 | 2.91 | 3.08 | -18.328360 | 83.663318 | 0.5803385 | 545.9117 | -480.69137 | -356.04984 | -584.25370 | 6. |
| 4 | 0.61 | 1.48 | -2.881237 | 72.478259 | 1.0491311 | 2408.0759 | 89.69434 | 170.09649 | 364.83280 | 5. |
| 5 | 4.65 | 4.04 | 33.085863 | -2.674661 | 1.5920987 | -637.5663 | 332.75175 | -166.67661 | 172.95218 | 5. |
| 6 | 4.85 | 3.94 | -19.560641 | -14.883016 | -2.2036581 | 1124.7674 | -269.21168 | 258.81023 | -39.92325 | 5. |

Plot response diversity over time

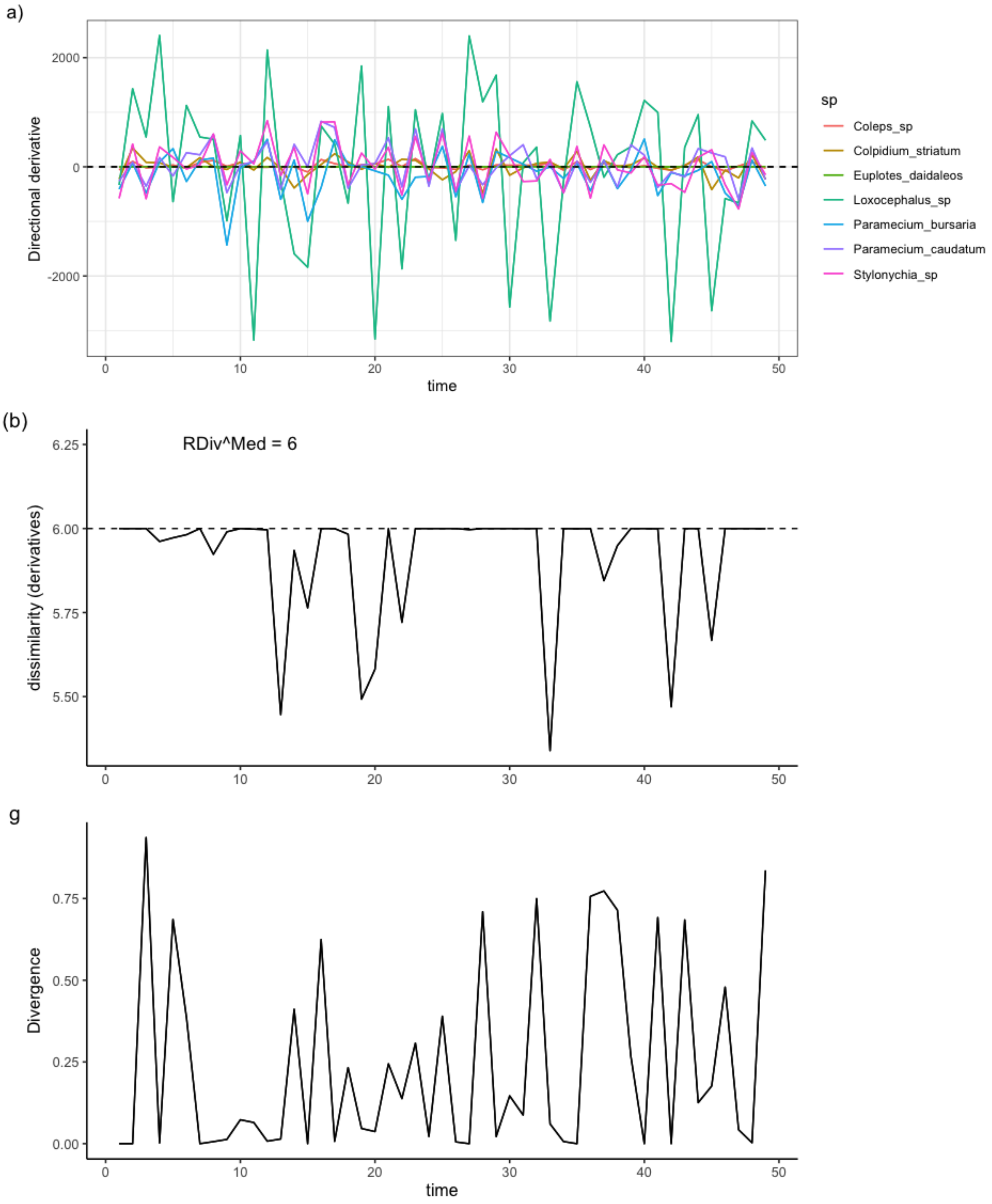


Figure 9.2: Directional derivatives and response diversity with known direction of env change. a. Species directional derivatives over time. b. Response diversity measured as similarity-based diversity metric. c. Response diversity measured as divergence (sign sensitive).

#### 9.2 Different community compositions

Now we calculate response diversity for three different community compositions we assembled randomly and we compare them.

Plotting

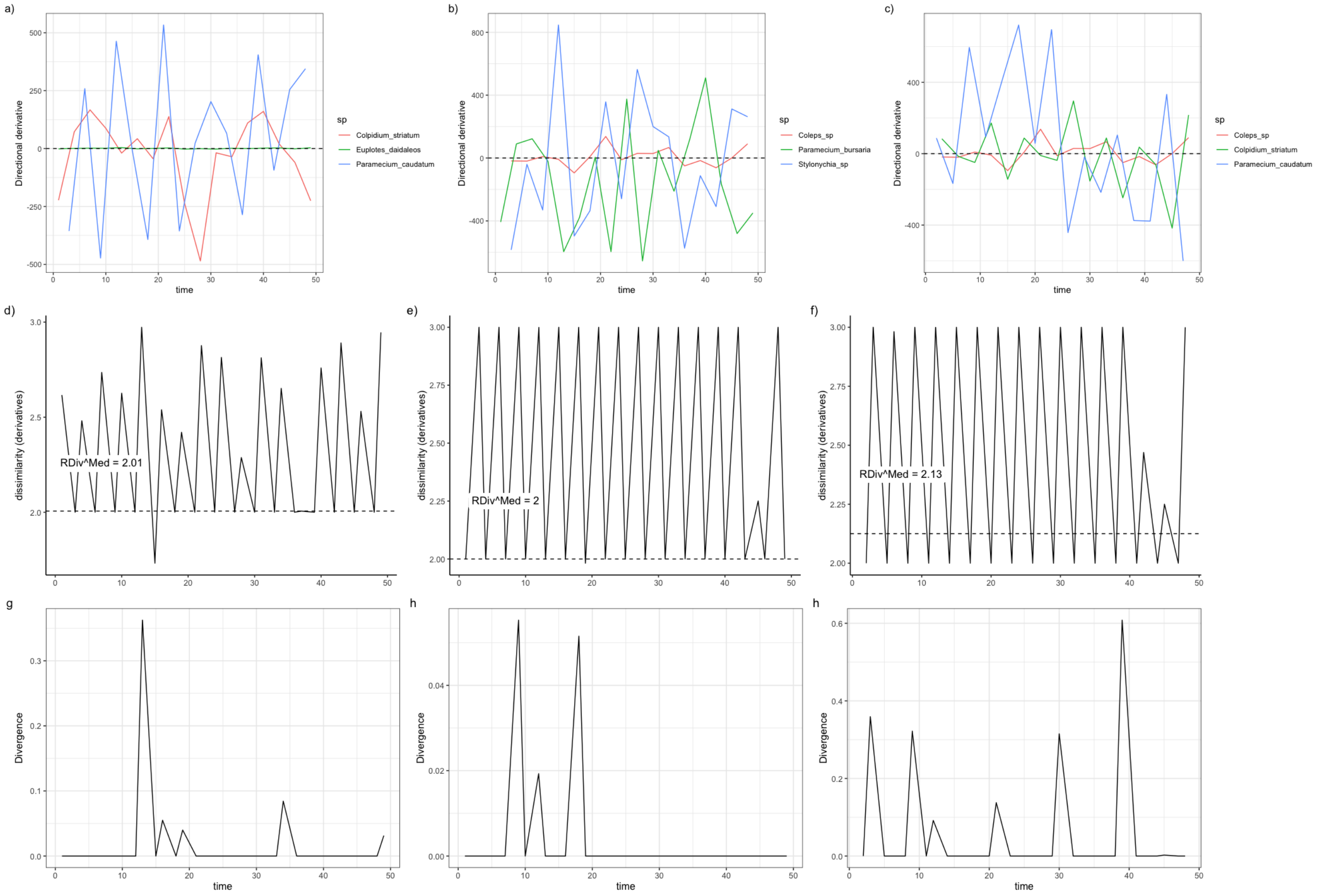


Figure 9.3: Directional derivatives and response diversity with known direction of env change for three different communities. a. Species directional derivatives over time. b. Response diversity measured as similarity-based diversity metric. c. Response diversity measured as divergence (sign sensitive).
